# *Escherichia coli* deletes *in vivo* the same domains from a double-mutant leucyl-tRNA synthetase gene that were deleted *in vitro* to make the LeuAC urzyme

**DOI:** 10.1101/2025.04.05.647346

**Authors:** Guo Qing Tang, Charles W. Carter

**Affiliations:** Department of Biochemistry and Biophysics, University of North Carolina at Chapel Hill, Chapel Hill, NC 27599-7260

**Keywords:** Origin of genetic coding, induced plasmid instability, single-turnover kinetics, aminoacylation, protein modularity, bidirectional coding

## Abstract

*E. coli* deletes CP1 and ABD domains from ∼40% of plasmids containing leucyl-tRNA synthetase if and only if both active-site signatures are mutated. Shortened ORFs occurred in all reading frames but form three discrete sets in the same frame. One had only the AVGA signature. Two longer ones both retained the same 24-residue segment containing the AMSAS signature. Large pre-steady-state bursts and steady-state acylation assays confirm that they encode active tRNA synthetases. In these respects, the ORFs resemble models for ancestral Class I aminoacyl-tRNA synthetases (AARS). Both signatures thus appear necessary and sufficient for aminoacylation. Growth at 4° C produced most of the middle-sized ORF, which is incompatible with the others and appears to result from a distinct mechanism. Three different linkages between the two parts of the active site acylate tRNA minihelix at similar rates. That result greatly expands the sequence space of active ancestral AARS. Widely spaced active-site mutants thus trigger deletions of modules acquired as full-length AARS evolved from simpler catalysts. We propose that the deletions survive because they limit mischarging due to disrupted coupling of active-site residues to domain motion. Such deletions may thus be a general phenomenon, opening broad access to primordial gene discovery.

**GRAPHICAL ABSTRACT:** Schematic rationale for formation and selection of LeuRS urzyme-like deletions. **A.** Full length LeuRS uses a complex allosteric network of interactions between Dom A (CP1), Dom B (ABD) and WT active-site catalytic histidine and lysine residues. **B.** Creation of the double mutant corrupts the allosteric effects of the two domains (faded colors). This creates a cytotoxic protein. **C.** Deletion of the domains whose functions have been corrupted by disrupting the allosteric network produces variants lacking the inactivated domains. These variants have considerably less cytotoxicity. They likely also resemble evolutionary precursors of the full-length protein. This may represent a general phenomenon for double mutant multi-domain protein genes.

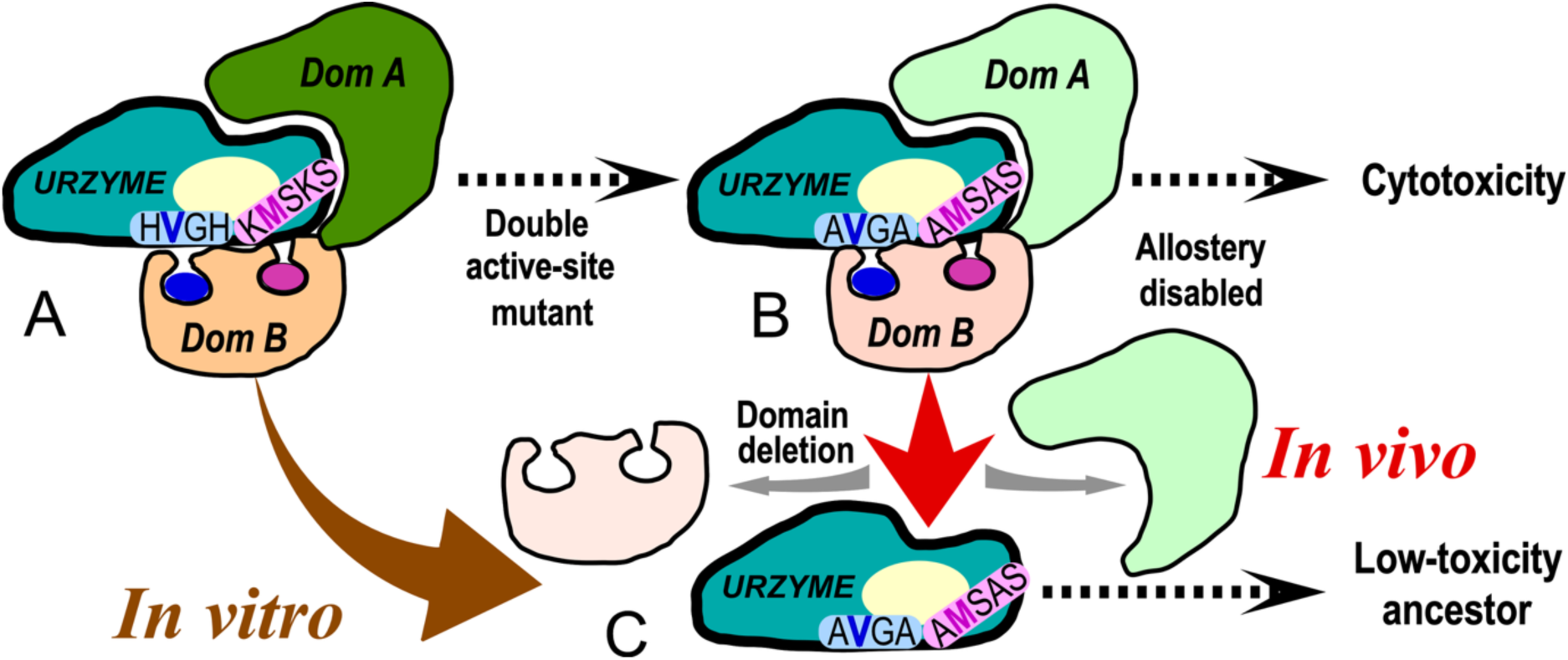

## INTRODUCTION

The Last Universal Common Ancestor, LUCA, lived about 4.2 billion years ago and had a nearly modern metabolism. Moody, et. al. proposed that LUCA had a nearly modern metabolism (<u>Moody, et al. 2024</u>). Similar estimates place the origins of organic chemistry on primitive earth to about the same time. Aminoacyl-tRNA synthetases (AARS) and their cognate tRNAs needed for translation appear to have acquired modern sequences before LUCA appeared (<u>Fournier, et al. 2011</u>; <u>Caetano-Anollés, et al. 2013</u>; <u>Fournier and Alm 2015</u>; <u>Wang, et al. 2025</u>). Thus, both nucleic acids (i.e., genes) and genetically-templated proteins must have achieved modern lengths within the errors of the various dates (<u>Ball 2025</u>). How did that happen?

To address that question we developed models of primordial AARS called “urzymes” (<u>Pham, et al. 2010</u>; <u>Li, et al. 2011</u>; <u>Carter 2014</u>) and “protozymes” (<u>Martinez-Rodriguez, et al. 2015</u>; <u>Onodera, et al. 2021</u>) to help define the transition leading to the code (<u>Carter and Wills 2021</u>; <u>Carter, et al. 2025</u>). Those models for very early stages of AARS evolution (<u>Li and Carter 2013</u>; <u>Weinreb, et al. 2014</u>; <u>Carter, et al. 2022</u>; <u>Tang, et al. 2023</u>; <u>Douglas, Bouckaert, et al. 2024</u>) are based on structural biology. “Urzymes” have 120-130 residues and nearly intact catalytic sites. “Protozymes” are ∼50-residue peptides containing only the ATP binding sites. The chief advantage of these experimental models is that their catalytic activities allow us still to study them experimentally. Some modern AARS, however, are nearly an order of magnitude longer than their presumptive primordial ancestors. That gap seems to have been filled by adding new, modular blocks of genetic information to the ancestral forms (<u>Douglas, Bouckaert, et al. 2024</u>).

A novel *in vivo* process in *Escherichia coli* may, at least in part, help to open a new window on how that happened. To better define the functional properties of AARS urzymes (<u>Tang, et al. 2023</u>), we had constructed a plasmid containing a mutant full-length *Pyrococcus horikoshii* leucyl-tRNA synthetase (LeuRS) gene in pET-11a (https://addgene.org/214193). The mutant gene had alanine mutations to both HVGH(AVGA) and KMSKS(AMSAS) signatures (<u>Tang, et al. 2023</u>*). E. coli* generated many discrete deletions from that exogenous, nicked plasmid output from high-fidelity PCR amplification. Curation of those deletions revealed sequences similar to those of the protozymes and urzymes we had made on purpose to mimic ancestral AARS.

It is routine to isolate and sequence plasmids from multiple transformants to identify one with the desired sequence (<u>Green and Sambrook 2012</u>). The polymerase chain reaction (<u>PCR; Mullis, et al. 1986</u>) can allow single point mutations to accrue. Mutant plasmids can also rearrange in cells grown after transformation. Heretofore, such variation has generally been thought to be a nuisance, as for example were the earliest observations of microRNAs in gels and northern blots (<u>Varani 2015</u>). This is especially true in cases such as ours, that may involve the orthogonality of the respective AARS (<u>Anderson and Schultz 2003</u>). Various precautions can increase the frequency of the desired plasmid (<u>Green and Sambrook 2012</u>) (see also (<u>Tomoiaga, et al. 2021</u>)).

To our knowledge, nobody has considered abundance profiling of plasmids produced after transformation as a coherent source of insight about the evolution of protein modularity. In our case, however, the surviving plasmids contained uniquely useful data. These data suggest several novel and functional configurations of the modules we previously showed to be models for ancestral AARS. We outline reasons to think this *in vivo* system may be generally useful in the discovery of primordial genes and thus offer an important new window on the modular assembly of protein genes.

## MATERIALS AND METHODS

### Materials

Genes for Clones 21 and 22 were inserted into pMal-c2x (Plasmid #75286. New England Biolabs, Ipswich, MA.). DNA oligos were ordered from IDT (Integrated DNA Technologies, Coralville, IA, United States). The detailed sequences of oligo DNAs used for mutagenesis (*a1/b1*) and for sequencing (*seqn1/seqn2*) are provided in the Table 2, together with the sequence of full-length LeuRS gene. Phusion™ Plus PCR Master Mix (Cat# F631S) was purchased from Thermo Fisher Scientific (Waltham, MA, United States). *E coli* competent cells were partially from Agilent (XL10-Gold ultracompetent cells, Cat# 200315, Santa Clara, CA, United States), and partially using DH5a, home-made, prepared according to the method described by Sharma in 2017. Restrict enzyme DpnI (Cat# 500402, United States) was purchased from Agilent. Purifications and handling of DNA fragment and plasmid used the QIAquick Gel Extraction Kit (Cat# 28704, Qiagen, Hilden, Germany) and QIAprep Spin Miniprep Kit (Cat# 27104, Qiagen, Hilden, Germany) according to instruction, unless stated otherwise.

### Methods

#### Method for preparation of double mutant plasmid

DNA oligo sequences were designed according to the manual of QuikChange II-E Site-Directed Mutagenesis Kit (Santa Clara, CA, United States) as published online at https://www.agilent.com/cs/library/usermanuals/public/200555.pdf. In order to create the double mutant plasmid DNA, we have implemented a megaprimer PCR method based on that described by Picard (<u>Picard, et al. 1994</u>) (Fig. 1.).

**Figure 1.**
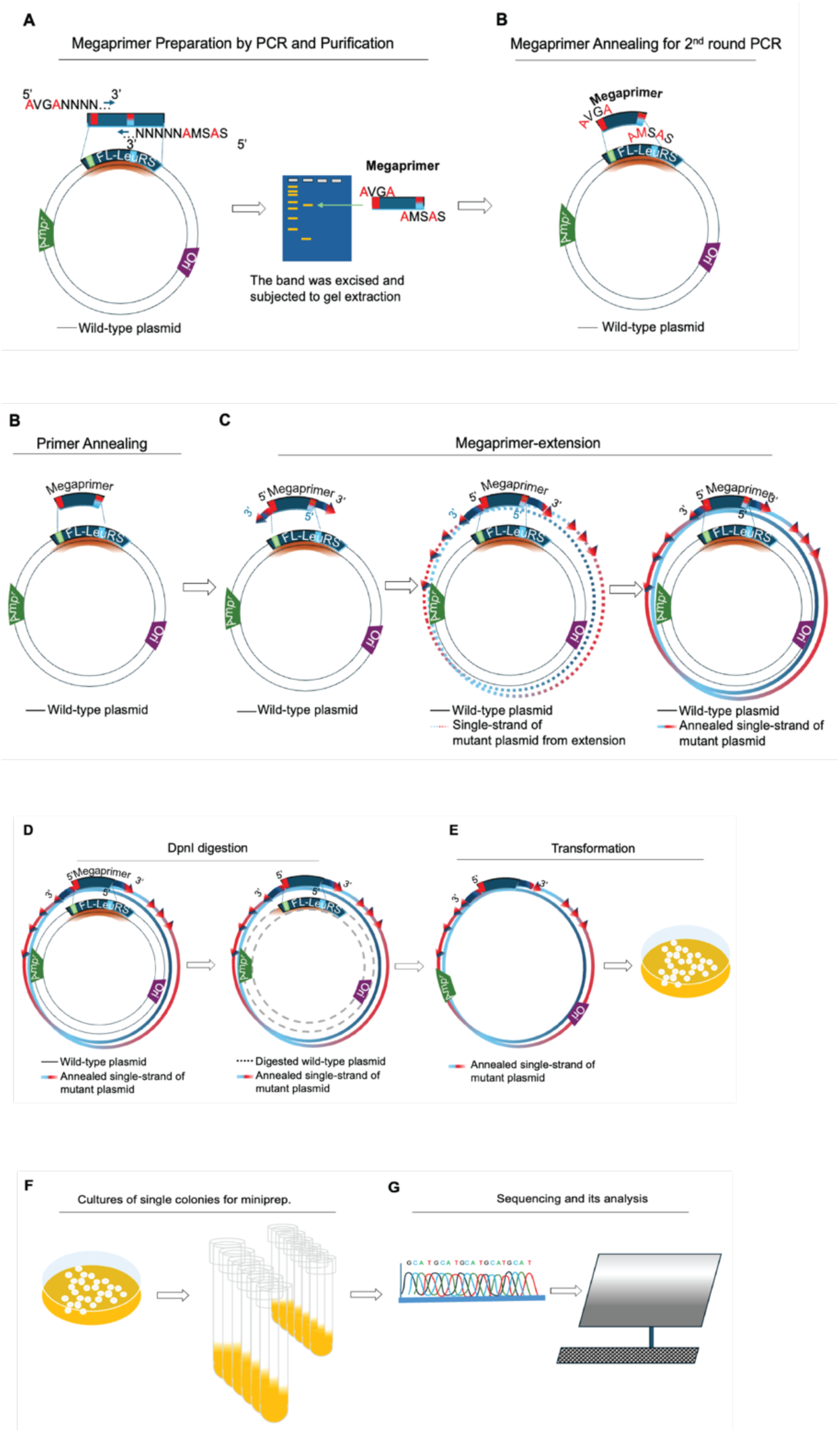
A flowchart illustration of megaprimer-PCR. The method employs a combination of the "Megaprimer-PCR" technique (<u>Picard, et al. 1994</u>) with conventional mutagenesis products (e.g., Agilent, Santa Clara, CA (<u>Technologies</u>)). **A**. Following a first round of PCR (left) with releasing a megaprimer from a target wild-type plasmid DNA, an agarose gel and the PCR product (namely, megaprimer) purification was carried out, in which the propriate mutations AVGA and AMSAS were installed as indicated in bended color squares and amino acid sequences. **B.** For the second round of PCR, annealing the newly purified and resultant megaprimer onto a target plasmid DNA. **C.** PCR extension of the second round of PCR is indicated. The desired PCR extension is initiated and proceeded using the sense-single strand and antisense-single strand of the megaprimer as indicated as colored dot circular lines. **D.** Restriction enzyme DpnI digestion of the second round PCR product, the destroy of original intact wild-type plasmid DNA is indicated as dash lines in grey. **E.** After DpnI digestion of the second round PCR mixture, single colonies of transformants on agar plates can be obtained via conventional transformation with DpnI-digested PCR mixture into competent *E. coli* cells. **F.** Following transformation, a variety of mutant plasmids can be identified through conventional plasmid miniprep. **E.** sequencing, sequence analysis and its identification.

First, megaprimer was prepared in a 1^st^ round of PCR. Phusion™ Plus PCR Master Mix (Cat# F631S) was employed according to instruction from manufacture (ThermoFisher Scientific, Waltham, MA, United States). The primer pairs for the 1^st^ round PCR was LeuRS_AVGAa1_F and LeuRS=AMSASb1_R as listed in Table 1 above. In the PCR reaction mix, the plasmid template concentration was 1-4 ng/mL. The primers were used at a concentration of 0.66 mM. The PCR was started at 98 C, 30 sec for initial template denaturation and was following by 44 cycles (98 °C, 12 sec; 61 °C, 30 sec; 72 °C, 3 min 20 sec). After the final extension at 72 °C, 5 min, the PCR reactions were kept at 4 °C overnight. After the PCR run, following by horizontal electrophoresis in a 1xTAE, 1.2% Agarose-gel, and visualization by UV transmitter, the megaprimer fragment from the 1^st^ round of PCR was collected from gel cutting out the desired band using a clean razor blade. After DNA fragment purification procedures using QIAquick Gel Extraction Kit (Cat# 28704, Hilden, Germany), to yield the megaprimer for the downstream operations.

**Table 1.**
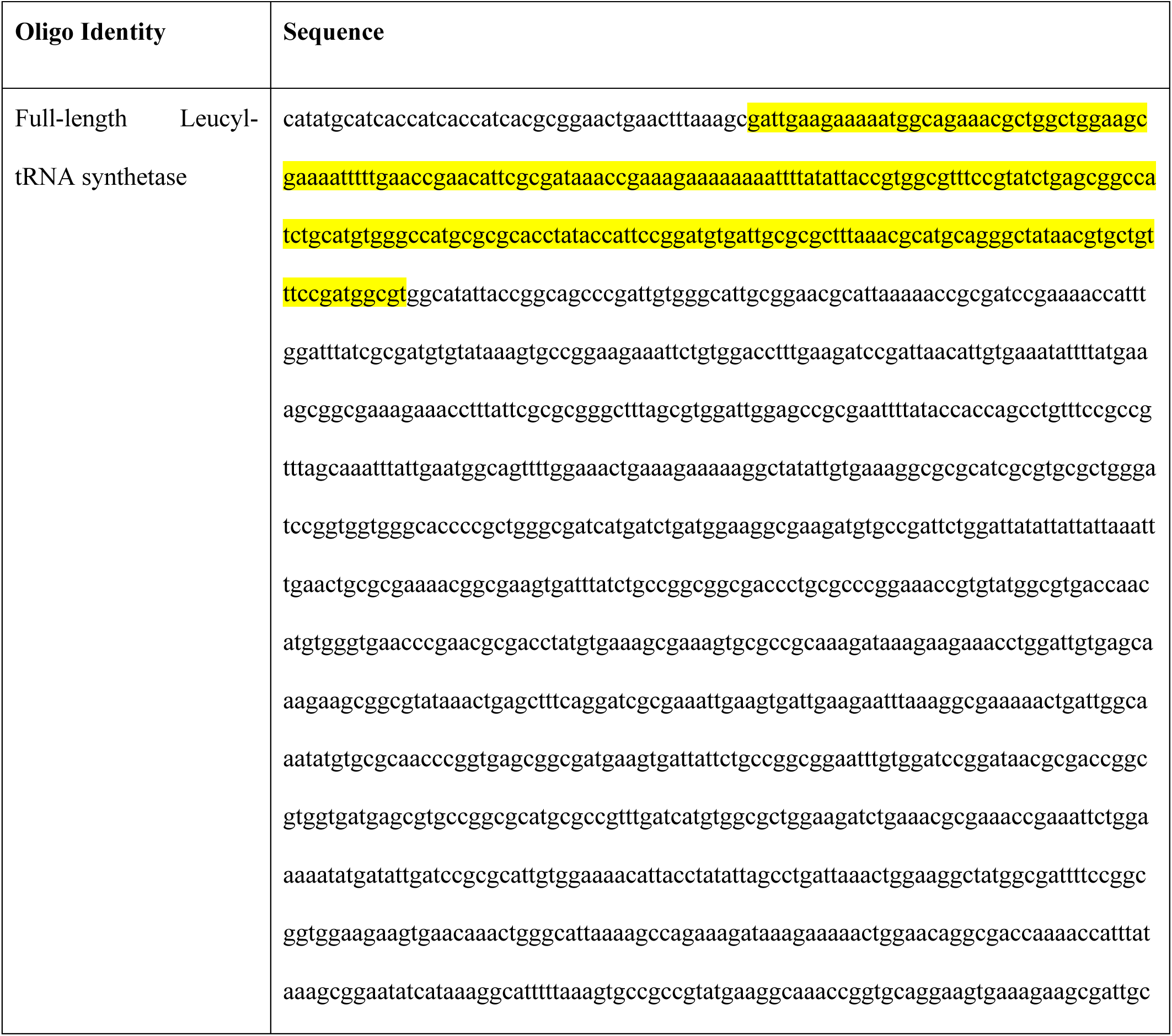
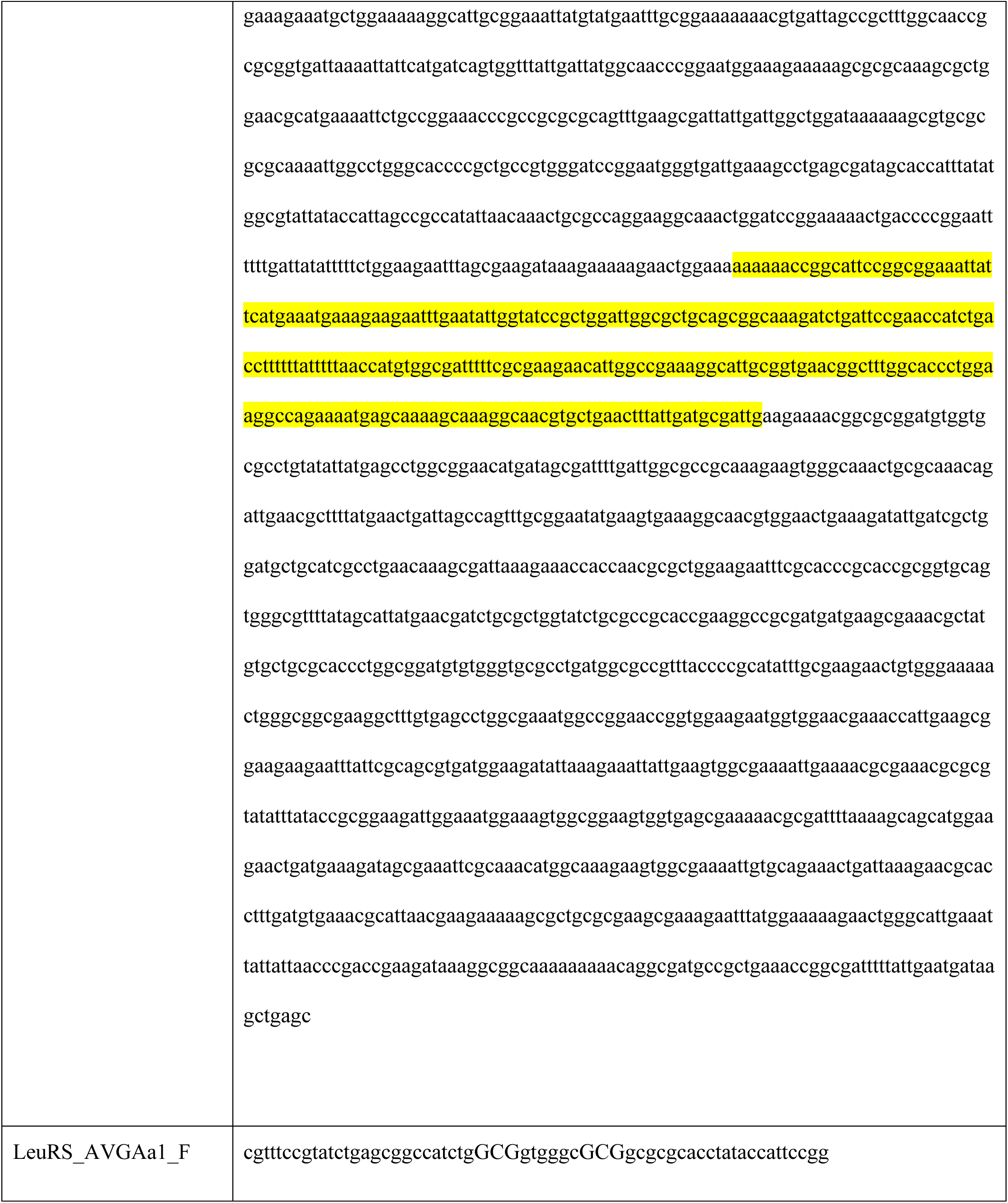
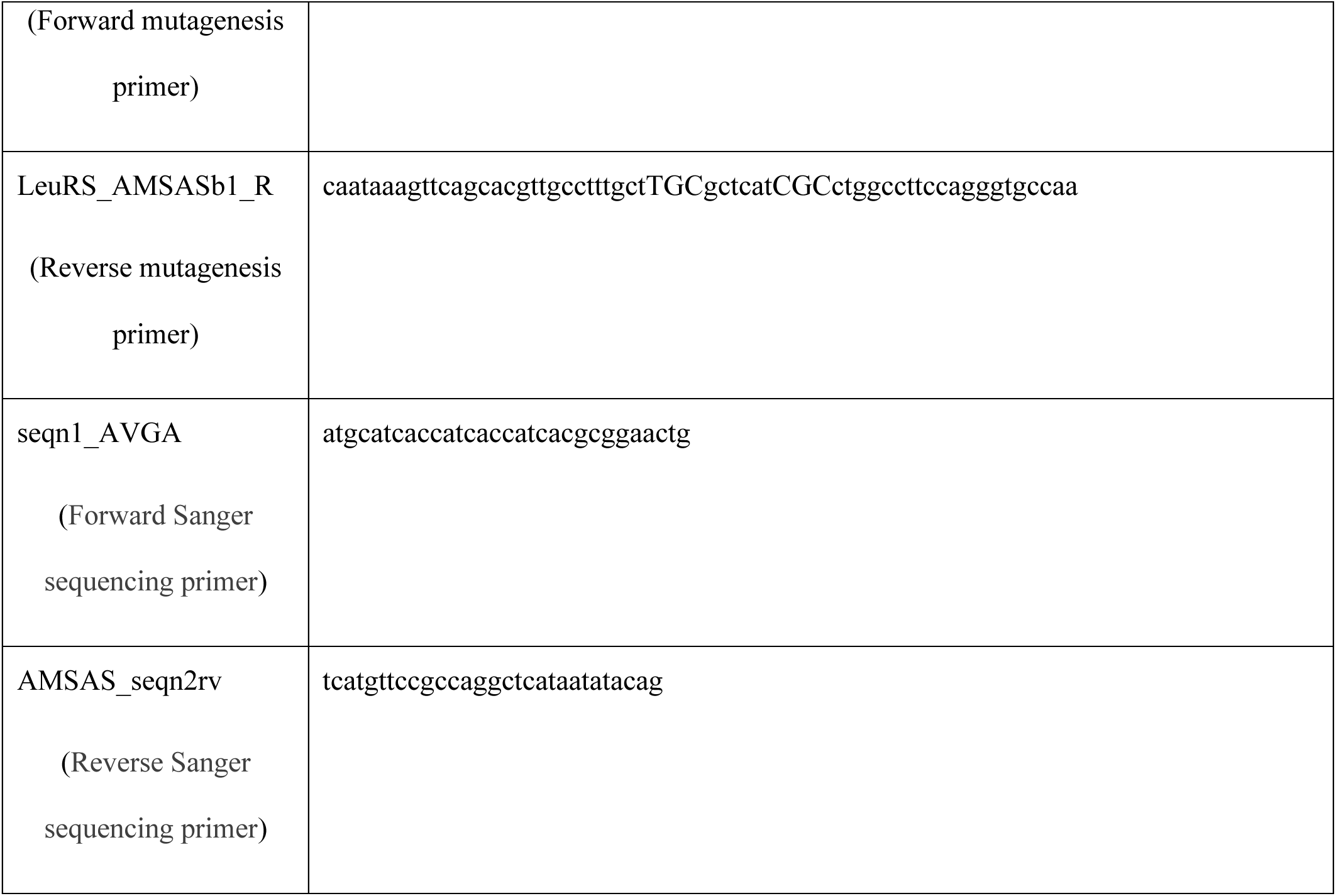
Summary of DNA template sequences of *P. horikoshii* Ph-FL-LeuRS and Oligos for the study. Sequences encoding WT LeuAC urzyme are highlighted in yellow.

In the second phase, the entire plasmid DNA was amplified using the megaprimer to generate the resultant plasmid harboring the mutant. A 2^nd^ round of PCR was performed using Phusion™ Plus PCR Master Mix (Cat#F631S) and the parameters were setup according to the instructions from the manufacture (ThermoFisher Scientific, Waltham, MA). The PCR procedures were the same as the 1^st^ PCR run. Very importantly, instead of oligo primers, the megaprimer was added at specific desired concentration. Basically, in a 25-40 mL PCR reaction volume, the amount of megaprimer fragment was within a range of 0.16mM-1.25 mM, and the template plasmid DNA was 0.5-1.5 ng.

After DpnI digestion of the PCR mixture at 37°C from hours to overnight according to instruction provided by Agilent (Santa Clara, CA, United States), the DpnI-digested PCR mixture is ready for *E. coli.* transformation. The transformation was done using 0.3 mL DpnI-digested PCR mixture per transformation.

#### Method for E. coli. transformation and the sample submission for sequencing

*E coli* transformation was proceeded according to conventional procedure (<u>Fritsch, et al. 1982</u>) and spread onto LB (Lysogeny broth) agar plates with antibiotics (Ampicillin 50 mg/mL, or Carbenicillin 100 mg/mL). After incubation overnight at 37 °C, individual colonies on the LB agar plate were readied for miniprep by following the standard protocol described by QIAprep Spin Miniprep Kit (Cat# 27104, Qiagen, Hilden, Germany)

The resultant plasmid DNA samples were subjected to sequencing by following standard sequencing sample preparation and submission described by Eton Bioscience (San Diego, CA) online https://www.genewiz.com/en/Public/Resources/Sample-Submission-Guidelines/Sanger-Sequencing-Sample-Submission-Guidelines/Sample-Preparation#sanger-sequence with minor modifications. Basically, each sequencing sample was in 15 mL, containing 100-200 ng plasmid DNA with 5-10 picomole sequencing primer, i.e. seqn1_AVGA or AMSAS_seqn2rv as detailed sequences in Table 1 above.

#### Method for plasmid sequencing and sequencing analysis

Sequence analysis was performed using conventional methods as follows. First, all the DNA sequences were translated into 6 reading frames in amino acid sequence using Expasy translate online at https://web.expasy.org/translate/. For screening the tentative urzyme candidates, each amino acid sequence from 6 reading frames from a single sequenced plasmid DNA sample were aligned with the sequence of urzyme template using conventional sequencing alignment software such as MAFFT online at https://www.ebi.ac.uk/jdispatcher/msa/mafft?stype=protein (<u>Katoh, et al. 2019</u>) or MUSCLE online at https://www.ebi.ac.uk/jdispatcher/msa/muscle?stype=protein(<u>Edgar 2004</u>). The list of criteria for selection of tentative candidate was according to the sequence length, similarity and identify in addition to the highly conserved catalytic motifs such as motif I HVGH and motif II KMSKS or their-like.

The resulting new candidates were designated protozyme, urzyme-like or Δ2ndXvr (for deleted second crossover) and goldilocks, according to their sequence content and were collected and subjected for further downstream identification at the enzymology level after protein biochemistry procedures.

#### tRNA^Leu^ and TΨC-minihelix^Leu^ preparation

A plasmid encoding the *P. horikoshii* tRNA^Leu^ (UAG anticodon) was synthesized by Integrated DNA Technologies and used as template for PCR amplification of the tRNA and upstream T7 promoter and downstream Hepatitis Delta Virus (HDV) ribozyme. The PCR product was used directly as template for T7 transcription. Following a 4-hour transcription at 37°C the RNA was cycled five times (90°C for 1 min, 60°C for 2 min, 25°C for 2 min) to increase the cleavage by HDV. The tRNA was purified by urea PAGE and crush and soak extraction. The tRNA 2’-3’ cyclic phosphate was removed by treatment with T4 PNK (New England Biolabs) following the manufacturer’s protocol. The tRNA was then phenol chloroform isoamyl alcohol extracted, filter concentrated, aliquoted, and stored at -20°C.

The original sequence information for composing and designing *Pyrococcus horikoshii* TΨC-minihelix-Leu was based on the mature *P. horikoshii* tRNA-Leu, the tRNA sequence was collected from the complement strand of the genomic sequence of *P. horikoshii* (OT3, GB# BA000001.2, between nucleotides 1448081 and 1448168). The designated minihelix sequence was obtained by combining the acceptor stem sequence including DCCA with the TΨC -stem-loop according to Schimmel (<u>Schimmel and Alexander 1998</u>). A plasmid harboring the minihelix, an upstream T7 promoter, and downstream Hepatitis Delta Virus (HDV) ribozyme was acquired from previous co-worker Jessica Elder. To reduce challenges posed by the high GC content in the stem portion of minihelix during the preparation, we implemented the Phi29 DNA polymerase-mediated isothermal amplification approach according to the manufacture’s protocol (New England Biolabs, Ipswich, MA). After the purification of the product from isothermal amplification, T7 transcription was carried out. Following a 4–6-hour transcription at 37 ^◦^C, the reaction mixture was subjected to urea PAGE fractionation, crush, and soak extraction. After 2’-3’ cyclic phosphate removal by T4 PNK, the minihelix RNA was then phenol chloroform isoamyl alcohol extracted, filter-concentrated, quantitated, and aliquoted, and stored at −80 ^◦^C.

#### Expression and purification of deletion variants

We expressed Clones 21 and 22 as MBP fusions from pMAL-c2x in BL21Star (DE3) (Invitrogen) as described in (<u>Tang, et al. 2024</u>). The lysis buffer contained 20 mM Tris, pH 7.4, 1 mM EDTA, 5 mM β-mercatoethanol (βME), 17.5% Glycerol, 0.1% NP40, 33 mM (NH_4_)_2_SO_4_, 1.25% Glycine, 300 mM Guanidine Hydrochloride) plus cOmplete protease inhibitor (Roche). After elution from Amylose FF resin (Cytiva) with 10 mM maltose in lysis buffer (200 mM HEPES, pH 7.4, 450 mM NaCl, 100 mM KCl, 10 mM β-ME. Fractions containing protein were concentrated and mixed to 50% glycerol and stored at -20°C. All protein concentrations were determined using the Pierce™ Detergent-Compatible Bradford Assay Kit (Thermo Scientific). We determined purity by running samples on PROTEAN® TGX (Bio-RAD) gels and active fractions for all variants were measured as described in the next section.

#### Single turnover active-site titration assays

Active-site titration assays were performed as described (<u>Fersht, et al. 1975</u>; <u>Francklyn, et al. 2008</u>) with the exception that ^32^P-ATP was labeled in the α position in order to follow the time-dependence of all three adenine nucleotides. of protein was added to 3 μM in 1x reaction mix (50 mM HEPES, pH 7.5, 10 mM MgCl_2_, 5 μM ATP, 50 mM amino acid, 1 mM DTT, inorganic pyrophosphatase, and α-labeled [^32^P] ATP) to start the reaction. Timepoints were quenched in 0.4 M sodium acetate 0.1% SDS and kept on ice until all points had been collected. Quenched samples were spotted on TLC plates, developed in 850 mM Tris, pH 8.0, dried and then exposed for varying amounts of time to a phosphor image screen and visualized with a Typhoon Scanner (Cytiva). The time-dependent of loss (ATP) or de novo appearance (ADP, AMP) of the three adenine nucleotide phosphates were estimated using the ImageJ measure function. Time-dependence was fitted using the nonlinear regression module of JMP16PRO™ to equation (1):

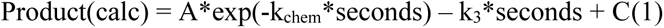

where k_chem_ is the first-order rate constant, k_3_ is the rate of turnover, A is the amplitude of the first-order process, and C is an offset.

For [ATP] decay curves, the fitted A value estimates the burst size, n, directly as n = A*[ATP]/[Enzyme]. C gives n = (1–C) * [ATP]/[Enzyme]. For exponentially increasing concentrations of product, the situation reverses, and n = (1–A) * [ATP]/[Enzyme]and n = C*[ATP]/[Enzyme]. These approximations are justified by the very small variance of multiple estimates.

#### Aminoacylation assays

We determined the active fraction of minihelix RNA by following extended acylation assays using the AVGA LeuAC mutant until they reached a plateau. That plateau value was used to compute minihelixconcentrations in assays with all variants.

Aminoacylations withs minihelix were performed as described (<u>Tang, et al. 2024</u>) in 50 mM HEPES, pH 7.5, 10 mM MgCl_2_, 20 mM KCl, 5 mM DTT with indicated amounts of ATP and isoleucine. The high affinity of Clones 21 and 22 for minihelix meant that we had to mix increased amounts of [α^32^P] A76-labeled tRNA for assays. The RNA substrates were heated in 30 mM HEPES, pH 7.5, 30 mM KCl to 90°C for 2 minutes and then cooled linearly (drop 1°C/30 seconds) until it reached 80°C when MgCl_2_ was added to a final concentration of 10 mM. The tRNA continued to cool linearly until it reached 20°C. Michaelis-Menten experiments were performed by repeating these assays at the indicated RNA concentrations. Concentration-dependence fitted to K_sp_ = k_cat_/K_M_ and kcat according to the modified formula (Eqn. 2) introduced by Johnson (<u>Johnson 2019</u>).

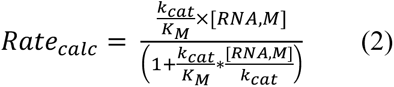

#### Data processing and Statistical analysis

Phosphor imaging screens of TLC plates were densitometered using ImageJ. Data were transferred to JMP16PRO™ Pro 16 via Microsoft Excel (version 16.49), after intermediate calculations. We fitted both active-site titration curves and Michaelis-Menten assays using the JMP16PRO™ nonlinear fitting module.

Factorial design matrices (e.g. Table 1) were processed using the Fit Model multiple regression analysis module of JMP16PRO™ Pro, using an appropriate form of equation (3) (<u>Box, et al. 1978</u>)ΔG‡

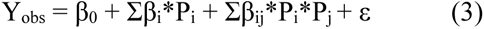

where Y_obs_ is a dependent variable, usually an experimental observation, β_0_ is a constant derived from the average value of Y_obs_, β_i_ and β_ij_ are coefficients to be fitted, P_i,j_ are independent predictor variables from the design matrix, and ε is a residual to be minimized. All rates and apparent affinities were converted to free energies of activation or binding, ΔG = – RTln(k), before regression analysis. Free energies are additive, whereas rates and apparent affinities are multiplicative. For example, the activation free energy for the first-order decay rate in single-turnover experiments is ΔG‡(kc_hem_).

Multiple regression analyses of factorial designs exploit the replication inherent in the full collection of experiments to estimate experimental variances. In most cases we estimated errors on the basis of t-test P-values. In some cases, we added error bars to histograms to show the variance of individual datapoints. Multiple regression analyses reported here also entail triplet experimental replicates, which enhance the associated analysis of variance.

## RESULTS

We describe a novel fragmentation of the leucyl-tRNA synthetase gene from *P. horikoshii* that occurs in ∼40% of the plasmids recovered from *E. coli* cells transformed with a double active-site mutant LeuRS plasmid. We described this plasmid as part of an earlier mutagenesis experiment (<u>Tang, et al. 2023</u>). There is nothing subtle about what we did. It is a standard mutagenesis experiment. No selection was applied other than antibiotic resistance. The *E.* coli strain lacked the RecA recombinase. To preclude adventitious loss of sequences during the PCR the double mutant full-length sequence served as a megaprimer (<u>Picard, et al. 1994</u>), as described in Methods. The expression vector, pET-11a, greatly reduces the leaky synthesis of mRNA. Single colonies were sampled without inducing protein expression and grown for sequencing. We had no reason to expect any unusual results.

However, the plasmid abundance profile had highly unusual aspects. To understand these, we summarize the LeuRS structural biology and discrete products formed after transformation, in Fig. 2.

**Figure 2.**
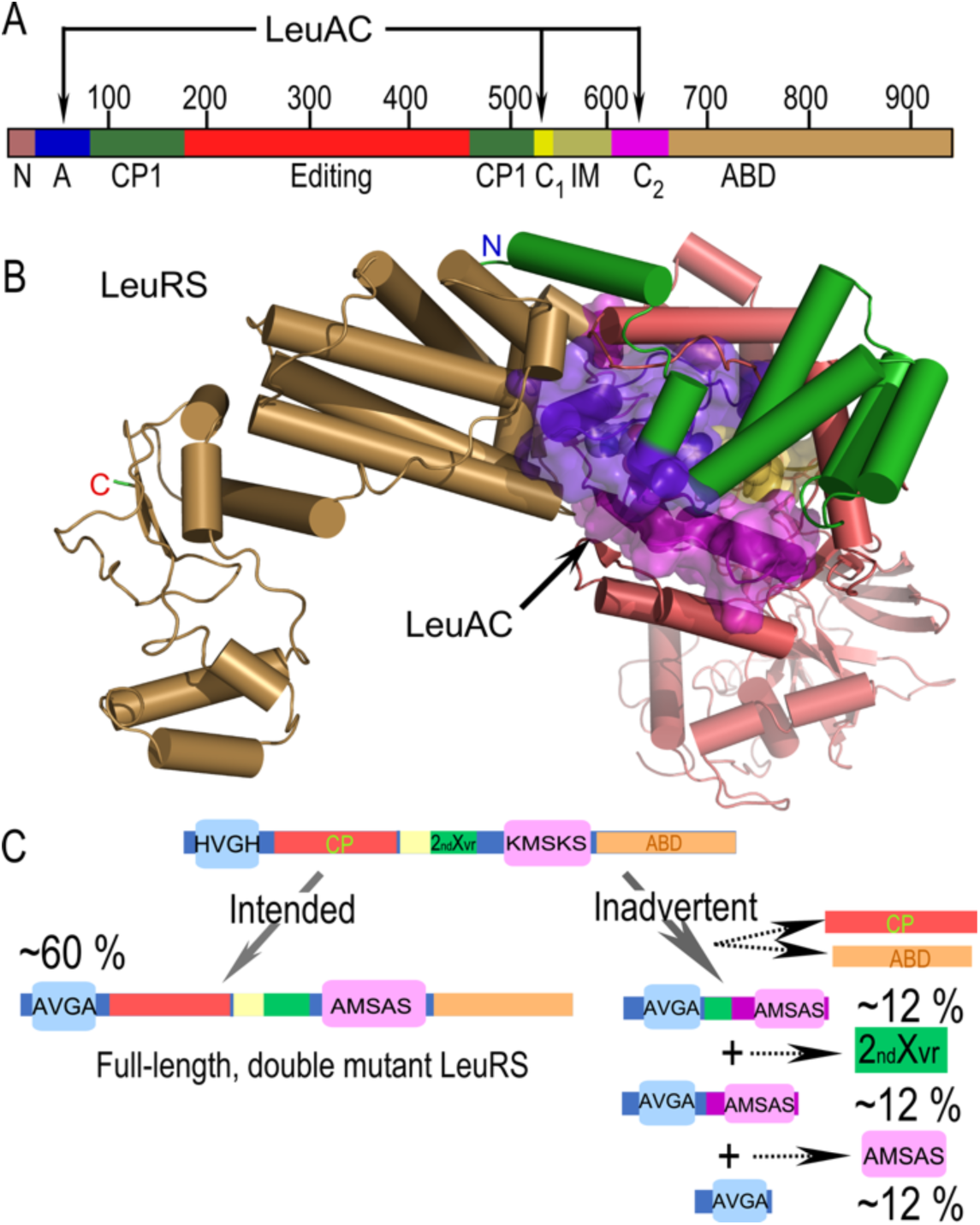
Reference structural schematic of the experiments described herein. **A**. Linear schematic of the 3D structure of the mosaic structure of *P. horikoshii* LeuRS and the LeuAC urzyme. CP1 denotes connecting peptide 1 as described (<u>Burbaum, et al. 1990</u>; <u>Burbaum and Schimmel 1991</u>). Editing denotes the editing domain embedded in CP1. ABD denotes the anticodon-binding domain. A, C1 and C2 are discontinuous segments of LeuRS that form LeuAC. They each are joined by single peptide bonds following excision of the intervening sequences to make a single polypeptide. **B.** 3D cartoon of LeuRS, colored as in A. The LeuAC urzyme we study as a model for the early ancestor of LeuRS is shown by the blue and magenta surface. The components of LeuAC are also indicated above **A. C.** Comparison of the intended double mutant and the three subsets of short, fragmentary ORFs formed *in vivo* from the starting full-length double mutant gene. Note that the 2^nd^ Crossover connection of the Rossmann fold (2ndXvr) is colored green here for clarity. The different N-terminal (blue) segments in truncated fragments are not to scale.

### *E. coli* creates a discrete population of LeuAC urzyme-like coding regions *in vivo*

Here, we describe the abundance profiling of the *in vivo* deletions (Fig. 2C, Fig. 3) we observed as part of the previous work (<u>Tang, et al. 2023</u>). We sequenced 180 plasmids from transformed single colonies to assess the degree of homogeneity in the ensemble. About 60% (108) of sequenced plasmids isolated had the desired, full-length double mutant. The remaining ∼40% had long deletions of coding sequences. The deletion plasmids all contained short stretches of LeuRS coding sequences. Rather than ∼2900 nt, they have between 290 – 450nt, far shorter than the full length LeuRS gene. These were re-sorted in idiosyncratic ways. Coding sequences occurred in all six reading frames. Yet, when annotated for continuous open reading frames and aligned to that of the full length LeuRS gene, they form the coherent alignment shown in Fig. 3A. The raw sequence data for the insertions in 34 of these plasmids from Fig. 3A and 3C are in Supplemental Section A. A fasta file with these and other, similar deletion plasmids is in Supplemental Section B. Genomic features, together with the temperature at which they were incubated and the respective reading frames are shown schematically in Supplementary Figs. S1-S7 and collected in Supplementary Table S1.

**Figure 3.**
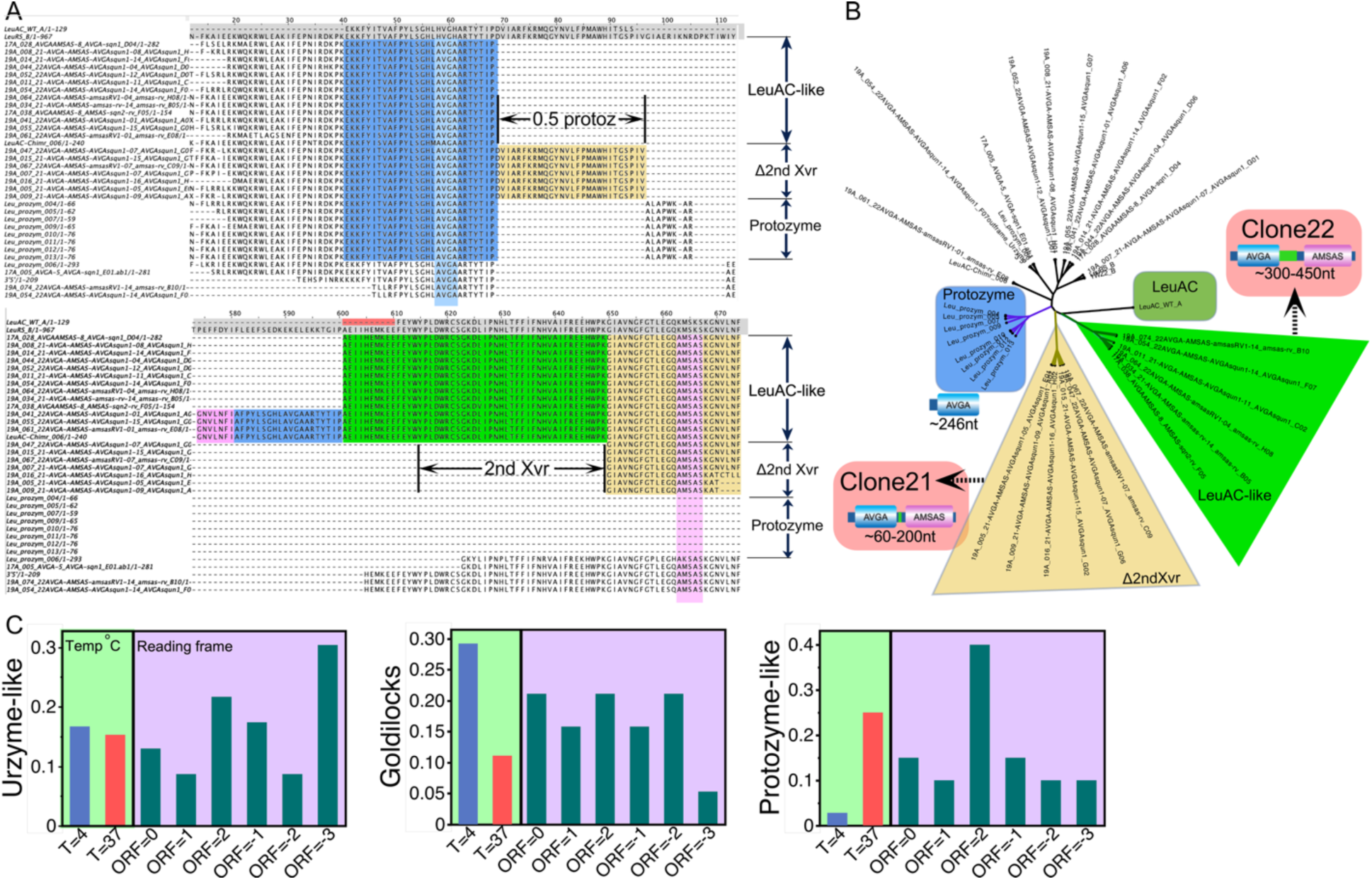
Shortened ORFs generated *in vivo* from full-length LeuRS double active-site mutants. **A.** Modular deletions and shuffling in excerpts from the multiple sequence alignment (<u>Tang and Carter 2025</u>). Both panels together show the LeuAC sequence and corresponding segments from full-length *P. horikoshii* leucyl-tRNA synthetase in rows 1 and 2 (light grey background at the top). Vertical light blue and magenta columns indicate the two mutant signatures. Green horizontal backgrounds highlight uniquely retained sequences of the LeuAC-like urzyme, Clone 22. These lack the second half of the protozyme (blue background). Sand backgrounds highlight urzyme sequences uniquely retained in the Goldilocks variant, Clone 21. It lacks the second crossover connection of the Rossman dinucleotide-binding fold (<u>Buehner, et al. 1973</u>; <u>Burbaum and Schimmel 1991</u>). **B.** Neighbor-joining tree from the alignment in A highlights the three principal clades of related sequences. Coloring is that in A. For subsequent reference Clone21, the Goldilocks deletion variant, was excerpted from one of the second crossover deletions (denoted Δ2nd Xvr). Clone 22 contains the second crossover lacking in Clone21 and has an extended segment preceding it (red band under sequence numbers). Clones 21 cannot be a subset of Clones 22.**C.** Genomic properties show that incubating transformant colonies at lower temperatures leads to the appearance of Goldilocks deletions (green panels). Magenta panels show the distribution of reading frame as follows. ORF 1 is the same reading frame as the intact LeuRS plasmid. ORFs 1,2 are advanced by 1 and 2 frames. ORFs -1, -2, -3 are corresponding frames in the 3’-5’ direction. The different temperature-dependence and mutually exclusive sequences in Goldilocks and Urzyme-like coding regions imply that they arise via different intracellular DNA rearrangements. Further details on the incidence of tandem repetition are included in Supplementary Table S1.

We describe here only the first 34 such sequences analyzed. ORFs from all the additional, subsequently sequenced plasmids fall into the same three categories as those highlighted by black vertical arrows in Fig. 3A.

Each clade differs in a distinct way from the analytical LeuAC urzyme (<u>Carter, et al. 2014</u>; <u>Hobson, et al. 2022</u>) we studied earlier (<u>Tang, et al. 2023</u>; <u>Tang, et al. 2024</u>). Fig. 3A summarizes this unexpectedly coherent mosaic. A neighbor joining tree based on the alignment in Fig. 3A reveals only three clades (Fig. 3B). All shortened ORFs deleted major segments from the full-length LeuRS gene. Notably, there are no remnants of the two large domains we deleted to generate LeuAC. The first missing domain is the 444-residue Connecting Peptide 1 (CP1; residues 83-527) that provides the editing function. The second domain missing in all deletions is the 304-residue anticodon-binding domain (ABD; residues 663-967).

The coding sequences retained in the shortened ORFs comprise three discrete sets, each with different properties. The shortest subset (Supplemental Figs S2-S4) retains only a 28-residue segment (residues 41-68) containing the AVGA sequence and which forms the ATP binding site. About 11% (8) of the 34 sequences encode truncated protozymes, (blue backgrounds) retain only these sequences. We refer to this set as protozyme-like (from “Proto”, relating to a precursor). These sequences are highlighted in blue in Fig. 1A. These residues are present in virtually all deletions.

A longer subset (Supplemental Figs. S1,4) contains, in addition, to the intact, 50-residue protozyme, residues 650-674 that contain the AMSAS motif. This 25-residue segment is against a sand background in Fig. 3A. Although it contains the entire length of the protozyme, it is the shorter of the two deletions that contain both mutant catalytic signature motifs. About ∼10% (7) of the sequences in Fig. 3A have this intermediate length. It is ∼81 residues long. We expressed this ORF as Clone 21 and for reasons outlined in detail below, we refer to it as the Goldilocks urzyme because it is just the right length to align antiparallel to minimal Class II urzymes.

The longest ORF in Fig. 3A (300 – 450nt; 19%; 14 sequences; Supplemental Figs. 1, 6,7), includes residues 588-636 which code for the second crossover (<u>Richardson 1973</u>) of the Rossmann fold (<u>Buehner, et al. 1973</u>) (green backgrounds in Fig. 3A). That segment contributes a third motif, **G**K**D**L (**G**x**D**Q in other Class I AARS). It contributes the aspartate side chain to the active site and we presume it coordinates the active-site Mg^++^ ion (<u>Weinreb and Carter 2008</u>; <u>Weinreb, et al. 2009</u>; <u>Williams, et al. 2015</u>). These sequences also have only a half-protozyme and an extra segment that is not part of LeuAC (red band just under the sequence numbers). We expressed this ORF as Clone 22 and refer to it as “urzyme-like”.

Curiously, the lengths of the truncated protozyme ORFs match exactly that portion of the protozyme in the LeuAC-like ORFs. The protozyme segment is twice as long in the intermediate-sized ORF which lacks the entire second crossover segment of the Rossmann fold (<u>Buehner, et al. 1973</u>). Each clade in the ensemble is remarkably discrete and reproducible. Occasional point mutations outside the LeuRS coding regions that differentiate the different truncated plasmids from one another are not highlighted.

We call the 81-residue ORF the “Goldilocks” urzyme because it is “just right” for antiparallel alignment with the shortest Class II urzymes (<u>Patra, et al. 2024</u>). We return to the Goldilocks urzyme in a subsequent section, because of its inherent compatibility with the shortest Class II AARS urzymes.

Supplementary Section C contains figures showing the variety of urzyme, Goldilocks, and protozyme sequences that appeared in sequenced plasmid samples as summarized in Fig. 3C. Curious genomic features include single open-reading frame deletions, (Supplemental Figure S1, S5); tandem-configurations, (Supplemental Figures S2, S3, S7); and multi-frame embedding within the same plasmid (Supplemental Figures S3, S4, S6, S7). The bottom four sequences in the second half of the LeuAC-like deletions illustrate such features. They have an inserted segment that derives from two different regions of the full-length sequence (highlighted in the bottom panel in Fig. 3A). Blue background shading represents a duplication of the first half of the protozyme. It does not vary in either protozyme-like or LeuAC-like sequences. A short N-terminal segment (magenta background) derives from the C-terminus of both the LeuAC-like and Goldilocks urzymes, C-terminal to the AMSAS signature. The last 5 sequences in Fig. 3A are aberrant deletions and difficult to classify. They contain additional deletions in both N- and C-terminal segments. Their coding sequences are nevertheless free of point mutations.

### The deletions must occur *in vivo*

Full-length mutants at only one site do not produce any deletions. The process thus requires both sets of mutations. Nor does the reciprocal mutagenesis of double mutant plasmid with WT primers. These control experiments also mean that deletions derive not from the PCR mutagenesis itself. Rather, they must arise *in vivo,* after transformation. The cellular process is dependent on dual mutagenesis to the active site, hence therefore also one-way. We note as well that most, if not all ORFs have open reading frames in frame on the opposite strand. These ORFS cannot be annotated in terms of known contemporary protein sequences. We have not yet investigated whether or not the opposite strand ORFs might encode proteins homologous to Class II AARS as predicted by the Rodin-Ohno hypothesis (<u>Rodin and Ohno 1995</u>).

We cannot suggest a detailed mechanism, but the deletions must have been created and selected *in vivo*. We consider factors influencing selection of the coding sequences retained in the Discussion. We note here that they are likely coupled to DNA replication because mRNA synthesis from the pET-11a host plasmid is minimal without induction. Notably, those incubated exclusively at 37° C had no Goldilocks deletions and fewer tandemly repeated and multi-frame encoding than did those at 4° C. The enrichment of deletions at low temperature (Fig. 3C) is consistent with the known accessibility of D-loops to recombination proteins and replication restart at low temperature (<u>Michel and Sandler 2017</u>).

### The deletions encode active tRNA synthetases

AARS urzymes retained three catalytic properties seen in full length AARS. (i) They accelerate amino acid activation. (ii) They show significant pre-steady state bursts in single-turnover kinetic assays (<u>Fersht, et al. 1975</u>; <u>Francklyn, et al. 2008</u>). (iii) They accelerate aminoacylation of tRNA and minihelix substrates (<u>Tang, et al. 2024</u>). Clone21 and Clone22 (Fig. 3B), retain all three catalytic activities at levels comparable to those observed for the urzymes generated intentionally on the basis of structural biology (<u>Carter 2014</u>). Experimental kinetics data confirm all three criteria (Figs. 4-6).

**Figure 4.**
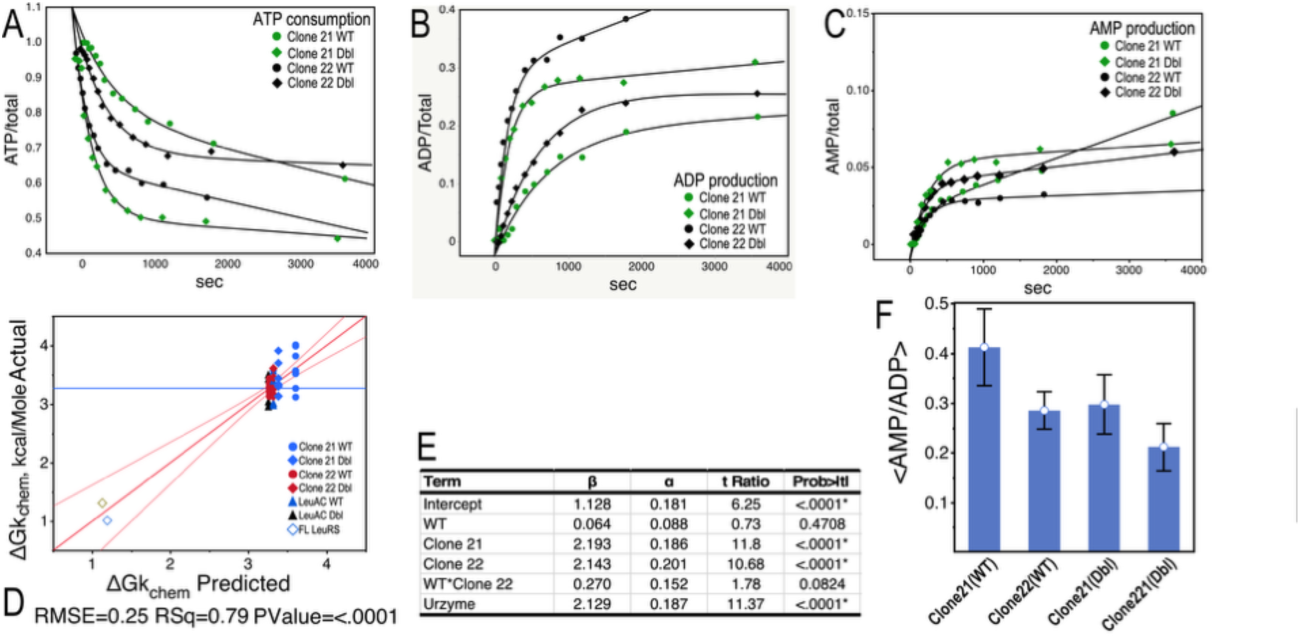
Single turnover analysis of the recombinant deletion urzymes. **A.** Time course for ATP consumption by all four variants. **B.** Time course for ADP production. **C.** Time course for AMP production. **D.** Regression model for the dependence of the first-order rate constant, k_chem_, on variant and mutant properties. **E.** Table of regression coefficients for the model in **D**, their estimated errors and P values. **F.** Histogram of the ratio of AMP to ADP produced by the four urzymes. WT Clone 21 has the highest ratio, hence uses ATP most efficiently. This ratio is consistent with the fact that the burst size of this variant in ADP production is lower than the others and the fact that the steady-state rate of AMP production is higher.

Fig. 4 summarizes single-turnover experiments on wild type (WT), double (AVGA/AMSAS; Dbl) Clone 21 and 22, as well as LeuAC urzyme (<u>Hobson, et al. 2022</u>; <u>Tang, et al. 2023</u>; <u>Tang, et al. 2024</u>) variants. Single-turnover experiments (<u>Fersht 1975</u>; <u>Francklyn, et al. 2008</u>) use large enzyme concentrations to build the first-round product formation into a measurable signal from the first round of catalysis. They use only a small excess of substrate; Most kinetic events are first order; hence they measure unimolecular reactions. If product release is rate-limiting, the bound product builds up in the enzyme. Such experiments show a pre-steady-state burst. Burst sizes represent the fraction of active enzyme molecules contributing to the experimental signal. These are shown in Supplementary Fig. S8. They range from ∼0.3 for the WT Clone 22 to ∼0.6 for the double mutant Clone 22 with both WT and double mutant Clone 21 values both close to 0.55. Single turnover data in Fig. 3 reveal differences between Clones 21 and 22.

Time courses of the concentrations of all three adenine nucleotides for all four variants (Fig. 4A-C) are similar to those observed previously for LeuAC (<u>Tang, et al. 2024</u>). The first-order rate constants for amino acid activation (Supplementary Table S1) are very nearly the same as those for LeuAC (Fig. 4D, E) and orders of magnitude slower than those of full-length LeuRS. Moreover, as we found for LeuAC (<u>Tang, et al. 2023</u>) the AVGA and AMSAS mutations actually enhance k_chem._ for Clone 21. But for Clone 22, which is slightly more active overall, that effect was reversed. P-values for the β-coefficients of the regression model in Figs. 4D and 4E) suggest that although the difference between the two variants is significant, the difference between the effects of mutant and WT active-site residues is smaller.

Single-turnover experiments yield two rate constants. The first-order rate constant, k_chem_, refers to the depletion of ATP in the chemical step. The turnover rate is k_3_. The mean ratios k_chem_/k_3_ for Clones 21 and 22 (230) are similar to that for LeuAC (223). Thus, the pre-steady state bursts are comparable. All retain the activated Ile-5’AMP with an increased affinity of ∼ –3.2 kcal/mole.

The ratio of AMP to ADP produced in such assays measures the efficiency ATP usage to produce activated amino acid (<u>Patra, et al. 2024</u>). Interestingly, the WT Clone 21 uniquely makes less ADP (green dots in Fig 3B) and more AMP. It makes more efficient use of ATP than the others (Fig. 4F).

Steady-state kinetics confirm that Clones 21 and 22 catalyze aminoacylation. The essence of translation in biology is forming a covalent bond between an amino acid and an RNA substrate. Our previous work with the LeuAC urzyme derived from LeuRS showed three crucial details (<u>Hobson, et al. 2022</u>; <u>Tang, et al. 2023</u>; <u>Tang, et al. 2024</u>). (i) The preferred RNA substrate, by a factor of ∼10, is the 35-nucleotide minihelix containing the acceptor stem and the TΨC-minihelix. (ii) The preferred amino acid substrate is isoleucine, not leucine. (iii) Catalysis of acyl transfer to RNA is minimally impacted by alanine substitution of the polar side chains in HVGA and KMSKS signatures. For these reasons, we measured rates for isoleucine and minihelix substrates for the new double mutant (Dbl) constructs generated *in vivo* as well as their counterparts (WT) with histidine and lysine in the respective signatures. Triplicate assays of acylation are shown in Fig. 5 and shown in Supplementary Table S2.

**Figure 5.**
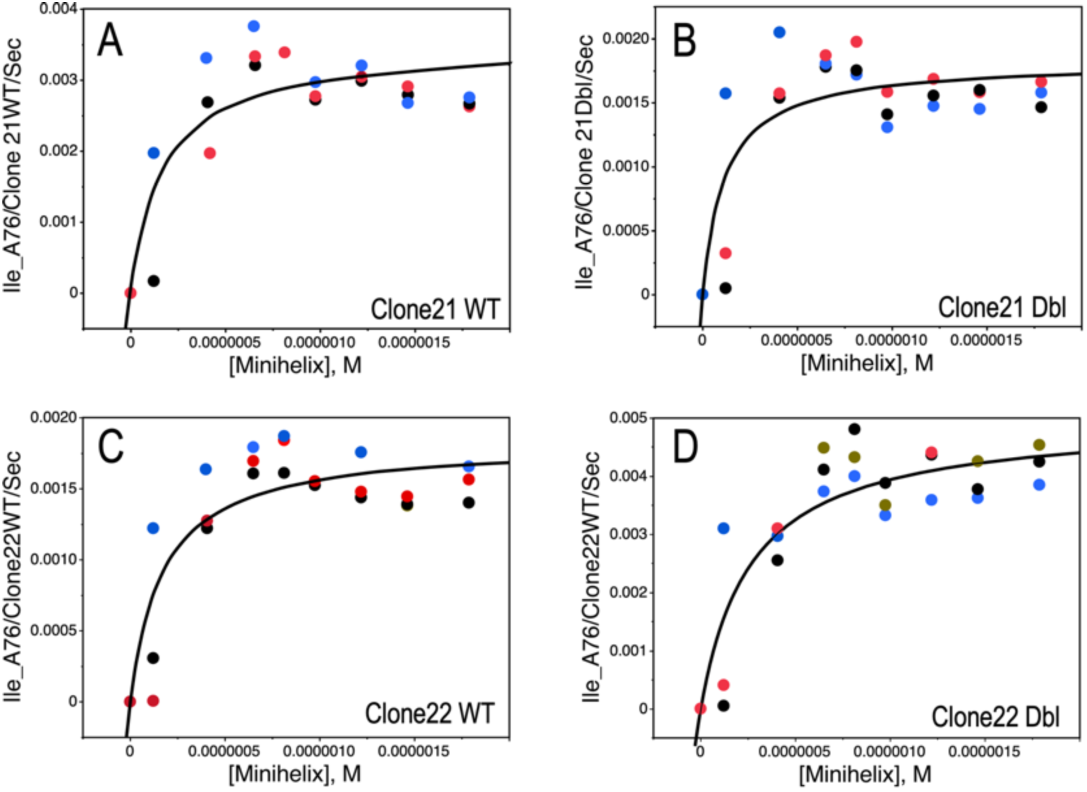
Michaelis-Menten plots of replicated aminoacylation assays. Aminoacylation of the leucine TΨC minihelix by Clone21, the Goldilocks urzyme with WT HVGH and KMSKS active site signatures (**A**) and the double mutant (**B**) sequences. **C, D.** Aminoacylation of the minihelix by Clone22 WT and DBL, respectively. Note that the vertical scales in **A-D** are different. The relative activities are compared in Fig. 5.

The distinction between single-turnover and steady state assays becomes blurred because we study catalysts that represent increasingly ancestral forms. As turnover decreases, steady-state assays require longer times and higher enzyme concentrations to build an adequate signal-to-noise ratio. Unpublished time-course data also suggest that the observed rates are, at least partially, first-order processes. ΔG(k_cat_) values from aminoacylation correlate with a linear combination of ΔG^‡^(k_chem_) and ΔG^‡^ (k_3_) values from single turnover experiments (R^2^ ∼ 0.97; P ∼ 0.03). Regression coefficients (see Supplemental Section E and Fig. S8) suggest that ΔG(k_chem_) contributes about 65% of the correlation and ΔG(k_3_) 35%. We have not found any way around this. It is a cost of working close to the boundary where experiments are no longer possible.

The curves in Fig. 5A, 5B, 5C do indeed suggest a decrease in observed rates at higher [minihelix]. The [substrate] necessary to detect dissociation between enzyme and minihelix RNA, and the apparent dissociation constants for the enzyme•RNA complex, ∼0.2 μMare an order of magnitude lower than the enzyme concentration. The RNA substrate is therefore rapidly consumed, and the acylated RNA product may remain bound, inhibiting further reaction by reducing [E_free_].

### Clones 21 and 22 connect the two active-site signatures in two, entirely new, functional ways

Fig. 3A shows that segments derived from the LeuRS active site are always present with their original sequences. That is what allows the neighbor joining algorithm used in Fig, 3B to identify only three clades. The original LeuAC design strategy preserved as much of the active site as possible. Only two short connections were made in order to delete the CP1 and CP2 peptides. Clones 21 and 22 have configurations that omit longer and different segments from those present in LeuAC. This leaves us puzzled as to how the two active-site signatures can assume similar configurations to those we imagine from the 1WZ2 crystal structure.

In turn, the significantly different linkages of the two parts of the active site introduce quite a new element into our thinking about ancestral Class I AARS (Fig. 6). LeuAC, designed *in vitro*, has 129 amino acid residues. The two *in vivo* LeuRS urzymes are both smaller. Clone 22 has residues 41-68 + 601-674 = 102 residues. Clone 21 has 41-95 + 650-674 = 81 residues. Yet, the averages of the four bars representing the WT and double mutant *in vivo* constructs are ∼ 2 kcal/mole more negative than the corresponding average for LeuAC (<u>Tang, et al. 2024</u>). Thus, they both catalyze minihelix acylation about 30 times better than the original LeuAC.

**Figure 6.**
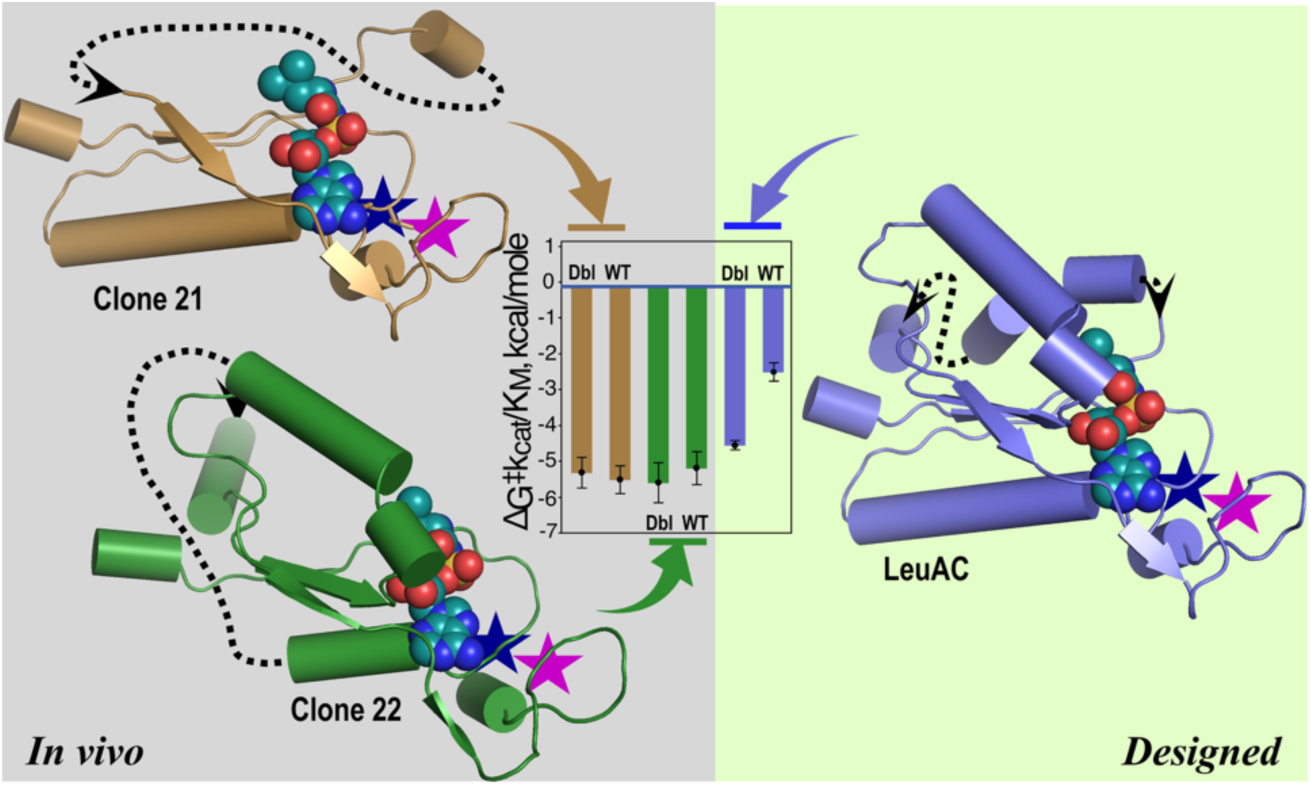
The N- and C-terminal parts of the LeuRS active site can be connected three distinct ways (dashed arrows) without changing the catalytic activity in RNA minihelix aminoacylation by either WT or double mutant forms (bar graph). The three active tRNA synthetases have, respectively, 81 (sand), 102 (green), and 129 (blue) residues. All cartoons are based on PDB 1WZ2. The construct on the right was previously designed to mimic the ancestral LeuRS. The two on the left were created *in vivo* by deletions to a full-length double mutant LeuRS plasmid. Wild type histidine (blue stars) and lysine (magenta stars) residues were added back by PCR mutagenesis for purposes of comparison.

### Regression modeling identifies and attributes selectable functions

The *in vivo* fragmentation allows us to analyze the enzymology of LeuAC at unprecedented resolution. The second crossover connection (missing in Clone 21) and second β-strand of the protozyme (missing in Clone 22), together with the WT vs double mutant active site residues form an incomplete factorial design (<u>Carter and Carter 1979</u>). The logarithmic transformation from rates to free energies of activation allow testing of linear models (<u>Weinreb, et al. 2012a</u>; <u>Li and Carter 2013</u>; <u>Tang, et al. 2023</u>; <u>Tang, et al. 2024</u>). Regression models for the Michaelis-Menten constants in Supplementary Table S2 show that the second half of the protozyme and the 2^nd^ crossover of the Rossmann fold have significant effects on the ΔG^‡^k_cat_, the *in* situ first-order process that transfers the acyl group to the cognate minihelix. Such effects offer an experimental basis on which to assess why particular modules might be selected.

Clones 21 and 22 are slightly better than the AVGA/AMSAS double mutant of LeuAC at aminoacylation. They aminoacylate minihelices significantly better than WT LeuAC does (Fig. 7). The enhanced values of k_cat_/K_M_ result both from increased k_cat_ (Fig. 7B) and higher affinity for minihelix RNA (Fig. 7C). Active-site mutations have opposite effects on the two *in vivo* constructs. The double mutant (Dbl) Clone 22 is the best catalyst of aminoacylation. It is nearly matched by the WT Clone 21. The WT Clone 22 is the least active variant. The decreased activity of WT variants that contain the second crossover connection of the Rossmann fold (i.e. LeuAC and Clone 22) has a P value <10^-4^. We noted a similar effect on the burst size (Supplementary Fig. S8).

**Figure 7.**
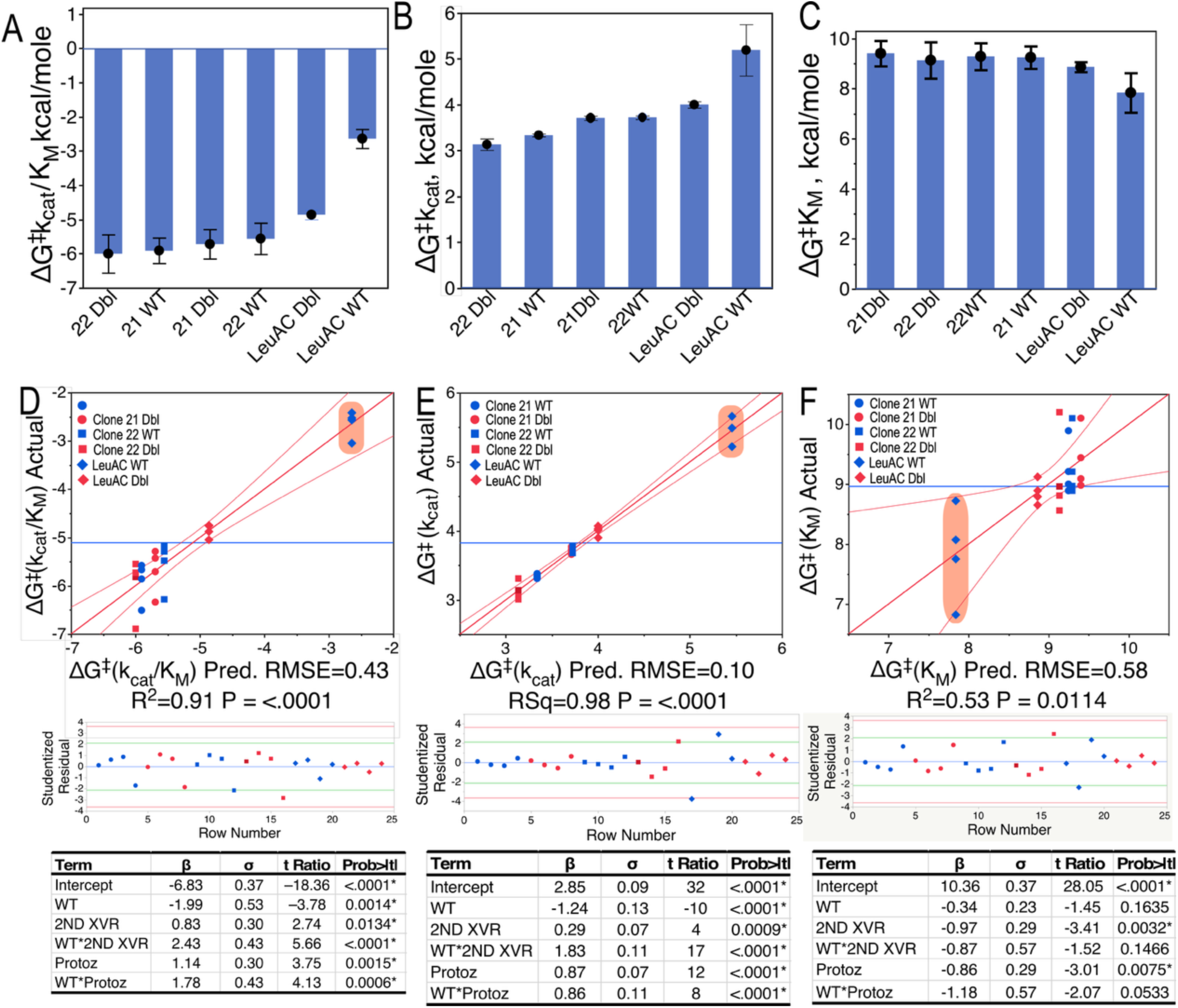
Michaelis-Menten parameters of AARS urzymes depend on the structural differences between constructs. Each plot is ordered in descending favorability from left to right. **A**. Bar plot of ΔG^‡^ (k_cat_/K_M_), the free energy for the second order rate constant for aminoacylation by reaction of free TΨC-minihelix with free substrate at saturating concentration of isoleucine, the amino acid substrate. **B**. ΔG^‡^(k_cat_), the free energy for the first-order rate constant. **C.** ΔG‡ (K_M_), the free energy for the apparent dissociation constant for the minihelix substrate. **D. E. F.** Regression models that relate steady-state parameters in **A-C** to the structural differences. Independent variables denote the following: WT refers to the presence of WT active-site histidine and lysine residues; 2ndXvr denotes the presence of the second crossover; Protoz denotes the presence of the intact protozyme at the N-terminus.

The wild-type active-site variants bind more tightly to substrate minihelix in both Clone 21 and Clone 22, (Fig. 7C), but not in LeuAC. All four *in vivo* variants are better catalysts in all respects than the double mutant LeuAC. All of the effects influencing ΔG^‡^k_cat_ have highly significant P-values (Fig. 7B). The WT active side residues change the effects of all three parameters. The second crossover connection reverses the effect of active-site mutations on Clone 22. This is apparent from the placement of the various symbols on the graph in both Fig. 7D and 7E.

The rose shading in all three plots shows that the kinetic properties of WT LeuAC are all significantly worse than all of the constructs produced *in vivo*. Estimation of higher-order effects is limited by the poor reproducibility of K_M_. Notwithstanding, significant evidence remains for a negative interaction between the second crossover connection and the second crossover connection of the Rossmann fold (Fig. 7D, E). That interaction reduces both ΔG^‡^k_cat_ and ΔG^‡^k_cat_/K_M_ by about 2 kcal/mole. Clones 21 and 22 therefore enhance studies of the evolution of protein modularity at unrivalled resolution.

### Class 1 and 2 AARS urzymes use similar secondary structures for aminoacylation

Clone 21 is the smallest functional Class I AARS gene ever produced. Clones 21 and 22 share the AMSAS signature. They differ in both the protozyme and second crossover. The C-terminal fragment shared by both Clone22 and Clone21 is key to binding RNA substrates. It is strong evidence that acquisition of that signature converted Class I protozymes into aminoacylating enzymes. Curiously, the secondary structure of the segment is a β-strand followed by a loop. The Motif 2 loop in Class 2 AARS also forms much of the RNA binding site. A striking resemblance of the RNA-binding secondary structures of Class 1 and 2 AARS is shown in Fig. 8. There are thus extraordinary similarities between the stereochemistry in both sets of polypeptide•RNA interactions leading to aminoacylation. We have suggested elsewhere how the contrasting substrate-binding interactions of Class I and II AARS can arise from the inversion symmetry of base-pairing in bidirectional ancestral genes (<u>Carter and Wills 2019a</u>; <u>Carter and Wills 2019b</u>, <u>2021</u>; <u>Carter 2024</u>; <u>Carter, et al. 2025</u>).

**Figure 8.**
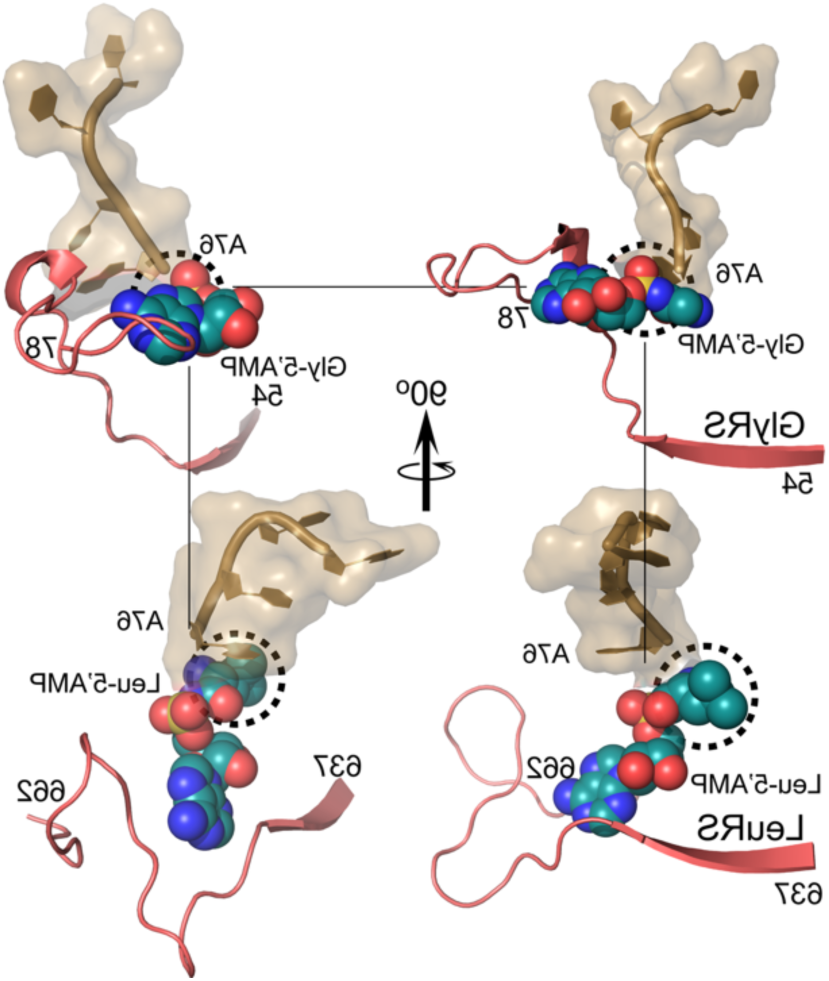
Corresponding segments of the active site that bind RNA substrates in Class 1 LeuRS (PDB ID 1WZ2) and Class II GlyRS (PDB ID 7YSE) have very similar secondary structures. Approximately orthogonal views of the complexes between RNA binding motifs in the two Classes with their respective activated amino acyl-5’AMP and RNA ligands. Dashed circles highlight the aminoacyl groups. The vertices of the square in the background indicate the 5’-phoshporyl group. They are located remarkably similarly with respect to the β-strand and loop. A chlorine atom that is part of the adenine ring in 7YSE has been omitted.

The modest size of the Class I AMSAS-containing module is reminiscent of the N-terminal fragment of the protozyme, which also is ∼25 residues long and has a similar β–>turn secondary structure and is key to the binding of ATP. In fact, the corresponding fragment in Class II AARS is actually the same fragment, residues 54-78 in Fig. 8. As we noted previously (<u>Tang, et al. 2023</u>), neither the histidine side chains in the HVGH nor the lysine residues in the KMSKS signatures are necessary for amino acid activation or aminoacylation. The full functionality of the Lys-to-Ala mutations implies that the function of that segment arises solely from polypeptide secondary structure.

### The Goldilocks urzyme suggests how one gene can encode both Class I, II AARS

The Rodin-Ohno hypothesis that bidirectional ancestral AARS genes coded for Class I and II urzymes on opposite strands (<u>Rodin and Ohno 1995</u>) led to a surprising number of retrodictions about the Class distinction (<u>Carter, et al. 2025</u>). The bidirectional protozyme gene was readily constructed (<u>Martinez-Rodriguez, et al. 2015</u>) because the corresponding segments from Class I and II AARS have the same number of amino acids. Others have verified its activation activity (<u>Onodera, et al. 2021</u>). We have failed, however, to design a bidirectional gene encoding Class I and II AARS urzymes on opposite strands because there are several loci where the combination of the two signatures has different numbers of amino acids and cannot be fitted into a consistent antiparallel alignment.

Because it lacks the second crossover connection of the Rossmann fold, Clone 21 is the same length (81 residues) as the Class I GlyCA urzyme (<u>Patra, et al. 2024</u>). Further, 81-residue Goldilocks excerpt is a catalytically active AARS. Moreover, the Class-defining signature sequences of Class I and II AARS can both be readily aligned opposite one another in antiparallel alignments by the homology suggested by the GlyCA structure (<u>Carter, et al. 2025</u>). Thus, the Goldilocks urzyme, Clone21, may show how to explore designed bidirectional Class I/II AARS urzyme genes.

We compare in Fig. 9 antiparallel alignments of Goldilocks-length urzymes drawn from 10-12 species of each synthetase from those in <u>aars.online</u> (<u>Douglas, Cui, et al. 2024</u>). The three Class I AARS subclassses match uniquely with Class II subclass urzyme genes. Superior matches entail both the total length and significantly elevated codon middle-base pairing. From the table in Fig. 9B, Class IA IleRS pairs best with IIB LysRS; Class IIA ProRS pairs best with Class IB GlnRS. Class IC TrpRS pairs best with Class IIC HisRS. We previously outlined extensive tests of the pairing frequencies under the null hypothesis of no bidirectional coding ancestry for such alignments (<u>Chandrasekaran, et al. 2013</u>). Standard errors for all mean values in Fig. 9B are derived from ∼140 bidirectional alignments. They about 1 percent of the values themselves. The middle-base pairing frequencies are thus all highly non-random and statistically different one from another. Moreover, the different lengths of the three sets of related pairings make it hard to evaluate cross pairing in the matrix in Fig. 9B.

**Figure 9.**
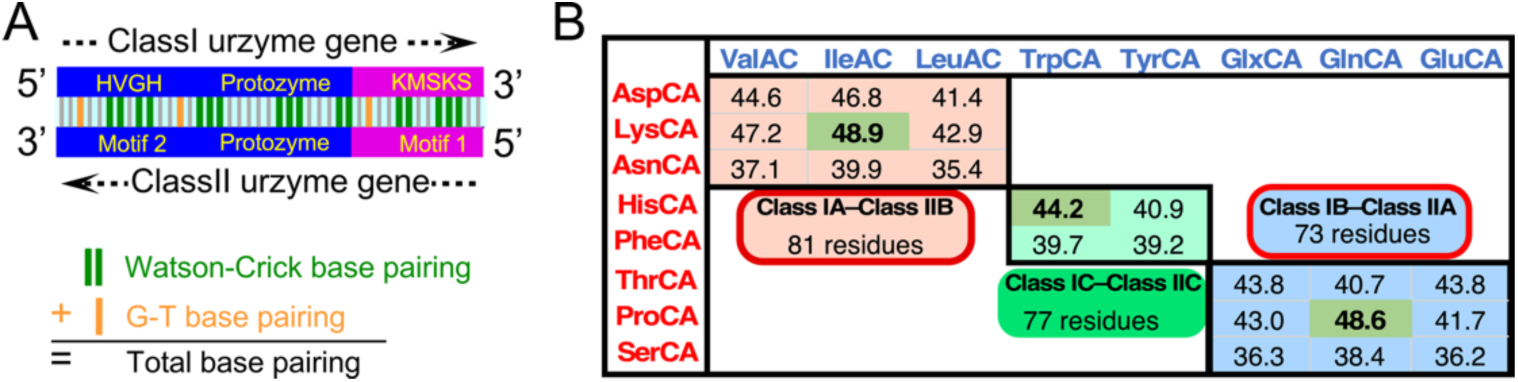
Goldilocks urzyme, Clone21, suggests a paradigm for designing bidirectional urzyme genes for Class I and II ancestors on opposing DNA strands. **A.** Schematic of sense/antisense encoding of Class I and II AARS urzyme genes, consistent with the architecture of the two AARS Classes. Vertical lines between the two genes suggest bases that would be paired in a bidirectional gene but are only partially paired in sequences derived from contemporary genes. **B.** Percentages of codon middle-base pairing in antiparallel alignments, calculated as indicated in **A** are summarized for three groups derived from the three consensus synthetase subclasses (<u>Carter, et al. 2025</u>). Values greater than ∼30% are highly unlikely under the null hypothesis.

Pairings in Fig. 9B conflict with the conventional sub-classification by pairing subclass A in Class I with subclass B in Class II and vice versa. Each such pairing brings along two additional AARS from the same respective subclasses. Thus, 16 of the 20 bacterial AARS sequence alignments show analogous pairings between Class I and II AARS genes (<u>Carter, et al. 2025</u>).

## DISCUSSION

We describe a novel host-plasmid interaction that occurs when a PCR-generated, nicked plasmid containing a Leucyl-tRNA synthetase double mutant is introduced into *E. coli*. LeuRS is a large, multidomain protein. Transforming *E. coli* with a variant containing mutations in two remote parts of the active site gives rise to a high proportion (∼40%) of a discrete set of very long deletions. Most of the short ORFs retain sequences encoding active leucyl-tRNA synthetases that resemble the LeuAC urzyme (<u>Hobson, et al. 2022</u>). The rest encode a shortened variant of the LeuRS protozyme.

### The LeuRS ORFs made *in vivo* validate previous studies of AARS ancestors

This result unexpectedly confirms many aspects of our *in vitro* studies of ancestral AARS (Pham, et al. 2007; Pham, et al. 2010; Li, et al. 2011; Li and Carter 2013; Li, et al. 2013; Carter 2014, 2015; Martinez-Rodriguez, et al. 2015; Carter and Wills 2021). We had thought that the full protozyme and 2^nd^ crossover segments were essential for acylation. Neither is actually required. Catalytic proficiencies of Clones 21, 22 are even better assignment catalysts than the LeuAC double mutant (see Fig. 6 and K_sp_ values in SupplementalTable S2). It is hard to overstate the significance of these parallels. They substantiate our assertion that AARS urzymes are valid models for early AARS evolution.

Analytical structure-based deconstruction took us ten years. Here, we found that *E. coli* creates ORFs *in vivo* and in a week or so, that are uncommonly similar to those we previously described. The Clone 21 and 22 deletions represent at least four discoveries we otherwise would not have made: (i) the mutant AVGA and AMSAS active-site signatures can form functional linkages in surprisingly simple ways we had not considered (Fig. 6). (ii) the module that enables the Class I protozyme to recognize and acylate RNA substrates is a short segment including the downstream AMSAS signature. That signature forms a structure quite similar to that of the RNA binding site of Class II AARS. (iii) the Goldilocks urzyme shows that the second crossover connection of the Rossmann fold is not required for catalysis of aminoacylation (Fig. 7). (iv) From that we also discovered how to configure bidirectional urzyme genes (Fig. 9).

Each of these new clues would have made it easier for nature to find a reflexive set of protein assignment catalysts to administer the coding table. A coherence of sequence and function unites the data in Figs. 1-5. That coherence (Fig. 6) suggests that although we were certainly on the right track working exclusively from superposing tertiary structures to infer ancestral AARS properties. *E. coli* has created similar catalytically active Class I urzymes that are different from those we had designed computationally and characterized experimentally. In fact, it has shown us how we can do an even better job. The full significance of Fig. 6 is that we were sampling what turns out to be a much broader functional landscape.

### New evidence for the modular origin of protein-based urzymes for both AARS Classes

The three new ORFs created *in vivo* complement previously reported hypotheses about the origins of protein-based AARS. We can now infer five new aspects about how amino acid activation and aminoacylation first emerged from the two Class I active-site signatures. (i) They have distinct functionality–ATP utilization (<u>Martinez-Rodriguez, et al. 2015</u>) and substrate RNA recognition (Fig. 8). (ii) They act somewhat independently because they can be linked together differently without disrupting function. (iii) The protozyme and C-terminal fragments (Fig. 8) suffice to catalyze both reactions needed for codon-directed protein synthesis. (iv) They need not even contain the complex amino acid side chains by which they first were identified. Finally, (v) we can now see how genes for Class I and II AARS urzymes can be aligned on opposite strands of a single bidirectional gene (Fig. 9).

The longer-term impact for the study of the origin of genetics is hard to underestimate. These results, together with previous work, provide new and broadly-based support for a model for the emergence of the first Class I protein AARS. We envision assembly from three peptide fragments, each about 25 residues long. There is significant evidence for exceptionally high codon middle-base pairing between antiparallel alignments of each fragment. Bidirectional ancestral genes could thus have provided each fragment, establishing a basis for the joint origin of aminoacylating urzymes of both AARS classes (Fig. 10).

**Figure 10.**
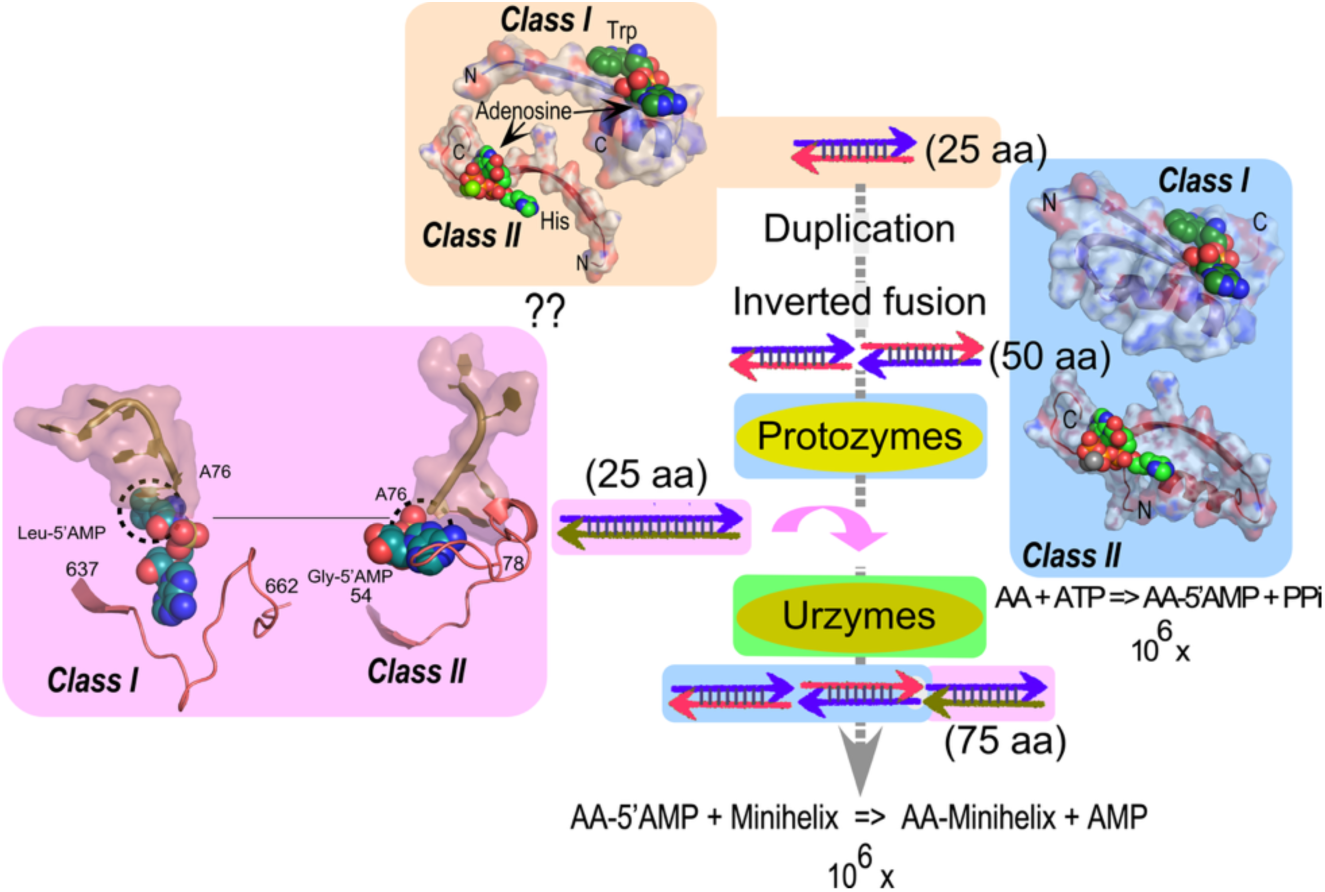
Key steps in the emergence of protein AARS. *In vivo* fragmentation of the LeuRS gene described here provides overlapping reinforcement for the idea that amino acid activating and RNA aminoacylation activities can be attributed to three polypeptide fragments each about 25 amino acids long highlighted by sand, blue, and magenta backgrounds, as described in the text. The two reactions catalyzed by all AARS are indicated with their respective rate accelerations, as shown in Figs. 3,4.

One fragment (sand background) provides an ATP binding site. The new result that both the half-protozyme and LeuAC-like ORFs retain this fragment unexpectedly supports its key role. To date we have only the structural evidence in the molecular cartoons that this fragment actually binds ATP. It does, however, contain the highly conserved ATP binding surfaces found in crystal structures from both Classes.

We proposed the half-protozyme as a progenitor of the Class I AARS protozyme (blue background) via gene duplication and fusion to form an inverted repeat (<u>Chandrasekaran, et al. 2013</u>). Protozymes with both contemporary and bidirectionally coded sequences accelerate amino acid activation by 10^6^-fold. The third fragment (magenta background) endows the protozyme with the additional ability to recognize RNA substrates and catalyze acyl-transfer. It is described here for the first time (Fig. 8) showing the similarity between polypeptide:RNA interactions in the two AARS Classes. It appears to be both necessary and sufficient to enable the protozyme to catalyze aminoacylation (Fig. 6).

Differences between the three *in vivo* ORFs described here and the corresponding species we created *in vitro* may allow more precise specification of the actual modularity of ancestral AARS. To that end, it will be of interest to deepen understanding of the underlying *in vivo* processes. We plan to follow the distribution of plasmid sizes and the functionality of their apparent coding regions as a function of time following transformation and at different levels of induced expression.

### Profiling short plasmids accompanying cloned mutations may help access other primordial genes

A central question about the origin and early evolution of translation is how to connect the much shorter coding sequences of ancestral AARS quasispecies to conventional ancestral sequence reconstructions. The latter are currently limited to essentially full-length species that had already been assembled by the time of the last universal common ancestor, LUCA (<u>Fournier, et al. 2011</u>; <u>Fournier and Alm 2015</u>). It seems likely that some process or processes added new genetic material as modules of significant size. How did AARS genes undergo that modular growth?

We argue that the LeuRS ORFs also signify a new and potentially general approach to the discovery of ancestral genes, both for AARS and perhaps also for other important protein families. Our argument is based on previously published evidence that appropriate double mutants disrupt coupling networks in the intact LeuRS sufficiently to cause metabolic damage. Although pET-11a should minimize expression in the absence of inducer IPTG, it is well-known that variable amounts of leakage invariably occur. For example, few, if any of the wide range of plasmids with mutant AARS we have made in that vector can be maintained as transformed cultures. Expression invariably requires fresh transformation.

It is remarkable that two of the three LeuRS ORFs preserve both AARS catalytic activities, albeit at greatly reduced levels. The simple coherence of the LeuRS ORFs (Fig. 9) suggest that some differential fitness may have led to their survival. We consider several questions in turn.

#### Why do so few variations survive competition with the full-length LeuRS double mutant?

The rate enhancement for tRNA aminoacylation by the double mutant is nearly an order of magnitude less than that for WT LeuRS (<u>Tang, et al. 2024</u>). That loss of function would mean that the WT enzyme present in the transformed cells should produce adequate cellular Leucyl-tRNA^Leu^. Loss of both domains only when both active site loci are mutated suggests that the double mutation causes a toxic gain of function. Active-site mutations increased acylation with isoleucine by LeuRS (<u>Tang, et al. 2023</u>; <u>Tang, et al. 2024</u>). Similarly, TrpRS mutants lacking either CP1 or the ABD increased activation of tyrosine (<u>Li and Carter 2013</u>; <u>Weinreb, et al. 2014</u>). Mischarging of tRNA is also considered a likely possible mechanism for causing genetic disease by mutant mitochondrial AARS (<u>Meyer-Schuman and Antonellis 2017</u>). Mischarging of tRNA by the double mutant full-length LeuRS is thus a possible culprit.

#### Why are both CP1 and ABD domains deleted entirely in all shortened plasmids?

It seems significant to us that the connecting peptide (CP1) and anticodon-binding (ABD) domains are both entirely missing in all deletions. Our earlier work on TrpRS (<u>Weinreb, et al. 2012a</u>; <u>Li and Carter 2013</u>; <u>Weinreb, et al. 2014</u>; <u>Chandrasekaran, et al. 2016</u>; <u>Carter, et al. 2017</u>; <u>Chandrasekaran and Carter 2017</u>) and LeuRS (<u>Tang, et al. 2023</u>) quantified an essential energetic coupling of the functional contribution of these two domains to catalysis and specificity to the active-site histidine and lysine residues. The four catalytic side chains increase both the rates and specificities of both Class I AARS by five orders of magnitude. However, the functional contributions of the catalytic side chains occurs if and only if both domains are present (<u>Li and Carter 2013</u>).

That linkage between the active-site and the two domains appears to be mediated by two different aspects of the tertiary structure in the full-length proteins. The first is a switching interaction within the urzyme. We identified (<u>Kapustina, et al. 2007</u>) and analyzed (<u>Weinreb, et al. 2012a</u>, <u>b</u>) that switching interaction in considerable detail in TrpRS. The second part of the coupling is a packing interaction between non-polar side chains in the ABD that provides “sockets” for the unique non-polar side chains, H**V**GH and K**M**SKS, in each catalytic signature (<u>Tang, et al. 2023</u>). Its role is to assure that the active site is sensitive to the arrangement of the domains. Both aspects appear characteristic of other (<u>Alexander and Schimmel 2001</u>; <u>Budiman, et al. 2007</u>; <u>Li, et al. 2015</u>) and perhaps all Class I AARS.

The linkage described in the previous paragraphs provides all-or-none functionality. If either histidine or lysine residues, or if one of the two domain is missing, the substantial rate enhancement and specific substrate recognition are lost. Neither domain can provide its normal function if both active-site loci are mutated. That interdependence suggests why both domains are lost.

Mischarging of tRNA is possible toxic gain of function in the present case. It would explain the discrete spectrum of deletions. The ten-fold preference for TΨC minihelix over tRNA (<u>Tang, et al. 2024</u>) means that deletion of both domains to produce urzyme-like species would be expected to reduce mischarging of full-length tRNA 10^7^-fold. That also shows that mischarging can be minimized by deleting major parts of the mature protein. Our studies of the Class I AARS mechanistic enzymology thus suggest a rationale for which variants survive.

#### What mechanisms might have favored the removal of such long segments of genetic information?

The deletions we observe must result from DNA metabolism, because they are inherited in the recovered plasmids. The genomic features summarized earlier (e.g. Fig. 3C and Supplemental Figs. 1-7) suggest multiple exchanges between different segments of coding information. These events likely occur during DNA replication, perhaps by formation of secondary or tertiary structures that promote stem-loop formation (<u>Bikard, et al. 2010</u>). The *E. coli* strain carries a RecA^−^ gene. Any recombination likely entailed other pathways (<u>Ivancic-Bace, et al. 2005</u>).

Whatever the mechanism, we note that such processes have been observed in other systems. A notable and functionally significant example is the proposed evolutionary path of the *Drosphophila melanogaster* SLIMP•SerRS heterodimer observed in mitochondria from many metazoans. The SLIMP (SerRS-Like Insect Mitochondrial Protetin) evolved by domain collapse and active-site ablation (<u>de Potter, et al. 2023</u>).

The ORFs in Clones 21, on the one hand, and the protozyme-like and Clones 22 on the other hand, are not compatible with successive recombinational events. The former contains the intact protozyme whereas the latter contains only the N-terminal half of the protozyme (Figs. 3A and 6). They must therefore arise independently. These distinctly different configurations of the two active site signatures and the fact that the Goldilocks urzyme (Clone21) appears mostly after prolonged incubation of colony plates at 4° C (Fig. 3C) suggest that multiple mechanisms may be required to explain the variety of deletions.

Another possible mechanism involves the inadvertent induction of the CRISPR/Cas defense machinery. As noted by Markulin and colleagues (<u>Markulin, et al. 2020</u>) CRISPR systems are activated only below 30° C. None of these arguments provide full explanations of the observations we report here. For example, we cannot explain why we do not see mutations that create pseudogenes that are no longer expressed.

#### Why might other complex proteins behave similarly?

What we observed may occur when similar experiments are done with a wider range of AARS and other complex multi-domain proteins. Many important protein families—enzymes, motor proteins, regulatory proteins—share a complex multidomain configuration with LeuRS. It has long been apparent that using intramolecular communication to induce functional cooperativity is an emergent property of such proteins (<u>Monod, et al. 1963</u>; <u>Monod, et al. 1965</u>; <u>Changeux and Edelstein 2005</u>; <u>Sadovsky and Yifrach 2007</u>; <u>Zandany, et al. 2008</u> ; <u>Ben-Abu, et al. 2009</u>; <u>Carter 2017</u>). It appears to be an important source of the fitness that supported the (rapid) evolutionary gain of new genetic sequences.

Although the details of coupling differ in many ways from protein family to protein family, it is irrelevant precisely how they function. The specific fragmentation we observed suggests that disrupting internal coupling networks can perturb the fitness of cloned proteins. Other protein families may use coupling mechanisms that can be made cytotoxic by widely spaced active-site mutations. Our experiments show that one way to eliminate cytotoxic effects of cloned variant full-length proteins is to reduce those variants to their simplest functional forms.

Our models for ancestral AARS show that the most highly conserved modules retain substantial catalytic activity. They also reduce putative cytotoxicity. Experiments similar to those we describe here may produce similar sets of much shorter, but still functional variants of many other proteins. The possibility of a wider generality of our protocol could provide a rapid access to urzyme-like variants. The eventual utility of such variants would have evolutionary significance. Such variants could also provide a means to reduce the large size of genes for delivery by viral capsid systems (<u>Hirsch, et al. 2010</u>; <u>Lewis 2014</u>). Protein engineering (<u>Kuhlman and Baker 2004</u>; <u>Romeroa, et al. 2018</u>; <u>Dauparas, et al. 2022</u>; <u>Dauparas, et al. 2023</u>) can further enhance function (<u>Patra, et al. 2025</u>).

## DATA AVAILABILITY

Data that are not provided in the supplement will be made available upon reasonable request.

## ACKNOWLEDGMENTS

This work was supported by the Matter-to-Life program of the Alfred P Sloan Foundation, Grant G-2021-16944. We are grateful for generous assistance from P. R. Wills and J. Douglas on the alignments in Fig. 8. We also thank H. Fried (Cursor Scientific Editing and Writing) for stylistic advice on the manuscript.

## SUPPLEMENTARY INFORMATION

Complete sequence alignments and additional information are provided in the file: Recombinant_deletions_supplement.pdf

