## Supplemental data tables and figures for "*Escherichia coli* deletes *in vivo* the same domains from a double-mutant leucyl-tRNA synthetase gene that were deleted *in vitro* to make the LeuAC urzyme"

***Escherichia coli* converts a double-mutant leucyl-tRNA synthetase gene into primordial genes *in vivo***

Supplemental Data

A. Raw sequencing data for 79 plasmids containing recombinant deletions from the Full-length double mutant LeuRS plasmid. These data can be found in the files:

Gene_Assembly_1.xltx

Gene_Assembly_2.xltx

Gene_Assembly_3.xltx

Gene_Assembly_4.xltx

Gene_Assembly_6.xltx

Note that in all of the excel files and subsequent supplemental figures, filenames beginning with 17A were incubated exclusively at 37^o^ C and those beginning with 19A were incubated overnight at 37^o^ C and then transferred to 4^o^ C.

B. Fasta file of selected alignments used in Fig. 1A of the main text. Sequences have been curated and aligned manually to facilitate comparison between one another.

>LeuAC_WT_A/1-129

----------------------------------------EKKFYITVAFPYLSGHLHVGHARTYTIPDVIA

RFKRMQGYNVLFPMAWHITSLS--------------------------------------------------

------------------------------------------------------------------------

------------------------------------------------------------------------

------------------------------------------------------------------------

------------------------------------------------------------------------

------------------------------------------------------------------------

------------------------------------------------------------------------

-------------------DSTIYMAYYTFEYWYPLDWRCSGKDLIPNHLTFFIFNHVAIFREEHWPKGIAV

NGFGTLEGQKMSKSKGNVLNF---------------------------------------------------

------------------------------------------------------------------------

------------------------------------------------------------------------

------------------------------------------------------------------------

------------------------------------------------------------------------

------------------------------------------------------------------------

------------------------------------------------------------------------

------------------------------------------------------------------------

------------------------------------------------------------------------

------------------------I

>LeuRS_B/1-967

M-------AELN-FKAIEEKWQKRWLEAKIFEPNIRDKPKEKKFYITVAFPYLSGHLHVGHARTYTIPDVIA

RFKRMQGYNVLFPMAWHITGSPIVGIAERIKNRDPKTIWIYRDVYKVPEEILWTFEDPINIVKYFMKAAKET

FIRAGFSVDWSREFYTTSLFPPFSKFIEWQFWKLKEKGYIVKGAHRVRWDPVVGTPLGDHDLMEGEDVPILD

YIIIKFELRENGEVIYLPAATLRPETVYGVTNMWVNPNATYVKAKVRRKDKEETWIVSKEAAYKLSFQDREI

EVIEEFKGEKLIGKYVRNPVSGDEVIILPAEFVDPDNATGVVMSVPAHAPFDHVALEDLKRETEILEKYDID

PRIVENITYISLIKLEGYGDFPAVEEVNKLGIKSQKDKEKLEQATKTIYKAEYHKGIFKVPPYEGKPVQEVK

EAIAKEMLEKGIAEIMYEFAEKNVISRFGNRAVIKIIHDQWFIDYGNPEWKEKARKALERMKILPETRRAQF

EAIIDWLDKKACARKIGLGTPLPWDPEWVIESLSDSTIYMAYYTISRHINKLRQEGKLDPEKLTPEFFDYIF

LEEFSEDKEKELEKKTGIPAEIIHEMKEEFEYWYPLDWRCSGKDLIPNHLTFFIFNHVAIFREEHWPKGIAV

NGFGTLEGQKMSKSKGNVLNFIDAIEENGAD-----------------------------------------

---------------------------------------VV-------------------------------

--------------------RLYI--------MS---------------------------------LAEHD

SDFDWRRKEVGKLRKQIERFYELISQFAEYEVKGNVE--------LKDIDRWMLHRLNKAIKE-TTNALEEF

RTRTAVQWAFYSIMNDLRWYLR--------------------------------------------------

--------------------RTEGRDDEAKRYVLRTLADVWVRLMAPFTPHICEELWEKLGGEGFVSLAKWP

EPVEEWWNETI-----------------------------------EAEEEFIRSVMEDIKEIIEVAKIENA

KRAYIYTAEDWKWKVAEVVSEKRDFKSSMEEL----------MKDS------EIRKHGKEVAKIVQKLIKER

TFDVKRIN--EEKAL--------------RE--------------------------AKEFMEKELGIEIII

NPTEDKGGKKKQAMPLKPAIFIE

>17A_028_AVGAAMSAS-8_AVGA-sqn1_D04/1-282

-------------FLSELRKMAERWLEAKIFEPNIRDKPKEKKFYITVAFPYLSGHLAVGAARTYTIP----

------------------------------------------------------------------------

------------------------------------------------------------------------

------------------------------------------------------------------------

------------------------------------------------------------------------

------------------------------------------------------------------------

------------------------------------------------------------------------

------------------------------------------------------------------------

-------------------AEIIHEMKEEFEYWYPLDWRCSGKDLIPNHLTFFIFNHVAIFREEHWPKGIAV

NGFGTLEGQAMSASKGNVLNFIDAIEENGADVVRLYIMSLAEHDSDFDWRRKEVGKLRKQIERFYELISQFA

EYEVKGNVELKDIDRWMLHRLNKAIKETTNALEEFRTRTAVQW--AFYSIMNDLRWYLRRTEGRDDEAKR--

----------------------YVLRTLADVWV-------------RLMAPFTPHICEELWEKL--------

GG----------------------------------------------------------------------

------------------------------------------------------------------------

------------------------------------------------------------------------

------------------------------------------------------------------------

------------------------------------------------------------------------

--------------E---------------------------------------------------------

---------------------AL

>19A_008_21-AVGA-AMSAS-AVGAsqun1-08_AVGAsqun1_H01/1-289

-------------F-KRLRKWQKRWLEAKIFEPNIRDKPKEKKFYITVAFPYLSGHLAVGAARTYTIP----

------------------------------------------------------------------------

------------------------------------------------------------------------

------------------------------------------------------------------------

------------------------------------------------------------------------

------------------------------------------------------------------------

------------------------------------------------------------------------

------------------------------------------------------------------------

-------------------AEIIHEMKEEFEYWYPLDWRCSGKDLIPNHLTFFIFNHVAIFREEHWPKGIAV

NGFGTLEGQAMSASKGNVLNFIDAIEENGADVVRLYIMSLAEHDSDFDWRRKEVGKLRKQIERFYELISQFA

EYEVKGNVEPKEKKFYITV-----------AFPYLSGHLAVGA--A--------------------------

--------------------RTYT--------IPAEIIHEMK-EEFEYWYPLDWRCSGKDLIPN--------

HLTFFIFTHVAIFREE--------------------------------------------------------

-----------------HLAE---RHCGERL---------W--HP--GGQAMIAT-----------------

------K-----------------------------------------------------------------

------------------------------------------------------------------------

------------------------------------------------------------------------

--------------A---------------------------------------------------------

---------------------MC

>19A_014_21-AVGA-AMSAS-AVGAsqun1-14_AVGAsqun1_F02/1-238

------------------RKWQKRWLEAKIFEPNIRDKPKEKKFYITVAFPYLSGHLAVGAARTYTIP----

------------------------------------------------------------------------

------------------------------------------------------------------------

------------------------------------------------------------------------

------------------------------------------------------------------------

------------------------------------------------------------------------

------------------------------------------------------------------------

------------------------------------------------------------------------

-------------------AEIIHEMKEEFEYWYPLDWRCSGKDLIPNHLTFFIFNHVAIFREEHWPKGIAV

NGFGTLEGQAMSASKGNVLNFIDAIEENGADVVRLYIMSLAEHDSDFDWRRKEVGKLRKQIERFYELISQFA

EYEVKGNVELKDIDRWMLHRLNKAIKETTNALEEFRTRTAVQW--AFYSIMNDLRWYLRRTE----------

------------------------------------------------------------------------

------------------------------------------------------------------------

------------------------------------------------------------------------

------------------------------------------------------------------------

------------------------------------------------------------------------

------------------------------------------------------------------------

--------------R---------------------------------------------------------

---------------------PR

>19A_044_22AVGA-AMSAS-AVGAsqun1-04_AVGAsqun1_D06/1-348

------------------RKWQKRWLEAKIFEPNIRHKPKEKKFYITVAFPYLSGHLAVGAARTYTIP----

------------------------------------------------------------------------

------------------------------------------------------------------------

------------------------------------------------------------------------

------------------------------------------------------------------------

------------------------------------------------------------------------

------------------------------------------------------------------------

------------------------------------------------------------------------

-------------------AEIIHEMKEEFEYWYPLDWRCSGKDLIPNHLTFFIFNHVAIFREEHWPKGIAV

NGFGTLEGQAMSASKGNVLNFIDAIEENGADVVRLYIMSLAEHDSDFDWRRKEVGKLRKQIERFYELISQFA

EYEVKGNVELKDIDRWMLHRLNKAIKETTNALEEFRTRTAVQW--AFYSIMNDLRWYLRRTEGRDDEAKR--

----------------------YVLRTLADVWV-------------RLMAPFTPHICEELWEKL--------

GGEGFV-------SLA--------------------------------------------------------

-----------------KWPE---PVE--EW---------WNETI--EA-----EEEFIRSVMEDIKEIIEV

AKIETRNARIFNTAE---------------------------------------------------------

----------IEMESGGVV----S------------------------------------------------

-----------------------------------------------ENA----------------------

------------ILK--------------------------------------------AA-----------

-------------------------

>19A_052_22AVGA-AMSAS-AVGAsqun1-12_AVGAsqun1_D07/1-366

-----------T-FLSRLRKWQKRWLEAKIFEPNIRDKPKEKKFYITVAFPYLSGHLAVGAARTYTIP----

------------------------------------------------------------------------

------------------------------------------------------------------------

------------------------------------------------------------------------

------------------------------------------------------------------------

------------------------------------------------------------------------

------------------------------------------------------------------------

------------------------------------------------------------------------

-------------------AEIIHEMKEEFEYWYPLDWRCSGKDLIPNHLTFFIFNHVAIFREEHWPKGIAV

NGFGTLEGQAMSASKGNVLNFIDAIEENGADVVRLYIMSLAEHDSDFDWRRKEVGKLRKQIERFYELISQFA

EYEVKGNVELKDIDRWMLHRLNKAIKETTNALEEFRTRTAVQW--AYYTISRHIN-KLRQ-EGKLDPEKLTP

EFFDYIFLEEFSEDKEKELEKKTG--------IPAEIIHEMK-EEFEYWYPLDWRCSGKDLIPN--------

HLTFFIFNHVAIFREE--------------------------------------------------------

-----------------HWAE---RHCGERL---------W--HP--GSQAMSAS-----------------

------KGNVLNFID---------------------------------------------------------

----------AIEENGADV----V------------------------------------------------

------------------------------------------------------------------------

------------RLL-----------------------------------------------------YL--

-------------------------

>19A_011_21-AVGA-AMSAS-AVGAsqun1-11_AVGAsqun1_C02/1-153

------------------RKWQKRWLEAKIFEPNIRDKPKEKKFYITVAFPYLSGHLAVGAARTYTIP----

------------------------------------------------------------------------

------------------------------------------------------------------------

------------------------------------------------------------------------

------------------------------------------------------------------------

------------------------------------------------------------------------

------------------------------------------------------------------------

------------------------------------------------------------------------

-------------------AEIIHEMKEEFEYWYPLDWRCSGKDLIPNHLTFFIFNHVAIFREEHWPKGIAV

NGFGTLEGQAMSASKGNVLNFIDAIEEKW-QKR--------------------WL------EAKIF------

---EPNIR----------------------------------------------------------------

---------------------------------------D-KPK----------------------------

------------------------------------------------------------------------

------------------------------------------------------------------------

------------------------------------------------------------------------

------------------------------------------------------------------------

------------------------------------------------------------------------

------------------------------E------------------------------K----------

-------------------------

>19A_054_22AVGA-AMSAS-AVGAsqun1-14_AVGAsqun1_F07outframe_Urzyme/1-227

-------------FLRRLRQWQKRWLEAKIFEPNIRDKPKEKKFYITVAFPYLSGHLAVGAARTYTIP----

------------------------------------------------------------------------

------------------------------------------------------------------------

------------------------------------------------------------------------

------------------------------------------------------------------------

------------------------------------------------------------------------

------------------------------------------------------------------------

------------------------------------------------------------------------

-------------------AEIIHEMKEEFEYWYPLDWRCSGKDLIPNHLTFFIFNHVAIFREEHWPKGIAV

NGFGTLEGQAMSASKGNVLNFIAFPY----------------------------------------------

-------------------------------------GHLAVGA----------------------------

--------------------ARTYT--------IPDVIARFKRMQGYNVLFPMAWHITGSPIV---------

------------------------------------------------------------------------

-------------------------GIAVNGF---------G--TL--EGQAMSAS----------------

-------KGNVLNFIA--------------------------------------------------------

-----------FP-VSERPSGGGRG-----------------------------------------------

--------------------------------------------------AHLYHS----------------

-------------GGN-----------------------------------------------YS-------

-------------------------

>19A_064_22AVGA-AMSAS-amsasRV1-04_amsas-rv_H08/1-163

-EHHHHHHAELN-FKAIEEKWQKRWLEAKIFEPNIRDKPKEKKFYITVAFPYLSGHLAVGAARTYTIP----

------------------------------------------------------------------------

------------------------------------------------------------------------

------------------------------------------------------------------------

------------------------------------------------------------------------

------------------------------------------------------------------------

------------------------------------------------------------------------

------------------------------------------------------------------------

-------------------AEIIHEMKEEFEYWYPLDWRCSGKDLIPNHLTFFIFNHVAIFREEHWPKGIAV

NGFGTLEGQAMSASKGNVLNFIDAMKKTA-TDG---------------------------------------

------------------------------------------------------------------------

---------------------------------WPXX-----------------XX----------------

------------------------------------------------------------------------

------------------------------------------------------------------------

------------------------------------------------------------------------

------------------------------------------------------------------------

--------------------------------------------------X---------------------

-------------XXX--------------F---------------------X-------------------

-------------------------

>19A_034_21-AVGA-AMSAS-amsas-rv-14_amsas-rv_B05/1-152

-MHHHHHHAELN-FKAIEEKWQKRWLEAKIFEPNIRDKPKEKKFYITVAFPYLSGHLAVGAARTYTIP----

------------------------------------------------------------------------

------------------------------------------------------------------------

------------------------------------------------------------------------

------------------------------------------------------------------------

------------------------------------------------------------------------

------------------------------------------------------------------------

------------------------------------------------------------------------

-------------------AEIIHEMKEEFEYWYPLDWRCSGKDLIPNHLTFFIFNHVAIFREEHWPKGIAV

NGFGTLEGQAMSASKGNVLNFIDAMKKTA-RM----------------------------------------

------------------------------------------------------------------------

------------------------------------------------------------------------

------------------------------------------------------------------------

------------------------------------------------------------------------

------------------------------------------------------------------------

------------------------------------------------------------------------

------------------------------------------------------------------------

------------------------------W------------------------------A----------

-------------------------

>17A_038_AVGAAMSAS-8_AMSAS-sqn2-rv_F05/1-154

-MHHHHHHAELN-FKAIEEKWQKRWLEAKIFEPNIRDKPKEKKFYITVAFPYLSGHLAVGAARTYTIP----

------------------------------------------------------------------------

------------------------------------------------------------------------

------------------------------------------------------------------------

------------------------------------------------------------------------

------------------------------------------------------------------------

------------------------------------------------------------------------

------------------------------------------------------------------------

-------------------AEIIHEMKEEFEYWYPLDWRCSGKDVIPNHLTFFIFNHVAIFREEHWPKGIAV

NGFGTLEGQAMSASKGNVLNFIDAMKKMA-RRG--------------------G------------------

------------------------------------------------------------------------

------------------------------------------------------------------------

------------------------------------------------------------------------

------------------------------------------------------------------------

------------------------------------------------------------------------

------------------------------------------------------------------------

------------------------------------------------------------------------

------------------------------P-----------------------------------------

----------------------G--

>19A_041_22AVGA-AMSAS-AVGAsqun1-01_AVGAsqun1_A06/1-377

-----------X-FLRRLRKWQKRWLEAKIFEPNIRDKPKEKKFYITVAFPYLSGHLAVGAARTYTIP----

------------------------------------------------------------------------

------------------------------------------------------------------------

------------------------------------AEIIHEMKEEFEYWYPLDWRCSGKDLIP--------

NHLTFFIFNHV-------------------------------------------------------------

------------------------------------------------------------------------

------------------------------------------------------------------------

---------------------AIFREEHWPKGIAVNGFGTLEGQAMS----ASK----------GNVLNFIA

FPYLSGHLAVGAARTYTIPAEIIHEMKEEFEYWYPLDWRCSGKDLIPNHLTFFIFNHVAIFREEHWPKGIAV

NGFGTLEGQAMSASKGNVLNFIDAIEENGADVVRLYIMSLAEHDSDFDWRRKEVGKLRKQIERFYELISQFA

EYEVKGNVELKDIDRWMLHRLNKAIKETTNALEEFRTRTAVQW--AFYSIMNDLRWYLRRTEAAMMKRNA--

----------------------MCCATLRCVGA-------------PD-GAVYPHICEEAVEKL--------

GR----------------------------------------------------------------------

------------------------------------------------------------------------

------------------------------------------------------------------------

--------------------------------RAL----------------------

>19A_055_22AVGA-AMSAS-AVGAsqun1-15_AVGAsqun1_G07/1-389

-----------H-FLSRLKIWQKRWLEAKIFEPNIRDKPKEKKFYITVAFPYLSGHLAVGAARTYTIP----

------------------------------------------------------------------------

------------------------------------------------------------------------

------------------------------------AEIIHEMKEEFEYWYPLDWRCSGKDLIP--------

NHLTFFIFNHV-------------------------------------------------------------

------------------------------------------------------------------------

------------------------------------------------------------------------

---------------------AIFREEHWPKGIAVNGFGTLEGQAMS----ASK----------GNVLNFIA

FPYLSGHLAVGAARTYTIPAEIIHEMKEEFEYWYPLDWRCSGKDLIPNHLTFFIFNHVAIFREEHWPKGIAV

NGFGTLEGQAMSASKGNVLNFIDAIEENGADVVRLYIMSLAEHDSDFDWRRKEVGKLRKQIERFYELISQFA

EYEVKGNVELKDIDRWMLHRLNKAIKETTNALEEFRTRTAVQW--AFYSIMNDLRWYLRRTEAAMMKRNA--

----------------------MCCAPCGMLLG-------------CADGAVTPHICEEAVGKL--------

GRRSFG-------ELX--------------------------------------------------------

-----------------NXXG---------------------------------------------------

------------------------------------------------------------------------

--------------------------XXW----------------------------

>19A_061_22AVGA-AMSAS-amsasRV1-01_amsas-rv_E08/1-232

------------------RKMAETLAGSENFEPNIRDKPKEKKFYISVAFPYLSGHLAVGAARTYTIP----

------------------------------------------------------------------------

------------------------------------------------------------------------

------------------------------------AEIIHEMKEEFEYWYPLDWRCSGKDLIP--------

NHLTFFIFNHV-------------------------------------------------------------

------------------------------------------------------------------------

------------------------------------------------------------------------

---------------------AIFREEHWPKGIAVNGFGTLEGQAMS----ASK----------GNVLNFIA

FPYLSGHLAVGAARTYTIPAEIIHEMKEEFEYWYPLDWRCSGKDLIPNHLTFFIFNHVAIFREEHWPKGIAV

NGFGTLEGQAMSASKGNVLNFIDAMKKT-TRMF---------------------------------------

------------------------------------------------------------------------

------------------------------------------------------------------------

------------------------------------------------------------------------

------------------------------------------------------------------------

------------------------------------------------------------------------

--------------------------------------PP-----------------

>LeuAC-Chimr_006/1-240

-L---------K-FKAIEEKWQKRWLEAKIFEPNTRDKPKEKKFYISVAFPYLSGHMAAGAARTYTIP----

------------------------------------------------------------------------

------------------------------------------------------------------------

------------------------------------AEIIHEMKEEFEYWYPLDWRCSGKDLIP--------

NHLTFFIFNHV-------------------------------------------------------------

------------------------------------------------------------------------

------------------------------------------------------------------------

---------------------AIFREEHWPKGIAVNGFGTLEGQAMS----ASK----------GNVLNFIA

FPYLSGHLAVGAARTYTIPAEIIHEMKEEFEYWYPLDWRCSGKDLIPNHLTFFIFNHVAIFREEHWPKGIAV

NGFGTLEGQAMSASKGNVLNFIDAMKKIGARLV---------------------------------------

------------------------------------------------------------------------

------------------------------------------------------------------------

------------------------------------------------------------------------

------------------------------------------------------------------------

------------------------------------------------------------------------

---------------------------------G"----------------------

>19A_047_22AVGA-AMSAS-AVGAsqun1-07_AVGAsqun1_G06/1-135

-----------F-FKR-LKKWQKRWLEAKIFEPNIRDKPKEKKFYITVAFPYLSGHLAVGAARTYTIPDVIA

RFKRMQGYNVLFPMAWHITGSPIV------------------------------------------------

------------------------------------------------------------------------

------------------------------------------------------------------------

------------------------------------------------------------------------

------------------------------------------------------------------------

------------------------------------------------------------------------

------------------------------------------------------------------------

--------------------------------------------------------------------GIAV

NGFGTLEGQAMSASKGNVLNFY-CVSVSE--RP---------------------------------------

------------------------------------------SGGGRG------------------------

---------------------AHLYH---------SG-G---------------NY----------------

------------------------------------------------------------------------

------------------------------------------------------------------------

------------------------------------------------------------------------

--------------------------------------------------S---------------------

------------------------------------------------------------------------

------------------------------------------------------------------------

------------------------A

>19A_015_21-AVGA-AMSAS-AVGAsqun1-15_AVGAsqun1_G02/1-140

-----------TFFKA-IEKWQKRWLEAKIFEPNIRDKPKEKKFYITVAFPYLSGHLAVGAARTYTIPDVIA

RFKRMQGYNVLFPMAWHITGSPIV------------------------------------------------

------------------------------------------------------------------------

------------------------------------------------------------------------

------------------------------------------------------------------------

------------------------------------------------------------------------

------------------------------------------------------------------------

------------------------------------------------------------------------

--------------------------------------------------------------------GIAV

NGFGTLEGQAMSASKGNVLNFI-AFPYLSG-HL---------------------------------------

-----------------------------------------AVGA---------------------------

---------------------ARTYT--------IPVCLA--------------PWKA--------------

------------------------------------------------------------------------

------------------------------------------------------------------------

------------------------------------------------------------------------

--------------------------------------------------R---------------------

------------------------------------------------------------------------

------------------------------------------------------------------------

------------------------R

>19A_067_22AVGA-AMSAS-amsasRV1-07_amsas-rv_C09/1-145

-HHHHHHHAELN-FKAIEEKWQKRWLEAKIFEPNIRDKPKEKKFYITVAFPYLSGHLAVGAARTYTIPDVIA

RFKRMQGYNVLFPMAWHITGSPIV------------------------------------------------

------------------------------------------------------------------------

------------------------------------------------------------------------

------------------------------------------------------------------------

------------------------------------------------------------------------

------------------------------------------------------------------------

------------------------------------------------------------------------

--------------------------------------------------------------------GIAV

NGFGTLEGQAMSASKGNVLNFY-CVSVFE--RP---------------------------------------

------------------------------------------SGGGRG------------------------

---------------------AHLYH---------SG-G---------------NY----------------

------------------------------------------------------------------------

------------------------------------------------------------------------

------------------------------------------------------------------------

------------------------------------------------------------------------

------------------------------------------------------------------------

------------------------------------------------------------------------

------------------------S

>19A_007_21-AVGA-AMSAS-AVGAsqun1-07_AVGAsqun1_G01/1-373

-----------P-FKPI-EKWQKRWLEAKIFEPNIRDKPKEKKFYITVAFPYLSGHLAVGAARTYTIPDVIA

RFKRMQGYNVLFPMAWHITGSPIV------------------------------------------------

------------------------------------------------------------------------

------------------------------------------------------------------------

------------------------------------------------------------------------

------------------------------------------------------------------------

------------------------------------------------------------------------

------------------------------------------------------------------------

--------------------------------------------------------------------GIAV

NGFGTLEGQAMSASKGNVLNFI-AFPYLSG-HL---------------------------------------

-----------------------------------------AVGA---------------------------

---------------------ARTYT--------IPDVIARFKRMQGYNVLFPMAWHITGSPIVGIAERIKN

RDPKTIWIYRDVYKVPEEILWTFEDPINIVKYFMKAAKETFIRAGFSV-----------DWSREFYTTSLFP

PFS--KFIEWQF--------WKLKEKGYIVKGAHRVRWDPVVG--TPLGDHDLMEGE---------------

--------DVPILDYIIIKFELRENG-----------------------------EVIYL-PAAT-------

------------------------------------------------------------------------

-LRPETVYGV--------------------------------------------------------------

TNMWVNPNATYVKAKVR--------------RKDKEETWIVSKEAAYKLSFQIAKFEVIEEFKGEKLIG---

-------------KYVRNP-XAAMK

>19A_016_21-AVGA-AMSAS-AVGAsqun1-16_AVGAsqun1_H02/1-116

-------------------DMAERWLEAKIFEPNIRDKPKEKKFYITVAFPYLSGHLAVGAARTYTIPDVIA

RFKRMQGYNVLFPMAWHITGSPIV------------------------------------------------

------------------------------------------------------------------------

------------------------------------------------------------------------

------------------------------------------------------------------------

------------------------------------------------------------------------

------------------------------------------------------------------------

------------------------------------------------------------------------

--------------------------------------------------------------------GIAV

NGFGTLEGQAMSASKATCTLLTLEGQAMSA-S----------------------------------------

------------------------------------------------------------------------

--------------------------------------------------------KA--------------

------------------------------------------------------------------------

------------------------------------------------------------------------

------------------------------------------------------------------------

--------------------------------------------------T---------------------

------------------------------------------------------------------------

------------------------------------------------------------------------

------------------------C

>19A_005_21-AVGA-AMSAS-AVGAsqun1-05_AVGAsqun1_E01/1-106

-----------N-FLRRLKKWQKRWLEAKIFEPNIRDKPKEKKFYITVAFPYLSGHLAVGAARTYTIPDVIA

RFKRMQGYNVLFPMAWHITGSPIV------------------------------------------------

------------------------------------------------------------------------

------------------------------------------------------------------------

------------------------------------------------------------------------

------------------------------------------------------------------------

------------------------------------------------------------------------

------------------------------------------------------------------------

--------------------------------------------------------------------GIAV

NGFGTLEGQAMSASKAT-------------------------------------------------------

------------------------------------------------------------------------

------------------------------------------------------------------------

------------------------------------------------------------------------

------------------------------------------------------------------------

------------------------------------------------------------------------

------------------------------------------------------------------------

------------------------------------------------------------------------

------------------------------------------------------------------------

------------------------C

>19A_009_21-AVGA-AMSAS-AVGAsqun1-09_AVGAsqun1_A02/1-105

-----------X-FKR-LRKWQKRWLEAKIFEPNIRDKPKEKKFYITVAFPYLSGHLAVGAARTYTIPDVIA

RFKRMQGYNVLFPMAWHITGSPIV------------------------------------------------

------------------------------------------------------------------------

------------------------------------------------------------------------

------------------------------------------------------------------------

------------------------------------------------------------------------

------------------------------------------------------------------------

------------------------------------------------------------------------

--------------------------------------------------------------------GIAV

NGFGTLEGQAMSASKAT-------------------------------------------------------

------------------------------------------------------------------------

------------------------------------------------------------------------

------------------------------------------------------------------------

------------------------------------------------------------------------

------------------------------------------------------------------------

------------------------------------------------------------------------

------------------------------------------------------------------------

------------------------------------------------------------------------

------------------------C

>Leu_prozym_004/1-66

-----------N-FLRRLRKWQKRWLEAKIFEPNIRDKPKEKKFYITVAFPYLSGHLAVGAARTYTIP----

------------------------ALAPWK-AR---------------------------------------

------------------------------------------------------------------------

------------------------------------------------------------------------

------------------------------------------------------------------------

------------------------------------------------------------------------

------------------------------------------------------------------------

------------------------------------------------------------------------

------------------------------------------------------------------------

------------------------------------------------------------------------

------------------------------------------------------------------------

------------------------------------------------------------------------

------------------------------------------------------------------------

------------------------------------------------------------------------

------------------------------------------------------------------------

------------------------------------------------------------------------

------------------------------------------------------------------------

-----------------------------------------------------------R------------

---------------------------------------------------"-

>Leu_prozym_005/1-62

----------------RLRKWQKRWLEAKIFEPNIRDKPKEKKFYITVAFPYLSGHLAVGAARTYTIP----

------------------------ALAPWK-AR---------------------------------------

------------------------------------------------------------------------

------------------------------------------------------------------------

------------------------------------------------------------------------

------------------------------------------------------------------------

------------------------------------------------------------------------

------------------------------------------------------------------------

------------------------------------------------------------------------

------------------------------------------------------------------------

------------------------------------------------------------------------

------------------------------------------------------------------------

------------------------------------------------------------------------

------------------------------------------------------------------------

------------------------------------------------------------------------

------------------------------------------------------------------------

------------------------------------------------------------------------

-----------------------------------------------------------R------------

---------------------------------------------------"-

>Leu_prozym_007/1-59

------------------EKWQKRWLEAKIFEPNIRDKPKEKKFYITVAFPYLSGHLAVGAARTYTIP----

------------------------ALAPWK-AR---------------------------------------

------------------------------------------------------------------------

------------------------------------------------------------------------

------------------------------------------------------------------------

------------------------------------------------------------------------

------------------------------------------------------------------------

------------------------------------------------------------------------

------------------------------------------------------------------------

------------------------------------------------------------------------

------------------------------------------------------------------------

------------------------------------------------------------------------

------------------------------------------------------------------------

------------------------------------------------------------------------

------------------------------------------------------------------------

------------------------------------------------------------------------

------------------------------------------------------------------------

------------------------------------------------------------------------

---------------------------------------------------R-

>Leu_prozym_009/1-65

-----------F-FKAI-EEMAERWLEAKIFEPNIRDKPKEKKFYITVAFPYLSGHLAVGAARTYTIP----

------------------------ALAPWK-AR---------------------------------------

------------------------------------------------------------------------

------------------------------------------------------------------------

------------------------------------------------------------------------

------------------------------------------------------------------------

------------------------------------------------------------------------

------------------------------------------------------------------------

------------------------------------------------------------------------

------------------------------------------------------------------------

------------------------------------------------------------------------

------------------------------------------------------------------------

------------------------------------------------------------------------

------------------------------------------------------------------------

------------------------------------------------------------------------

------------------------------------------------------------------------

------------------------------------------------------------------------

-----------------------------------------------------------R------------

---------------------------------------------------"-

>Leu_prozym_010/1-76

-EHHHHHHAELN-FKAIEEKWQKRWLEAKIFEPNIRDKPKEKKFYITVAFPYLSGHLAVGAARTYTIP----

------------------------ALAPWK-AR---------------------------------------

------------------------------------------------------------------------

------------------------------------------------------------------------

------------------------------------------------------------------------

------------------------------------------------------------------------

------------------------------------------------------------------------

------------------------------------------------------------------------

------------------------------------------------------------------------

------------------------------------------------------------------------

------------------------------------------------------------------------

------------------------------------------------------------------------

------------------------------------------------------------------------

------------------------------------------------------------------------

------------------------------------------------------------------------

------------------------------------------------------------------------

------------------------------------------------------------------------

-----------------------------------------------------------R------------

---------------------------------------------------"-

>Leu_prozym_011/1-76

-EHHHHHHAELN-FKAIEEKWQKRWLEAKIFEPNIRDKPKEKKFYITVAFPYLSGHLAVGAARTYTIP----

------------------------ALAPWK-AR---------------------------------------

------------------------------------------------------------------------

------------------------------------------------------------------------

------------------------------------------------------------------------

------------------------------------------------------------------------

------------------------------------------------------------------------

------------------------------------------------------------------------

------------------------------------------------------------------------

------------------------------------------------------------------------

------------------------------------------------------------------------

------------------------------------------------------------------------

------------------------------------------------------------------------

------------------------------------------------------------------------

------------------------------------------------------------------------

------------------------------------------------------------------------

------------------------------------------------------------------------

-----------------------------------------------------------R------------

---------------------------------------------------"-

>Leu_prozym_012/1-76

-EHHHHHHAELN-FKAIEEKWQKRWLEAKIFEPNIRDKPKEKKFYITVAFPYLSGHLAVGAARTYTIP----

------------------------ALAPWK-AR---------------------------------------

------------------------------------------------------------------------

------------------------------------------------------------------------

------------------------------------------------------------------------

------------------------------------------------------------------------

------------------------------------------------------------------------

------------------------------------------------------------------------

------------------------------------------------------------------------

------------------------------------------------------------------------

------------------------------------------------------------------------

------------------------------------------------------------------------

------------------------------------------------------------------------

------------------------------------------------------------------------

------------------------------------------------------------------------

------------------------------------------------------------------------

------------------------------------------------------------------------

-----------------------------------------------------------R------------

---------------------------------------------------"-

>Leu_prozym_013/1-76

-EHHHHHHAELN-FKAIEEKWQKRWLEAKIFEPNIRDKPKEKKFYITVAFPYLSGHLAVGAARTYTIP----

------------------------ALAPWK-AR---------------------------------------

------------------------------------------------------------------------

------------------------------------------------------------------------

------------------------------------------------------------------------

------------------------------------------------------------------------

------------------------------------------------------------------------

------------------------------------------------------------------------

------------------------------------------------------------------------

------------------------------------------------------------------------

------------------------------------------------------------------------

------------------------------------------------------------------------

------------------------------------------------------------------------

------------------------------------------------------------------------

------------------------------------------------------------------------

------------------------------------------------------------------------

------------------------------------------------------------------------

-----------------------------------------------------------R------------

---------------------------------------------------"-

>Leu_prozym_006/1-293

-----------F-LKRIEEKWQKRWLEAKIFEPNIRDKPKEKKFYITVAFPYLSGHLAVGAARTYTIP----

---------------------------------------EEIIHEMKEQFEYWYPLDWRCR-----------

------------------------------------------------------------------------

------------------------------------------------------------------------

------------------------------------------------------------------------

------------------------------------------------------------------------

------------------------------------------------------------------------

------------------------------------------------------------------------

-----------------------------------------GKYLIPNPLTFFIFNRVAIFREKHWPKGIAV

NGFGPLEGHAKSASKGNVLNFI--------------------------------------------------

--------------------------------AFPYLSGPLAGGA---------------------------

---------------------ARTYT--------IPAEIIREMK-KKFEYWYPLEWRCRGKYLIPN------

--PLTFFIFNRVAIFREK------------------------------------------------------

-------------------HWPK---GIAENGF---------G--TL--ERHAKSAS---------------

--------KGNVLFHMA-------------------------------------------------------

-------------WHITGSPS----VGIAERIK-HRYPNTIWIYRDVYKVREDILCTFEY------------

---------------------------------PINIVIYYMSAARDTYTRA--------------------

--------------GFR--------------V----------------------------------------

-----------------------"-

>17A_005_AVGA-5_AVGA-sqn1_E01.ab1/1-281 5'3'frame3

---------------SRLRKWQKRWLEAKIFEPNIRDKPKEKKFYITVAFPYLSGHLAVGAARTYTIP----

---------------------------------------AEIIHEMKEEFEYWYPLDWRCS-----------

------------------------------------------------------------------------

------------------------------------------------------------------------

------------------------------------------------------------------------

------------------------------------------------------------------------

------------------------------------------------------------------------

------------------------------------------------------------------------

-----------------------------------------GKDLIPNHLTFFIFNHVAIFREEHWPKGIAV

NGFGTLEGQAMSASKGNVLNFI-DAIEEKW-QKR--------------------WL------EAKIF-----

----EPNIRDKPKEKKFYITV-----------AFPYLSGHLAVGA---------------------------

---------------------ARTYT--------IPDVIARFKRMQGYNVLFPMAWHITGSPIV--------

------------------------------------------------------------------------

--------------------------GIAE------------------------------------------

------------------------------------------------------------------------

------------------------------RIK-NRDPKTIWIYRDVYKVPEEILWTFED------------

---------------------------------PINIVKYFMKAAKETFIRA--------------------

--------------GFSVDWSREFYTTRPVSA----------------------------------------

-----------------------V-

>3'5'/1-209 Frame 2_1+2

-------------------------------TEHSPINRKKKKFYITVAFPYLSGHLAVGAARTYTIP----

---------------------------------------AEII-----------------------------

------------------------------------------------------------------------

------------------------------------------------------------------------

------------------------------------------------------------------------

------------------------------------------------------------------------

------------------------------------------------------------------------

------------------------------------------------------------------------

-----------------------HEMKEEFEYWYPLDWRCSGKDLIPNHLTFFIFNHVAIFREEHWPKGIAV

NGFGTLEGQAMSASKGNVLNFI-AFPY---------------------------------------------

--------------------------------------GHLAVGA---------------------------

---------------------ARTYT--------IPDVIARFKRMQGYNVLFPMAWHITGSPIV--------

------------------------------------------------------------------------

--------------------------GIAVNGF---------G--TL--EGQAMSAS---------------

--------KGNVLNFIA-------------------------------------------------------

------------FP-VFERPSGGGRG----------------------------------------------

---------------------------------------------------AHLYHS---------------

--------------GGN-------------------------------------------------------

-----------------------YS-

>19A_074_22AVGA-AMSAS-amsasRV1-14_amsas-rv_B10/1-110

---------------------------------------------TLLRFPYLSGHLAVGAARTYTIP----

---------------------------------------AEII-----------------------------

------------------------------------------------------------------------

------------------------------------------------------------------------

------------------------------------------------------------------------

------------------------------------------------------------------------

------------------------------------------------------------------------

------------------------------------------------------------------------

-----------------------HEMKEEFEYWYPLDWRCSGKDLIPNHLTFFIFNHVAIFREEHWPKGIAV

NGFGTLEGQAMSASKGNVLNFI-DAMKKTA-RMW--------------------------------------

------------------------------------------------------------------------

------------------------------------------------------------------------

------------------------------------------------------------------------

------------------------------------------------------------------------

------------------------------------------------------------------------

------------------------------------------------------------------------

------------------------------------------------------------------------

-------------------------------A----------------------------------------

------------------------P

>19A_054_22AVGA-AMSAS-AVGAsqun1-14_AVGAsqun1_F07/1-151

---------------------------------------------TLLRFPYLSGHLAVGAARTYTIP----

---------------------------------------AEII-----------------------------

------------------------------------------------------------------------

------------------------------------------------------------------------

------------------------------------------------------------------------

------------------------------------------------------------------------

------------------------------------------------------------------------

------------------------------------------------------------------------

-----------------------HEMKEEFEYWYPLDWRCSGKDLIPNHLTFFIFNHVAIFREEHWPKGIAV

NGFGTLESQAMSASKGNVLNFI-DAIEENG-ADVVRLYIMSLAEHDSDFDWRRKEVGKLRKQIERFYELISQ

FADM--------------------------------------------------------------------

------------------------------------------------------------------------

------------------------------------------------------------------------

------------------------------------------------------------------------

------------------------------------------------------------------------

------------------------------------------------------------------------

------------------------------------------------------------------------

------------------------------------------------------------------------

------------------------K

C. Varieties of genomic structure. The following figures illustrate the variety of urzyme, Goldilocks and protozyme sequences appearing in sequenced plasmid samples. These include single open-reading frame, (Supplemental figure 1, 5); tandem-configurations, (Supplemental Figure 2, 3, 7); and multiple embedding in multiple frames in the same plasmid, (Supplemental Figure 3, 4, 6, 7). Interestingly, Goldilocks and protozyme sequences appeared only at lower temperature 4^o^ C.

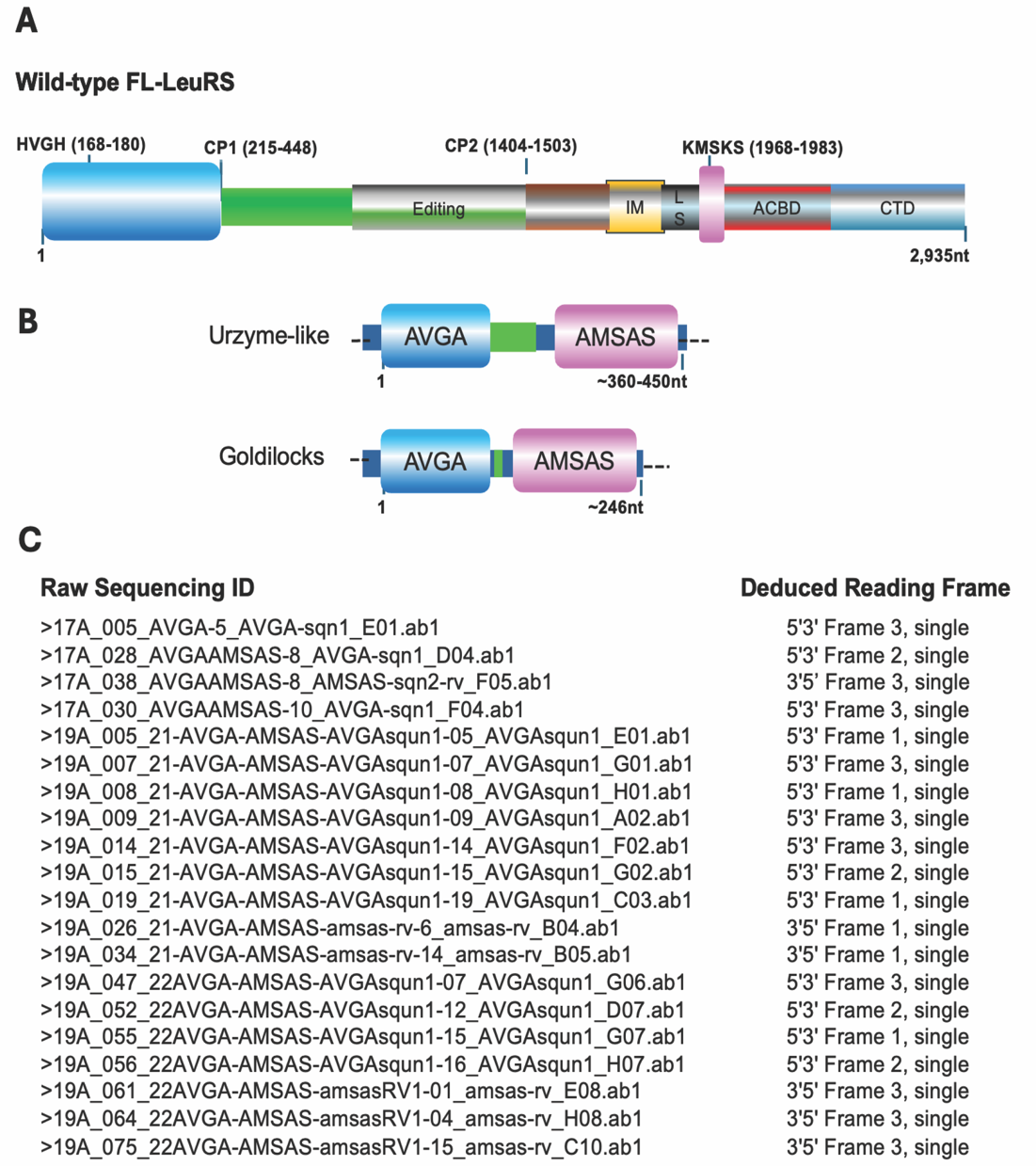

**Supplemental Figure S1.** Schematic diagram depicting genomic annotation of the sequenced insert from plasmids collected from two sets of small experiments. **A.** A cartoon describing Full-length Leucyl-tRNA synthetase (FL-LeuRS) labelled with domains and motifs. **B.** Symbolic scheme representation of urzyme-like sequence and goldilocks with AVGA and AMSAS mutations of the conventional catalytic signatures HVGH and KMSKS. **C.** The IDs of individual raw sequencing samples with single urzyme-like and goldilocks ORFs, corresponding to 6 possible ORFs. Raw sequencing identification (ID) on left; deduced open reading frame on right.

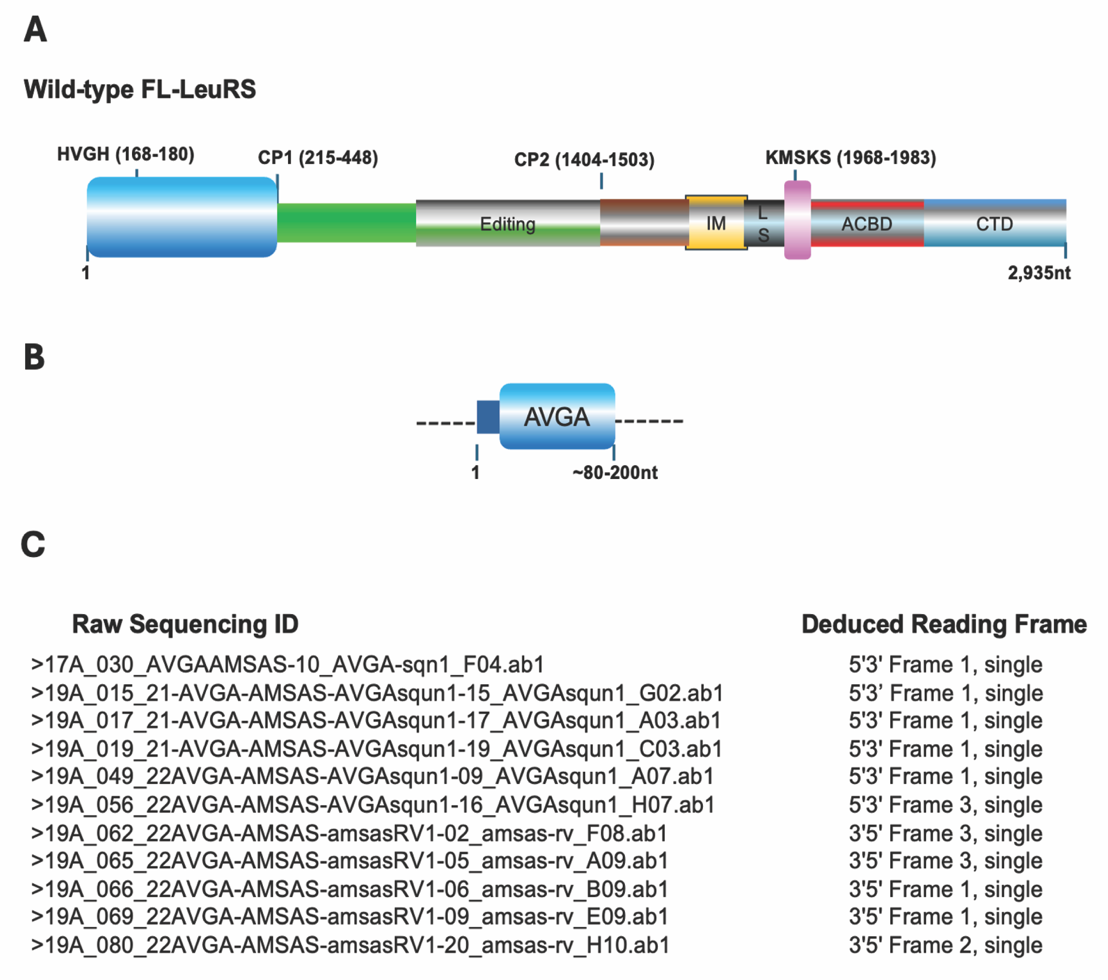

**Supplemental Figure S2.** Schematic diagram depicting genomic features of truncated protozyme sequences at the same sequenced insert region of same plasmid in a sequenced population. **A.** Full-length Leucyl-tRNA synthetase (FL-LeuRS) labelled with domains and motifs. **B.** Symbolic representation of genomic features of the truncated protozyme sequences as indicated in combinations of multi-frames and tandem-array in each sequenced insert. **C.** The sequenced plasmids with IDs of individual raw sequencing sample, and genomic features of each individual truncated protozyme are summarized.

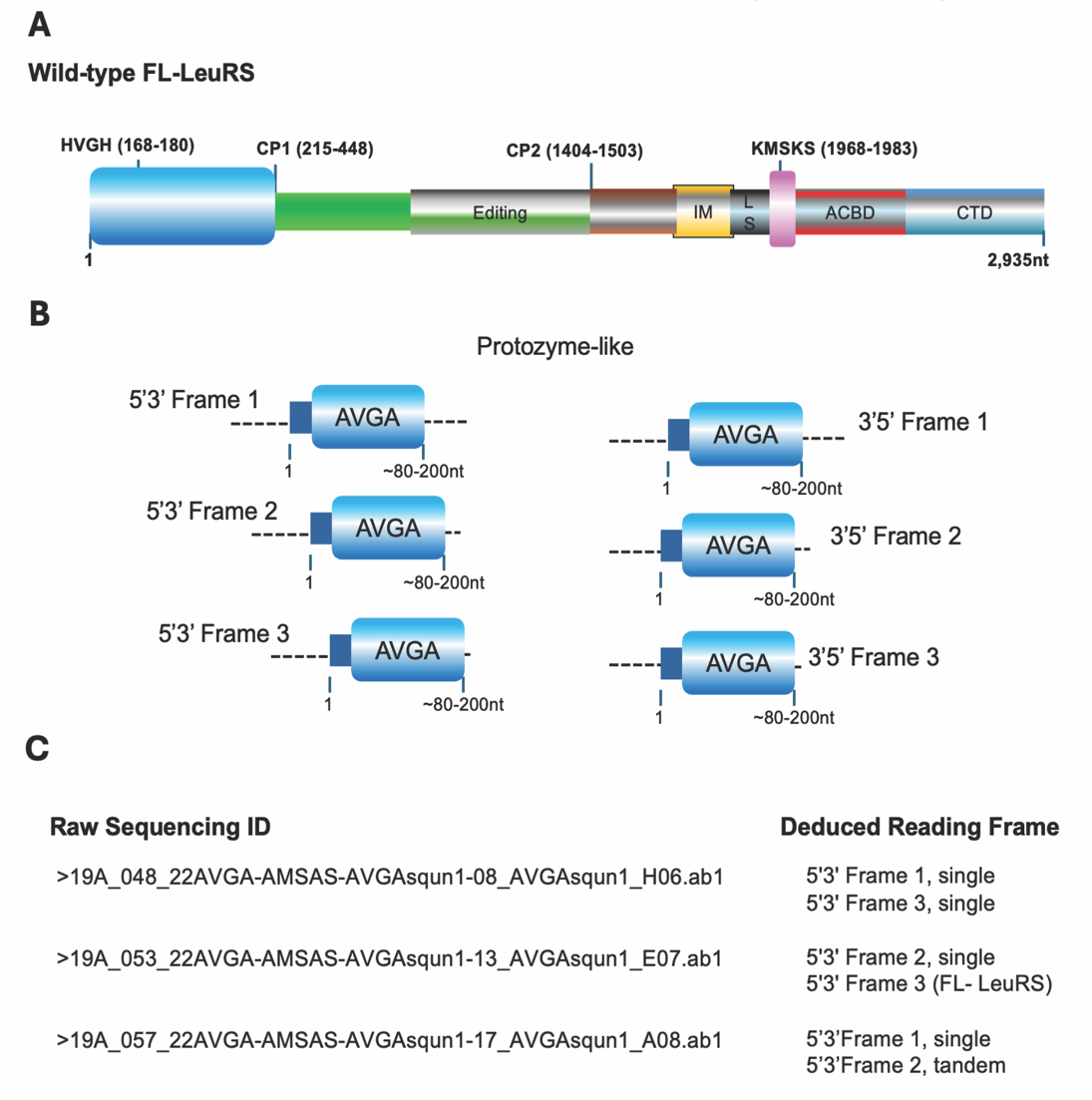

**Supplemental Figure S3.** A schematic display of single ORF genomic features of truncated protozyme sequences in the list of sequenced inserts at tested pool of plasmids. **A.** Full-length Leucyl-tRNA synthetase (FL-LeuRS) labelled with domains and motifs. **B.** Symbolic representation of the organization of protozyme-like sequences in single ORFs as indicated. **C.** IDs of individual raw sequencing data containing the genomic features indicated in **B**.

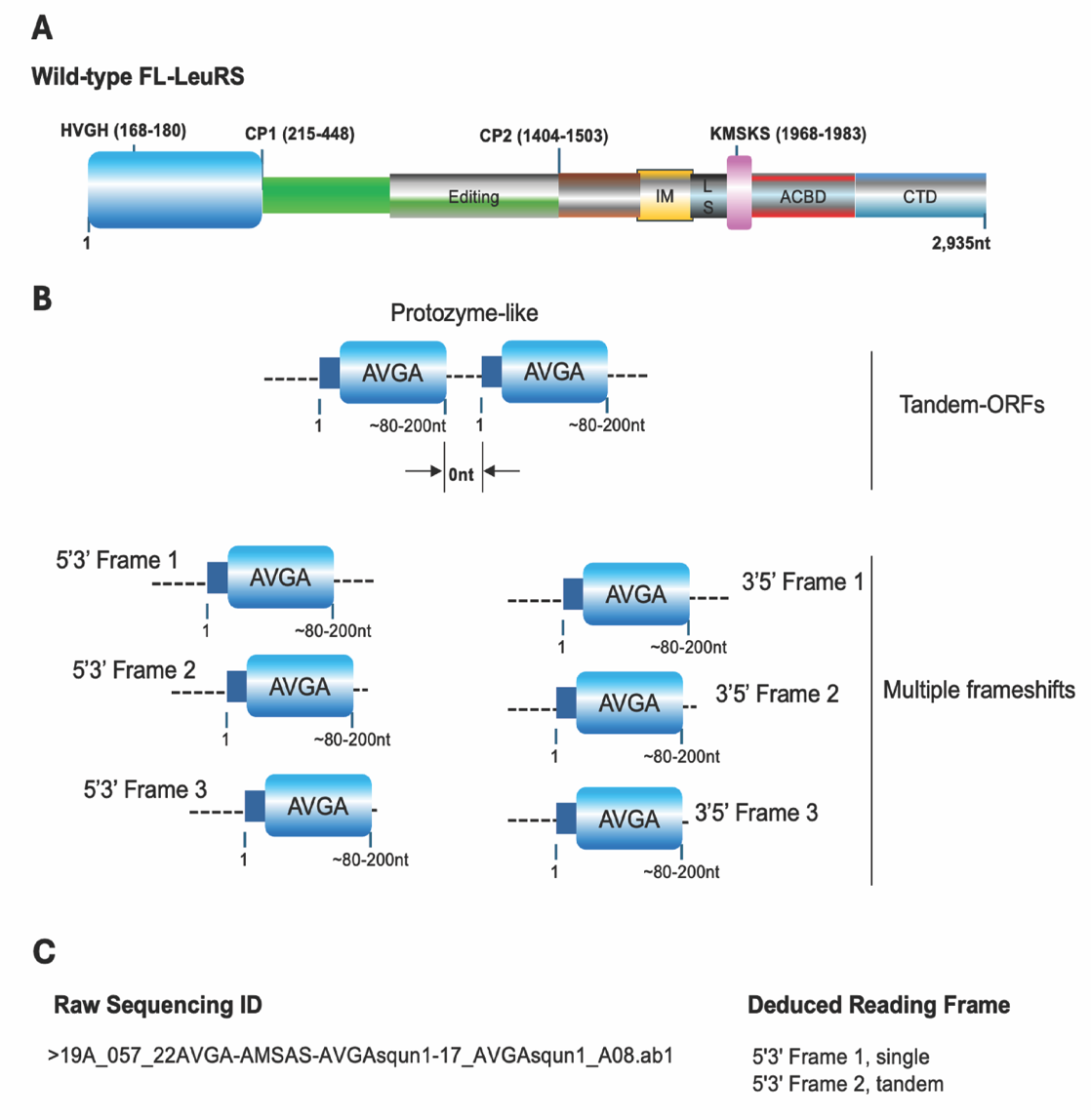

**Supplemental Figure S4.** Schematic diagram depicting of genomic features with multiple-frames of truncated protozyme sequences at sequenced insert regions of plasmid population. **A.** Full-length Leucyl-tRNA synthetase (FL-LeuRS) labelled with domains and motifs. **B.** Symbolic representation of genomic features of protozyme sequences as indicated with combinations of 6 reading frames in each sequenced insert. **C.** The sequenced plasmids with IDs of individual raw sequencing sample, and genomic features of each individual truncated protozyme.

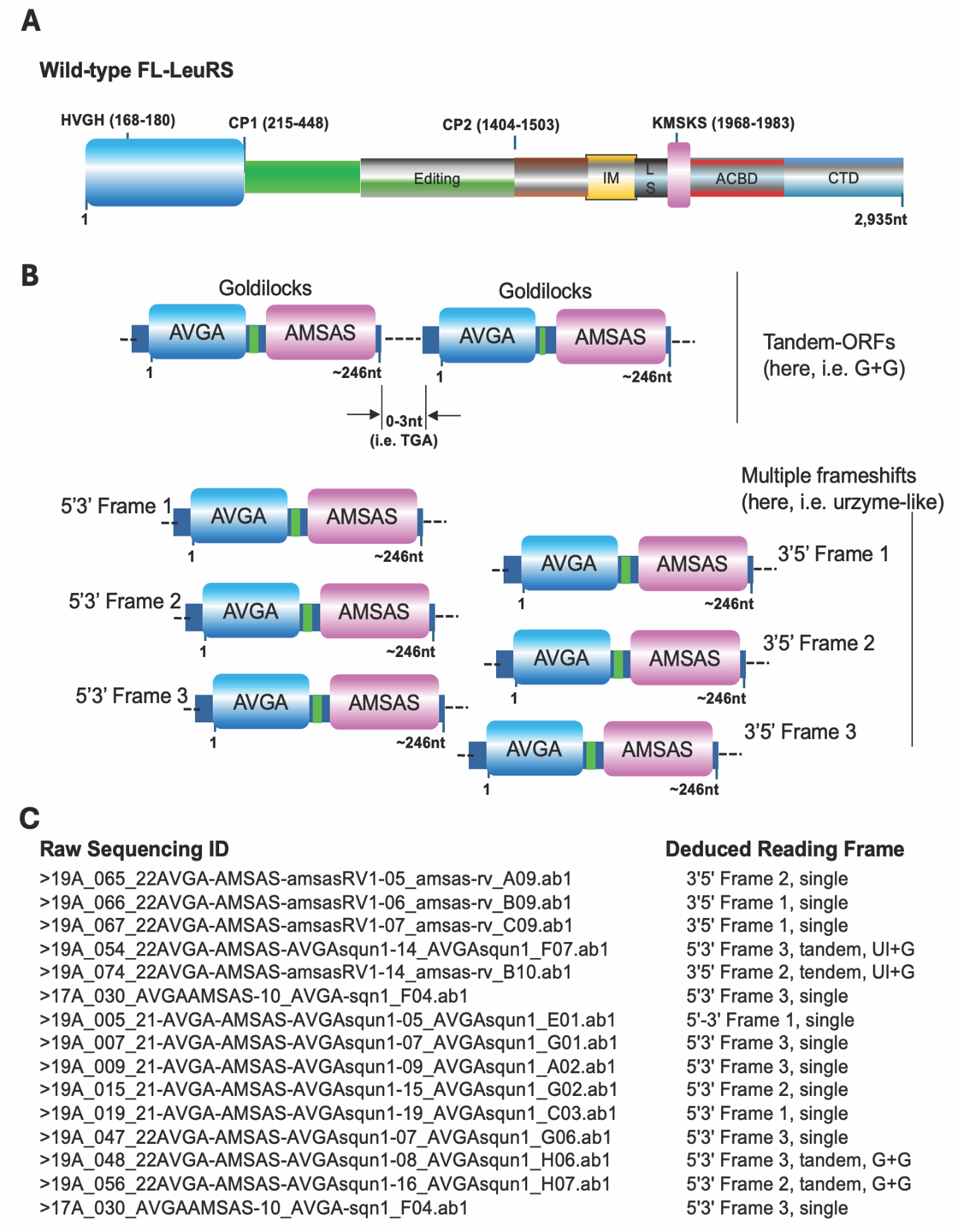

**Supplemental Figure S5.** Schematic diagram depicting Goldilocks sequences from sequenced insert regions of plasmid population with multiple Goldilocks ORFs in the same sequenced insert. **A.** Full-length Leucyl-tRNA synthetase (FL-LeuRS) labelled with domains and motifs. **B.** Symbolic representation of genomic features of Goldilocks sequences as indicated, showing combinations of multi-frames and ***6 reading frames***. **C.** The sequenced plasmids with IDs of individual raw sequencing sample, and genomic features of each individual Goldilocks element are summarized.

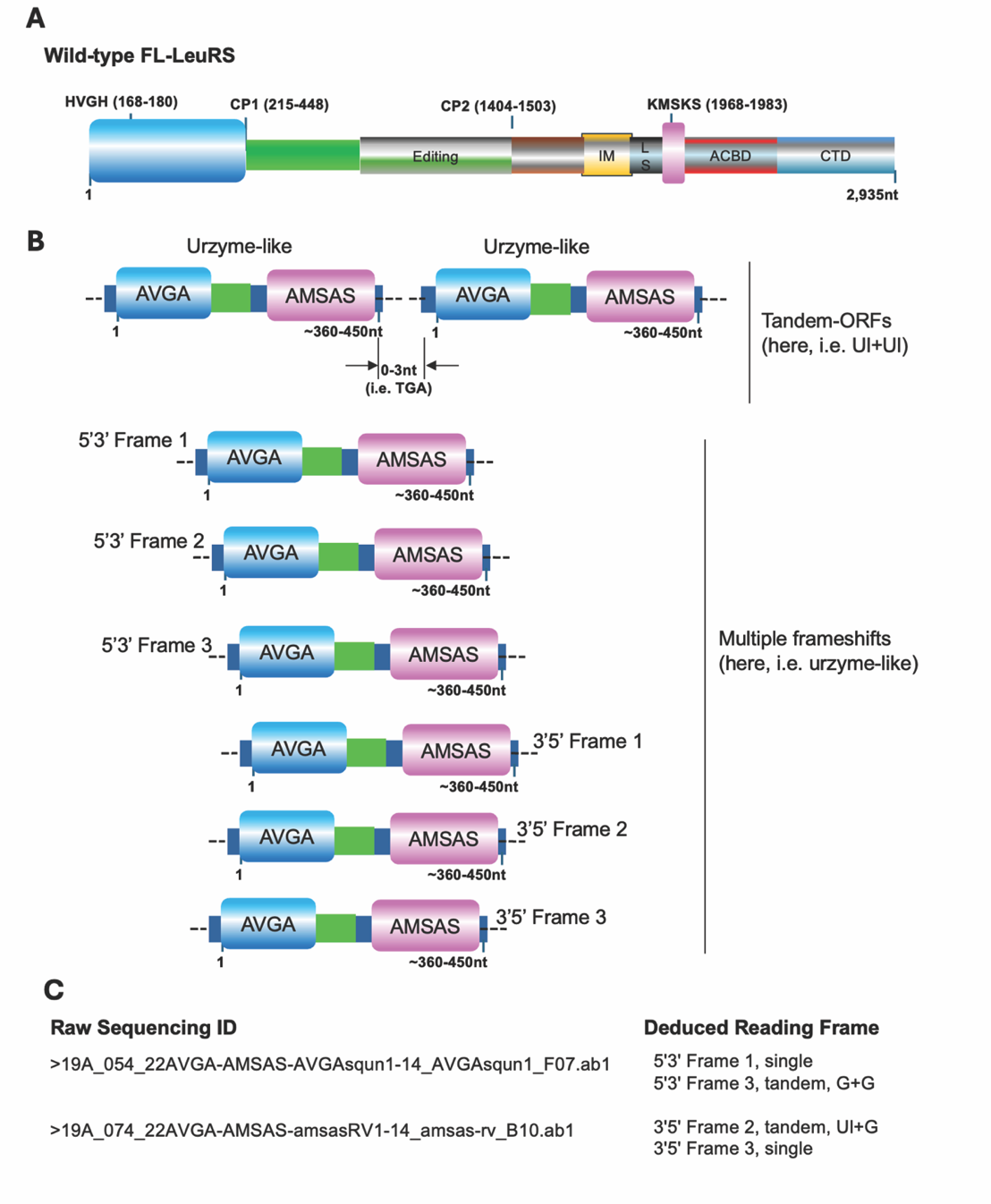

**Supplemental Figure S6.** Schematic representation of tandemly arranged urzyme-like elements in sequenced inserts at plasmids from two sets of experiments in small scales. **A.** Diagram indicates Full-length Leucyl-tRNA synthetase (FL-LeuRS) labelled with domains and motifs. **B.** Symbolic representation of the organization of urzyme-like ORFs in tandem arrays with a gap as indicated. **C.** A list of the sequenced plasmids with IDs of individual raw sequencing sample that include tandem-ORFs embedded, corresponding to 6 possible ORFs as indicated. Abbreviations: Ul, urzyme-like; G, goldilocks.

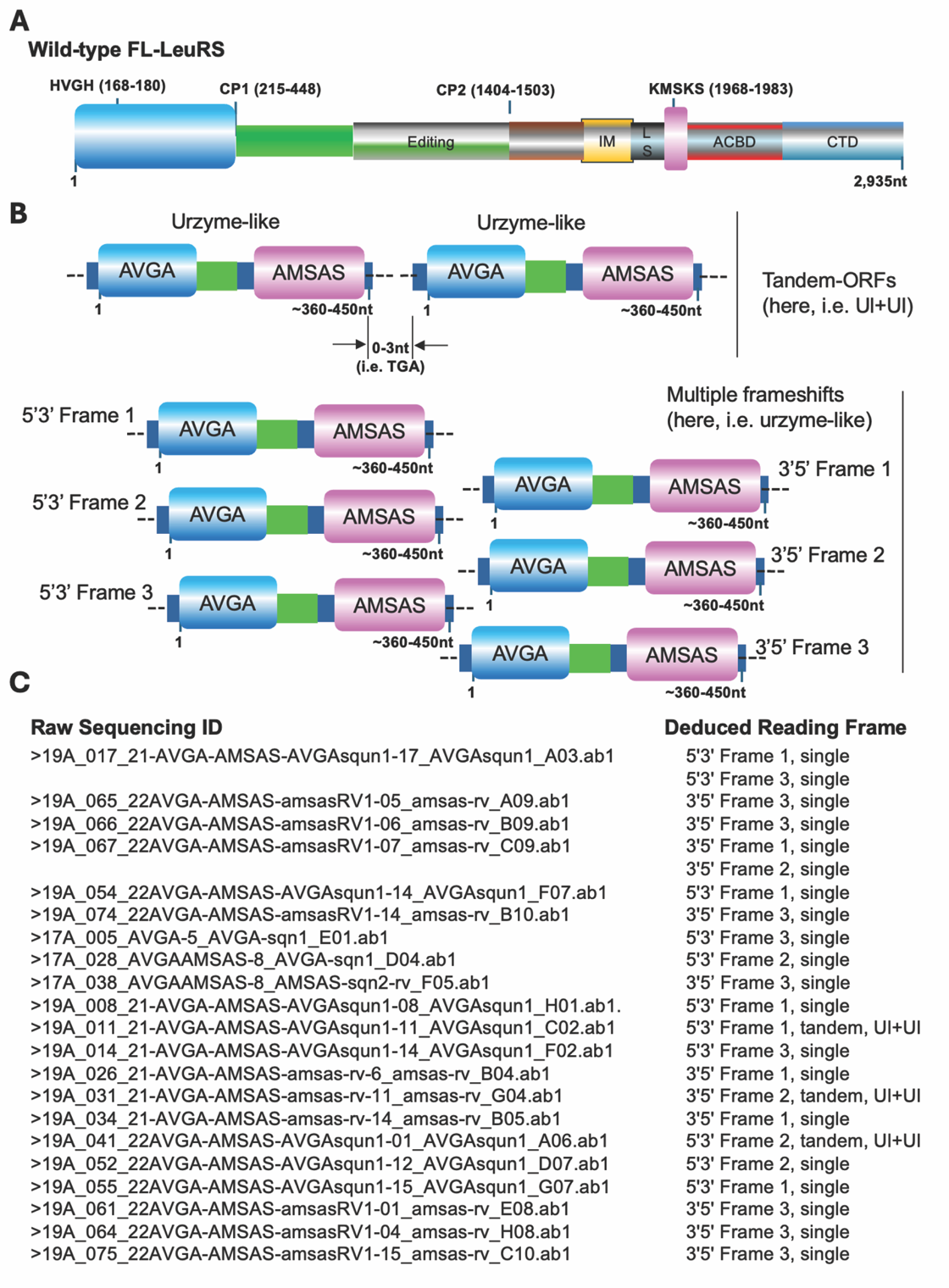

**Supplemental Figure S7.** A schematic display of genomic features of urzyme-like sequences found in either multiple-frame shift ORFs or tandemy-repeated elements in sequenced inserts. **A.** Full-length Leucyl-tRNA synthetase (FL-LeuRS) labelled with domains and motifs. **B.** Symbolic representation of organization of urzyme-like and goldilocks sequences in both of multiple-frameshifts and tandem arrays with a gap as indicated. **C.** List of individual raw sequencing samples with IDs on left, genomic features (both single and tandem-ORFs) were listed on right, corresponding to 6 possible ORFs as indicated in **B** and **C**.

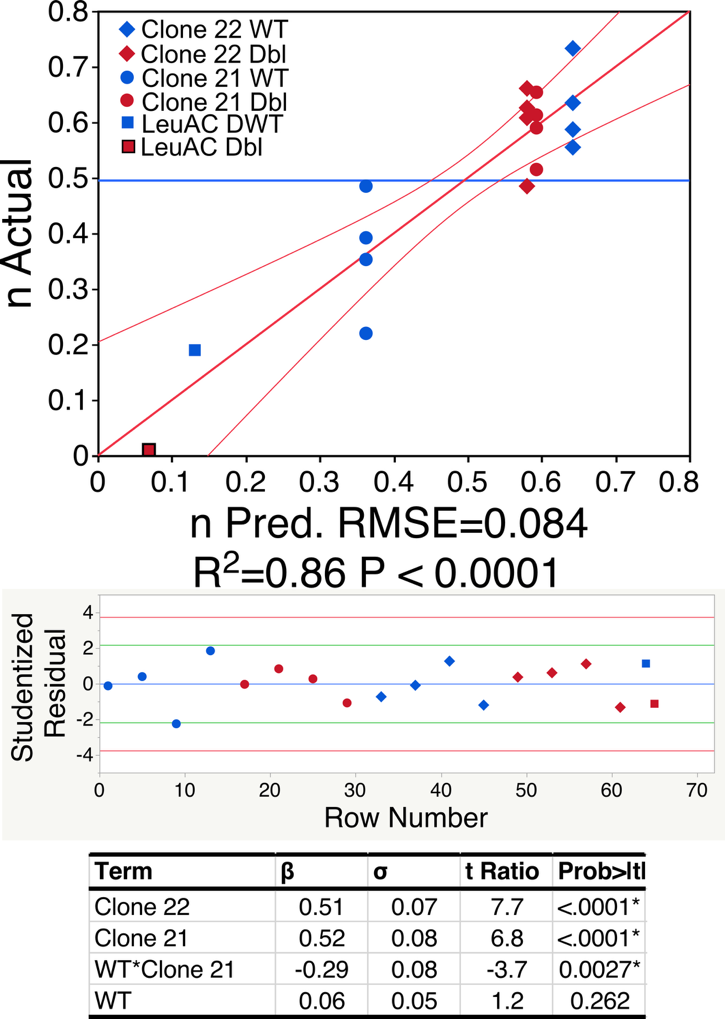

Supplementary Figure S8. Multiple regression model for the n-value derived from burst sizes. The table of coefficients omits the intercept, which is ill-determined (P>0.3) and so is sorted in order of decreasing statistical significance. Note that the effects of WT active-site residues is the opposite for Clone 21 (red circles have higher n-values than blue circles) and Clone 22 (blue diamonds have higher n-values that the red diamonds). That is consistent with the negative sign of the WT*Glone21 interaction term.

**Table S1 Genomic Features of deletion plasmid coding regions**

| **Sequence ID** | **Reading Frame** | **Identification** | **Temp** | **ORF** | **Urzyme-like** | **Goldilocks** | **Protozyme-like** | **Full Length** | **Tandem** |
| --- | --- | --- | --- | --- | --- | --- | --- | --- | --- |
| >17A_005_AVGA-5_AVGA-sqn1_E01.ab1 | 5'3' Frame 3 | **Urzyme-like** | 37 | 2 | 1 | 0 | 0 | 0 | 0 |
| >17A_028_AVGAAMSAS-8_AVGA-sqn1_D04.ab1 | 5'3' Frame1 | Urzyme-like | 37 | 1 | 1 | 0 | 0 | 0 | 0 |
| >17A_038_AVGAAMSAS-8_AMSAS-sqn2-rv_F05.ab1 | 5'3' Frame 1 | Urzyme-like | 37 | -3 | 1 | 0 | 0 | 0 | 0 |
| >17A_030_AVGAAMSAS-10_AVGA-sqn1_F04.ab1 | 5'3' Frame 1 | Goldilocks | 37 | 2 | 0 | 1 | 0 | 0 | 0 |
| >19A_005_21-AVGA-AMSAS-AVGAsqun1-05_AVGAsqun1_E01.ab1 | 5'3' Frame 3 | Goldilocks | 4 | 0 | 0 | 1 | 0 | 0 | 0 |
| >19A_007_21-AVGA-AMSAS-AVGAsqun1-07_AVGAsqun1_G01.ab1 | 5'3' Frame 1 | Goldilocks | 4 | 2 | 0 | 1 | 0 | 0 | 0 |
| >19A_008_21-AVGA-AMSAS-AVGAsqun1-08_AVGAsqun1_H01.ab1. | 5'3' Frame 2 | Urzyme-like | 4 | 0 | 1 | 0 | 0 | 0 | 0 |
| >19A_009_21-AVGA-AMSAS-AVGAsqun1-09_AVGAsqun1_A02.ab1 | 5'3' Frame 3 | Goldilocks | 4 | 1 | 0 | 1 | 0 | 0 | 0 |
| >19A_011_21-AVGA-AMSAS-AVGAsqun1-11_AVGAsqun1_C02.ab1 | 5'3' Frame 3 | Urzyme-like | 4 | 2 | 1 | 0 | 0 | 0 | 1 |
| >19A_014_21-AVGA-AMSAS-AVGAsqun1-14_AVGAsqun1_F02.ab1 | 5'3' Frame 1 | Urzyme-like | 4 | 0 | 1 | 0 | 0 | 0 | 0 |
| >19A_015_21-AVGA-AMSAS-AVGAsqun1-15_AVGAsqun1_G02.ab1 | 5'3' Frame 2 | Goldilocks | 4 | 1 | 0 | 1 | 0 | 0 | 0 |
| >19A_019_21-AVGA-AMSAS-AVGAsqun1-19_AVGAsqun1_C03.ab1 | 3'5' Frame 3 | Goldilocks | 4 | -3 | 0 | 1 | 0 | 0 | 0 |
| >19A_026_21-AVGA-AMSAS-amsas-rv-6_amsas-rv_B04.ab1 | 3'5' Frame 3 | Urzyme-like | 4 | -1 | 1 | 0 | 0 | 0 | 0 |
| >19A_031_21-AVGA-AMSAS-amsas-rv-11_amsas-rv_G04.ab1 | 3'5' Frame 1 | Urzyme-like | 4 | -1 | 1 | 0 | 0 | 0 | 1 |
| >19A_034_21-AVGA-AMSAS-amsas-rv-14_amsas-rv_B05.ab1 | 3'5' Frame 1 | Urzyme-like | 4 | -1 | 1 | 0 | 0 | 0 | 0 |
| >19A_041_22AVGA-AMSAS-AVGAsqun1-01_AVGAsqun1_A06.ab1 | 3'5' Frame 2 | Urzyme-like | 4 | -2 | 1 | 0 | 0 | 0 | 1 |
| >19A_047_22AVGA-AMSAS-AVGAsqun1-07_AVGAsqun1_G06.ab1 | 5'3' Frame 1 | Goldilocks | 4 | 0 | 0 | 1 | 0 | 0 | 0 |
| >19A_048_22AVGA-AMSAS-AVGAsqun1-08_AVGAsqun1_H06.ab1 | 5'3' Frame 3 | Goldilocks | 4 | 2 | 0 | 1 | 0 | 0 | 0 |
| >19A_052_22AVGA-AMSAS-AVGAsqun1-12_AVGAsqun1_D07.ab1 | 5'3' Frame 2 | Urzyme-like | 4 | 1 | 1 | 0 | 0 | 0 | 0 |
| >19A_055_22AVGA-AMSAS-AVGAsqun1-15_AVGAsqun1_G07.ab1 | 5'3' Frame 1 | Urzyme-like | 4 | 0 | 1 | 0 | 0 | 0 | 0 |
| >19A_056_22AVGA-AMSAS-AVGAsqun1-16_AVGAsqun1_H07.ab1 | 5'3' Frame 2 | Goldilocks | 4 | 1 | 0 | 1 | 0 | 0 | 0 |
| >19A_061_22AVGA-AMSAS-amsasRV1-01_amsas-rv_E08.ab1 | 3'5' Frame 3 | Urzyme-like | 4 | -3 | 1 | 0 | 0 | 0 | 0 |
| >19A_064_22AVGA-AMSAS-amsasRV1-04_amsas-rv_H08.ab1 | 3'5' Frame 3 | Urzyme-like | 4 | -3 | 1 | 0 | 0 | 0 | 0 |
| >19A_075_22AVGA-AMSAS-amsasRV1-15_amsas-rv_C10.ab1 | 3'5' Frame 3 | Urzyme-like | 4 | -3 | 1 | 0 | 0 | 0 | 0 |
| >17A_030_AVGAAMSAS-10_AVGA-sqn1_F04.ab1 | 5'3' Frame 3 | Goldilocks | 37 | 2 | 0 | 1 | 0 | 0 | 0 |
| >19A_065_22AVGA-AMSAS-amsasRV1-05_amsas-rv_A09.ab1 | 3'5' Frame 2 | Goldilocks | 4 | -2 | 0 | 1 | 0 | 0 | 0 |
| >19A_066_22AVGA-AMSAS-amsasRV1-06_amsas-rv_B09.ab1 | 3'5' Frame 1 | Goldilocks | 4 | -1 | 0 | 1 | 0 | 0 | 0 |
| >19A_067_22AVGA-AMSAS-amsasRV1-07_amsas-rv_C09.ab1 | 3'5' Frame 1 | Goldilocks | 4 | -1 | 0 | 1 | 0 | 0 | 0 |
| >19A_054_22AVGA-AMSAS-AVGAsqun1-14_AVGAsqun1_F07.ab1 | 5'3' Frame 3 | Goldilocks | 4 | 2 | 0 | 1 | 0 | 0 | 0 |
| >19A_074_22AVGA-AMSAS-amsasRV1-14_amsas-rv_B10.ab1 | 3'5' Frame 2 | Goldilocks | 4 | -2 | 0 | 1 | 0 | 0 | 0 |
| >17A_030_AVGAAMSAS-10_AVGA-sqn1_F04.ab1 | 5'3' Frame 3 | Goldilocks | 37 | 2 | 0 | 1 | 0 | 0 | 0 |
| >19A_005_21-AVGA-AMSAS-AVGAsqun1-05_AVGAsqun1_E01.ab1 | 5'3' Frame 1 | Goldilocks | 4 | 0 | 0 | 1 | 0 | 0 | 0 |
| >19A_007_21-AVGA-AMSAS-AVGAsqun1-07_AVGAsqun1_G01.ab1 | 5'3' Frame 3 | Goldilocks | 4 | 2 | 0 | 1 | 0 | 0 | 0 |
| >19A_009_21-AVGA-AMSAS-AVGAsqun1-09_AVGAsqun1_A02.ab1 | 5'3' Frame 3 | Goldilocks | 4 | 2 | 0 | 1 | 0 | 0 | 0 |
| >19A_015_21-AVGA-AMSAS-AVGAsqun1-15_AVGAsqun1_G02.ab1 | 5'3' Frame2 | Goldilocks | 4 | 1 | 0 | 1 | 0 | 0 | 0 |
| >19A_019_21-AVGA-AMSAS-AVGAsqun1-19_AVGAsqun1_C03.ab1 | 5'3' Frame 1 | Goldilocks | 4 | 0 | 0 | 1 | 0 | 0 | 0 |
| >19A_047_22AVGA-AMSAS-AVGAsqun1-07_AVGAsqun1_G06.ab1 | 5'3' Frame 3 | Goldilocks | 4 | 2 | 0 | 1 | 0 | 0 | 0 |
| >19A_048_22AVGA-AMSAS-AVGAsqun1-08_AVGAsqun1_H06.ab1 | 5'3' Frame 3 | Goldilocks | 4 | 2 | 0 | 1 | 0 | 0 | 0 |
| >19A_056_22AVGA-AMSAS-AVGAsqun1-16_AVGAsqun1_H07.ab1 | 5'3' Frame 2 | Goldilocks | 37 | 1 | 0 | 1 | 0 | 0 | 0 |
| >17A_030_AVGAAMSAS-10_AVGA-sqn1_F04.ab1 | 5'3' Frame 3 | Goldilocks | 4 | 2 | 0 | 1 | 0 | 0 | 0 |
| >19A_069_22AVGA-AMSAS-amsasRV1-09_amsas-rv_E09.ab1 | 3'5' Frame 1 | Protozyme-like | 37 | -1 | 0 | 0 | 1 | 0 | 0 |
| >19A_080_22AVGA-AMSAS-amsasRV1-20_amsas-rv_H10.ab1 | 3'5' Frame 2 | Protozyme-like | 37 | -2 | 0 | 0 | 1 | 0 | 0 |
| >17A_030_AVGAAMSAS-10_AVGA-sqn1_F04.ab1 | 5'3' Frame 1 | Protozyme-like | 4 | 0 | 0 | 0 | 1 | 0 | 0 |
| >19A_017_21-AVGA-AMSAS-AVGAsqun1-17_AVGAsqun1_A03.ab1 | 5'3' Frame 1 | Urzyme-like | 37 | 0 | 1 | 0 | 0 | 0 | 0 |
| >19A_017_21-AVGA-AMSAS-AVGAsqun1-17_AVGAsqun1_A03.ab1 | 5'3' Frame 3 | Urzyme-like | 37 | 2 | 1 | 0 | 0 | 0 | 0 |
| >19A_065_22AVGA-AMSAS-amsasRV1-05_amsas-rv_A09.ab1 | 3'5' Frame 2 | Goldilocks | 37 | -2 | 0 | 1 | 0 | 0 | 0 |
| >19A_065_22AVGA-AMSAS-amsasRV1-05_amsas-rv_A09.ab1 | 3'5' Frame 3 | Urzyme-like | 37 | -3 | 1 | 0 | 0 | 0 | 0 |
| >19A_066_22AVGA-AMSAS-amsasRV1-06_amsas-rv_B09.ab1 | 3'5' Frame 1 | Goldilocks | 37 | -1 | 0 | 1 | 0 | 0 | 0 |
| >19A_066_22AVGA-AMSAS-amsasRV1-06_amsas-rv_B09.ab1 | 3'5' Frame 3 | Urzyme-like | 37 | -3 | 1 | 0 | 0 | 0 | 0 |
| >19A_067_22AVGA-AMSAS-amsasRV1-07_amsas-rv_C09.ab1 | 3'5' Frame 1 | Urzyme-like | 37 | -1 | 1 | 0 | 0 | 0 | 0 |
| >19A_067_22AVGA-AMSAS-amsasRV1-07_amsas-rv_C09.ab1 | 3'5' Frame 2 | Urzyme-like | 37 | -2 | 1 | 0 | 0 | 0 | 0 |
| >19A_054_22AVGA-AMSAS-AVGAsqun1-14_AVGAsqun1_F07.ab1 | 5'3' Frame 1 | Urzyme-like | 37 | 0 | 1 | 0 | 0 | 0 | 0 |
| >19A_054_22AVGA-AMSAS-AVGAsqun1-14_AVGAsqun1_F07.ab1 | 5'3' Frame 3 | Goldilocks | 37 | 2 | 0 | 1 | 0 | 0 | 0 |
| >19A_074_22AVGA-AMSAS-amsasRV1-14_amsas-rv_B10.ab1 | 3'5' Frame 2 | Goldilocks | 37 | -2 | 0 | 1 | 0 | 0 | 0 |
| >19A_074_22AVGA-AMSAS-amsasRV1-14_amsas-rv_B10.ab1 | 3'5' Frame 3 | Urzyme-like | 37 | -3 | 1 | 0 | 0 | 0 | 0 |
| >19A_048_22AVGA-AMSAS-AVGAsqun1-08_AVGAsqun1_H06.ab1 | 5'3'Frame1 | Protozyme-like | 37 | 0 | 0 | 0 | 1 | 0 | 0 |
| >19A_048_22AVGA-AMSAS-AVGAsqun1-08_AVGAsqun1_H06.ab1 | 5'3' Frame1 | Protozyme-like | 37 | 0 | 0 | 0 | 1 | 0 | 0 |
| >19A_015_21-AVGA-AMSAS-AVGAsqun1-15_AVGAsqun1_G02.ab1 | 5'3' Frame 1 | Protozyme-like | 37 | 0 | 0 | 0 | 1 | 0 | 0 |
| >19A_017_21-AVGA-AMSAS-AVGAsqun1-17_AVGAsqun1_A03.ab1 | 5'3' Frame 1 | Protozyme-like | 37 | 0 | 0 | 0 | 1 | 0 | 0 |
| >19A_019_21-AVGA-AMSAS-AVGAsqun1-19_AVGAsqun1_C03.ab1 | 5'3' Frame 3 | Protozyme-like | 37 | 2 | 0 | 0 | 1 | 0 | 0 |
| >19A_049_22AVGA-AMSAS-AVGAsqun1-09_AVGAsqun1_A07.ab1 | 5'3' Frame 1 | Protozyme-like | 37 | 0 | 0 | 0 | 1 | 1 | 0 |
| >19A_053_22AVGA-AMSAS-AVGAsqun1-13_AVGAsqun1_E07.ab1 | 5'3' Frame 2 | Protozyme-like | 37 | 1 | 0 | 0 | 1 | 1 | 0 |
| >19A_053_22AVGA-AMSAS-AVGAsqun1-13_AVGAsqun1_E07.ab1 | 5'3' Frame 3 | Protozyme-like | 37 | 2 | 0 | 0 | 1 | 0 | 0 |
| >19A_056_22AVGA-AMSAS-AVGAsqun1-16_AVGAsqun1_H07.ab1 | 5'3' Frame 3 | Protozyme-like | 37 | 2 | 0 | 0 | 1 | 0 | 0 |
| >19A_057_22AVGA-AMSAS-AVGAsqun1-17_AVGAsqun1_A08.ab1 | 5'3' Frame 1 | Protozyme-like | 37 | 0 | 0 | 0 | 1 | 0 | 0 |
| >19A_057_22AVGA-AMSAS-AVGAsqun1-17_AVGAsqun1_A08.ab1 | 5'3' Frame 2 | Protozyme-like | 37 | 1 | 0 | 0 | 1 | 0 | 0 |
| >19A_062_22AVGA-AMSAS-amsasRV1-02_amsas-rv_F08.ab1 | 3'5' Frame 3 | Protozyme-like | 37 | -3 | 0 | 0 | 1 | 0 | 0 |
| >19A_065_22AVGA-AMSAS-amsasRV1-05_amsas-rv_A09.ab1 | 3'5' Frame 3 | Protozyme-like | 37 | -3 | 0 | 0 | 1 | 0 | 0 |
| >19A_066_22AVGA-AMSAS-amsasRV1-06_amsas-rv_B09.ab1 | 3'5' Frame 1 | Protozyme-like | 37 | -1 | 0 | 0 | 1 | 0 | 0 |
| >19A_069_22AVGA-AMSAS-amsasRV1-09_amsas-rv_E09.ab1 | 3'5' Frame 1 | Protozyme-like | 37 | -1 | 0 | 0 | 1 | 0 | 0 |
| >19A_080_22AVGA-AMSAS-amsasRV1-20_amsas-rv_H10.ab1 | 3'5' Frame 2 | Protozyme-like | 37 | -2 | 0 | 0 | 1 | 0 | 0 |
| >17A_030_AVGAAMSAS-10_AVGA-sqn1_F04.ab1 | 5'3' Frame 1 | Protozyme-like | 4 | 0 | 0 | 0 | 1 | 0 | 0 |

**E. Design matrices for regression analysis**.

Table S2. Design matrix for regression analysis of active-site titration parameters. Independent parameters in the last three columns are: WT, the presence of full LeuAC; 2ndXvr, the presence of the second crossover of the Rossmann dinucleotide binding fold; Protoz, the presence of the full length of the protozyme.

| Variant | ΔG^‡^k_chem_ | ΔG^‡^k_3_ | ΔG^‡^k_chem_/k_3_ | WT | 2^ND^ XVR | Protoz |
| --- | --- | --- | --- | --- | --- | --- |
| WT 21–All | 3.3 | 6.03 | -2.73 | 1 | 0 | 1 |
| WT 21–All | 3.44 | 6.66 | -3.22 | 1 | 0 | 1 |
| WT 21–All | 2.85 | 6.72 | -3.87 | 1 | 0 | 1 |
| WT 21–All | 2.65 | 6.21 | -3.56 | 1 | 0 | 1 |
| WT 21–1 | 3.7 | 5.87 | -2.17 | 1 | 0 | 1 |
| WT 21–1 | 3.91 | 7.11 | -3.2 | 1 | 0 | 1 |
| WT 21–1 | 3.11 | 6.5 | -3.39 | 1 | 0 | 1 |
| WT 21–1 | 3.22 | 6.1 | -2.88 | 1 | 0 | 1 |
| WT 21-2 | 3.32 | 5.89 | -2.57 | 1 | 0 | 1 |
| WT 21-2 | 3.13 | 6.29 | -3.16 | 1 | 0 | 1 |
| WT 21-2 | 3.63 | 6.75 | -3.12 | 1 | 0 | 1 |
| WT 21-2 | 3.78 | 6.39 | -2.61 | 1 | 0 | 1 |
| WT 21-3 | 3.14 | 5.65 | -2.51 | 1 | 0 | 1 |
| WT 21-3 | 3.34 | 5.39 | -2.05 | 1 | 0 | 1 |
| WT 21-3 | 3.02 | 5.7 | -2.68 | 1 | 0 | 1 |
| WT 21-3 | 2.66 | 5.84 | -3.18 | 1 | 0 | 1 |
| DBL 21-All | 3.26 | 5.82 | -2.56 | 0 | 1 | 1 |
| DBL 21-All | 3.45 | 6.58 | -3.13 | 0 | 1 | 1 |
| DBL 21-All | 2.82 | 6.51 | -3.69 | 0 | 1 | 1 |
| DBL 21-All | 2.93 | 6.11 | -3.18 | 0 | 1 | 1 |
| DBL 21-1 | 3.46 | 5.88 | -2.42 | 0 | 1 | 1 |
| DBL 21-1 | 3.61 | 7.44 | -3.83 | 0 | 1 | 1 |
| DBL 21-1 | 3.2 | 6.34 | -3.14 | 0 | 1 | 1 |
| DBL 21-1 | 3.3 | 5.96 | -2.66 | 0 | 1 | 1 |
| DBL 21-2 | 3.13 | 5.96 | -2.83 | 0 | 1 | 1 |
| DBL 21-2 | 3.17 | 6.34 | -3.17 | 0 | 1 | 1 |
| DBL 21-2 | 2.9 | 6.82 | -3.92 | 0 | 1 | 1 |
| DBL 21-2 | 3.27 | 6.45 | -3.18 | 0 | 1 | 1 |
| DBL 21-3 | 3.16 | 5.74 | -2.58 | 0 | 1 | 1 |
| DBL 21-3 | 3.13 | 5.25 | -2.12 | 0 | 1 | 1 |
| WT 22–All | 3.52 | 6.56 | -3.06 | 1 | 0 | 0 |
| WT 22–All | 3.54 | 6.58 | -2.58 | 1 | 0 | 0 |
| WT 22–1 | 3.99 | 7.35 | -3.04 | 1 | 0 | 0 |
| WT 22–1 | 4.01 | 7.05 | -3.04 | 1 | 0 | 0 |
| WT 22-2 | 3.27 | 6.66 | -4.66 | 1 | 0 | 0 |
| WT 22-2 | 3.12 | 6.73 | -4.36 | 1 | 0 | 0 |
| WT 22-3 | 3.57 | 7.03 | -3.36 | 1 | 0 | 0 |
| WT 22-3 | 3.82 | 7.84 | -3.04 | 1 | 0 | 0 |
| DBL 22–All | 3.26 | 6.05 | -4.36 | 0 | 1 | 0 |
| DBL 22–All | 3.44 | 6.34 | -3.39 | 0 | 1 | 0 |
| DBL 22-1 | 3.37 | 6.04 | -3.39 | 0 | 1 | 0 |
| DBL 22-1 | 3.29 | 6.26 | -3.61 | 0 | 1 | 0 |
| DBL 22-2 | 3.13 | 6.3 | -4.26 | 0 | 1 | 0 |
| DBL 22-2 | 3.2 | 6.73 | -3.9 | 0 | 1 | 0 |
| DBL 22-3 | 3.14 | 5.65 | -3.46 | 0 | 1 | 0 |
| DBL 22-3 | 3.34 | 5.39 | -4.02 | 0 | 1 | 0 |
| LeuAC_WT_Ile | 2.98 | 6.24 | -4.5 | 1 | 1 | 1 |
| LeuAC_DOUBLE_Ile | 3.43 | 6.27 | -3.99 | 0 | 1 | 1 |
| LeuAC_WT_LEU | 3.26 | 5.96 | -2.79 | 1 | 1 | 1 |
| LeuAC_DOUBLE_LEU | 2.49 | 5.95 | -2.9 | 0 | 1 | 1 |
| LeuAC_WT_Ile_tRNA | 3.67 | 6.58 | -4.43 | 1 | 1 | 1 |
| LeuAC_DOUBLE_Ile_tRNA | 3.63 | 9.2 | -4.01 | 0 | 1 | 1 |
| LeuAC_WT_Ile_MINI | 3.57 | 6.37 | -2.67 | 1 | 1 | 1 |
| LeuAC_DOUBLE_Ile_MINI | 3.04 | 6.17 | -2.97 | 0 | 1 | 1 |

Table S3. Design matrix for the analysis of steady-state kinetics of aminoacylation by Clones 21 and 22 as isolated (AVGA, AMSAS; Dbl) and corresponding constructs with wild-type catalytic signatures (HVGH, KMSKS; WT). Dependent variables are the free energies and were computed as –0.592*ln(Ksp, kcat, KM). Independent variables are coded as (0,1) according to the presence or absence of the WT catalytic signatures, the second crossover connection of the Rossmann dinucleotide binding fold, and the full ~50-residue protozyme.

| Dataset | K_sp_ | k_cat_ | K_M_ | ΔG^‡^K_sp_ | ΔG G^‡^k_cat_ | ΔG G^‡^K_M_ | WT | 2^nd^ Xvr | Protoz |
| --- | --- | --- | --- | --- | --- | --- | --- | --- | --- |
| Clone21_WT | 2.00E+04 | 3.51E-03 | 1.75E-07 | -5.86E+00 | 3.35E+00 | 9.21E+00 | 1 | 0 | 1 |
| Clone21_WT | 1.46E+04 | 3.66E-03 | 2.51E-07 | -5.67E+00 | 3.32E+00 | 9.00E+00 | 1 | 0 | 1 |
| Clone21_WT | 1.24E+04 | 3.74E-03 | 3.00E-07 | -5.58E+00 | 3.31E+00 | 8.89E+00 | 1 | 0 | 1 |
| Clone21_WT | 6.00E+04 | 3.33E-03 | 5.55E-08 | -6.51E+00 | 3.38E+00 | 9.89E+00 | 1 | 0 | 1 |
| Clone21_DBL | 1.54E+04 | 1.82E-03 | 1.18E-07 | -5.71E+00 | 3.73E+00 | 9.44E+00 | 0 | 1 | 1 |
| Clone21_DBL | 7.60E+03 | 1.97E-03 | 2.60E-07 | -5.29E+00 | 3.69E+00 | 8.98E+00 | 0 | 1 | 1 |
| Clone21_DBL | 9.66E+03 | 2.06E-03 | 2.13E-07 | -5.43E+00 | 3.66E+00 | 9.09E+00 | 0 | 1 | 1 |
| Clone21_DBL | 4.44E+04 | 1.71E-03 | 3.86E-08 | -6.34E+00 | 3.77E+00 | 1.01E+01 | 0 | 1 | 1 |
| Clone22_WT | 1.05E+04 | 1.83E-03 | 1.74E-07 | -5.48E+00 | 3.73E+00 | 9.21E+00 | 1 | 0 | 0 |
| Clone22_WT | 6.24E+03 | 1.89E-03 | 3.03E-07 | -5.17E+00 | 3.71E+00 | 8.89E+00 | 1 | 0 | 0 |
| Clone22_WT | 7.65E+03 | 2.00E-03 | 2.62E-07 | -5.29E+00 | 3.68E+00 | 8.97E+00 | 1 | 0 | 0 |
| Clone22_WT | 4.07E+04 | 1.69E-03 | 4.16E-08 | -6.28E+00 | 3.78E+00 | 1.01E+01 | 1 | 0 | 0 |
| Clone22_DBL_Mt | 1.87E+04 | 4.99E-03 | 2.66E-07 | -5.82E+00 | 3.14E+00 | 8.96E+00 | 0 | 1 | 0 |
| Clone22_DBL_Mt | 1.18E+04 | 6.16E-03 | 5.24E-07 | -5.55E+00 | 3.01E+00 | 8.56E+00 | 0 | 1 | 0 |
| Clone22_DBL_Mt | 1.60E+04 | 5.46E-03 | 3.42E-07 | -5.73E+00 | 3.08E+00 | 8.81E+00 | 0 | 1 | 0 |
| Clone22_DBL_Mt | 1.13E+05 | 3.74E-03 | 3.32E-08 | -6.89E+00 | 3.31E+00 | 1.02E+01 | 0 | 1 | 0 |
| LeuAC_Ile_Mini_WT | 7.18E+01 | 1.48E-04 | 2.06E-06 | -2.53E+00 | 5.22E+00 | 7.75E+00 | 1 | 1 | 1 |
| LeuAC_Ile_Mini_WT | 5.93E+01 | 5.92E-04 | 9.99E-06 | -2.42E+00 | 4.40E+00 | 6.82E+00 | 1 | 1 | 1 |
| LeuAC_Ile_Mini_WT | 1.73E+02 | 6.99E-05 | 4.04E-07 | -3.05E+00 | 5.66E+00 | 8.72E+00 | 1 | 1 | 1 |
| LeuAC_Ile_Mini_WT | 7.73E+01 | 9.36E-05 | 1.21E-06 | -2.57E+00 | 5.49E+00 | 8.07E+00 | 1 | 1 | 1 |
| LeuAC_Ile_Mini_DOUBLE | 3.79E+03 | 1.14E-03 | 3.00E-07 | -4.88E+00 | 4.01E+00 | 8.89E+00 | 0 | 1 | 1 |
| LeuAC_Ile_Mini_DOUBLE | 3.04E+03 | 1.37E-03 | 4.51E-07 | -4.75E+00 | 3.90E+00 | 8.65E+00 | 0 | 1 | 1 |
| LeuAC_Ile_Mini_DOUBLE | 5.11E+03 | 1.04E-03 | 2.03E-07 | -5.05E+00 | 4.07E+00 | 9.12E+00 | 0 | 1 | 1 |
| LeuAC_Ile_Mini_DOUBLE | 3.11E+03 | 1.10E-03 | 3.54E-07 | -4.76E+00 | 4.03E+00 | 8.79E+00 | 0 | 1 | 1 |

Table S4 Design matrix for regression analysis of the <AMP/ADP> ratio

| Variant | <AMP/ADP> | Clone 21 | WT |
| --- | --- | --- | --- |
| Clone21(WT) | 0.412 | 1 | 1 |
| Clone21(Dbl) | 0.297 | 1 | 0 |
| Clone22(WT) | 0.285 | 0 | 1 |
| Clone221(Dbl) | 0.212 | 0 | 0 |

Table S5. Design matrix for estimating steady-state ΔG^‡^kcat from single turnover data.

| Variant | ΔG^‡^k_chem_ | ΔG^‡^k_3_ | ΔG^‡^k_cat_ |
| --- | --- | --- | --- |
| 21WT | 3.26 | 6.19 | 3.34 |
| 21DBL | 3.20 | 6.23 | 3.67 |
| 22WT | 3.60 | 6.98 | 3.73 |
| 22DBL | 3.27 | 6.10 | 3.14 |
| LeuAC | 3.26 | 6.59 | 4.60 |

Table S6. Design matrix for regression modeling of Michaelis-Menten parameters.

| Dataset | Ksp | kcat | KM | ΔGKsp | Mean/Stdev | ΔGkcat | Mean/Stdev | ΔGKM | Mean/Stdev | WT | 2^nd^ Xvr | Protoz |
| --- | --- | --- | --- | --- | --- | --- | --- | --- | --- | --- | --- | --- |
| Clone21_WT | 2.00E+04 | 3.51E-03 | 1.75E-07 | -5.86E+00 |  | 3.35E+00 |  | 9.21E+00 |  | 1 | 0 | 1 |
| Clone21_WT | 1.46E+04 | 3.66E-03 | 2.51E-07 | -5.67E+00 |  | 3.32E+00 |  | 9.00E+00 |  | 1 | 0 | 1 |
| Clone21_WT | 1.24E+04 | 3.74E-03 | 3.00E-07 | -5.58E+00 | -5.91 | 3.31E+00 | 3.34 | 8.89E+00 | 9.25 | 1 | 0 | 1 |
| Clone21_WT | 6.00E+04 | 3.33E-03 | 5.55E-08 | -6.51E+00 | ±0.42 | 3.38E+00 | ±0.03 | 9.89E+00 | ±0.45 | 1 | 0 | 1 |
| Clone21_DBL | 1.54E+04 | 1.82E-03 | 1.18E-07 | -5.71E+00 |  | 3.73E+00 |  | 9.44E+00 |  | 0 | 0 | 1 |
| Clone21_DBL | 7.60E+03 | 1.97E-03 | 2.60E-07 | -5.29E+00 |  | 3.69E+00 |  | 8.98E+00 |  | 0 | 0 | 1 |
| Clone21_DBL | 9.66E+03 | 2.06E-03 | 2.13E-07 | -5.43E+00 | -5.69 | 3.66E+00 | 3.71 | 9.09E+00 | 9.40E+00 | 0 | 0 | 1 |
| Clone21_DBL | 4.44E+04 | 1.71E-03 | 3.86E-08 | -6.34E+00 | ±0.46 | 3.77E+00 | ±0.05 | 1.01E+01 | ±0.51 | 0 | 0 | 1 |
| Clone22_WT | 1.05E+04 | 1.83E-03 | 1.74E-07 | -5.48E+00 |  | 3.73E+00 |  | 9.21E+00 |  | 1 | 1 | 0 |
| Clone22_WT | 6.24E+03 | 1.89E-03 | 3.03E-07 | -5.17E+00 |  | 3.71E+00 |  | 8.89E+00 |  | 1 | 1 | 0 |
| Clone22_WT | 7.65E+03 | 2.00E-03 | 2.62E-07 | -5.29E+00 | -5.56 | 3.68E+00 | 3.73 | 8.97E+00 | 9.28 | 1 | 1 | 0 |
| Clone22_WT | 4.07E+04 | 1.69E-03 | 4.16E-08 | -6.28E+00 | ±0.50 | 3.78E+00 | ±0.04 | 1.01E+01 | ±0.54 | 1 | 1 | 0 |
| Clone22_DBL_Mt | 1.87E+04 | 4.99E-03 | 2.66E-07 | -5.82E+00 |  | 3.14E+00 |  | 8.96E+00 |  | 0 | 1 | 0 |
| Clone22_DBL_Mt | 1.18E+04 | 6.16E-03 | 5.24E-07 | -5.55E+00 |  | 3.01E+00 |  | 8.56E+00 |  | 0 | 1 | 0 |
| Clone22_DBL_Mt | 1.60E+04 | 5.46E-03 | 3.42E-07 | -5.73E+00 | -6.00 | 3.08E+00 | 3.14 | 8.81E+00 | 9.13 | 0 | 1 | 0 |
| Clone22_DBL_Mt | 1.13E+05 | 3.74E-03 | 3.32E-08 | -6.89E+00 | ±0.60 | 3.31E+00 | ±0.13 | 1.02E+01 | ±0.73 | 0 | 1 | 0 |
| LeuAC_Ile_Mini_WT | 7.18E+01 | 1.48E-04 | 2.06E-06 | -2.53E+00 |  | 5.22E+00 |  | 7.75E+00 |  | 1 | 1 | 1 |
| LeuAC_Ile_Mini_WT | 5.93E+01 | 5.92E-04 | 9.99E-06 | -2.42E+00 |  | 4.40E+00 |  | 6.82E+00 |  | 1 | 1 | 1 |
| LeuAC_Ile_Mini_WT | 1.73E+02 | 6.99E-05 | 4.04E-07 | -3.05E+00 | -2.64E+00 | 5.66E+00 | 5.19 | 8.72E+00 | 7.84 | 1 | 1 | 1 |
| LeuAC_Ile_Mini_WT | 7.73E+01 | 9.36E-05 | 1.21E-06 | -2.57E+00 | 0.28 | 5.49E+00 | ±0.56 | 8.07E+00 | ±0.79 | 1 | 1 | 1 |
| LeuAC_Ile_Mini_DOUBLE | 3.79E+03 | 1.14E-03 | 3.00E-07 | -4.88E+00 |  | 4.01E+00 |  | 8.89E+00 |  | 0 | 1 | 1 |
| LeuAC_Ile_Mini_DOUBLE | 3.04E+03 | 1.37E-03 | 4.51E-07 | -4.75E+00 |  | 3.90E+00 |  | 8.65E+00 |  | 0 | 1 | 1 |
| LeuAC_Ile_Mini_DOUBLE | 5.11E+03 | 1.04E-03 | 2.03E-07 | -5.05E+00 | -4.86E+00 | 4.07E+00 | 4.00 | 9.12E+00 | 8.86 | 0 | 1 | 1 |
| LeuAC_Ile_Mini_DOUBLE | 3.11E+03 | 1.10E-03 | 3.54E-07 | -4.76E+00 | 0.14 | 4.03E+00 | ±0.07 | 8.79E+00 | ±0.20 | 0 | 1 | 1 |

F. **Estimating steady-state rates from single turnover rate constants**.

To investigate further the evident difficulty of performing steady-state assays of aminoacylation by AARS urzymes, we developed a regression model expressing ΔGk_cat_ in terms of the free energy for the first order rate constant, ΔGk_chem_ and that for turnover, ΔGk_3_. The data themselves are very noisy with respect to this question. However, as we had sufficient multiple estimates for all rate constants to reduce the standard error of the mean value for each variable to ~1%. The design matrix in Supplementary Table S4 shows this design matrix. The regression model is summarized in Supplementary Fig. 9. The table of regression coefficients implies that the proportional contributions to the ΔGk_cat_ value of the first-order and turnover are in the ratio of 2:1.

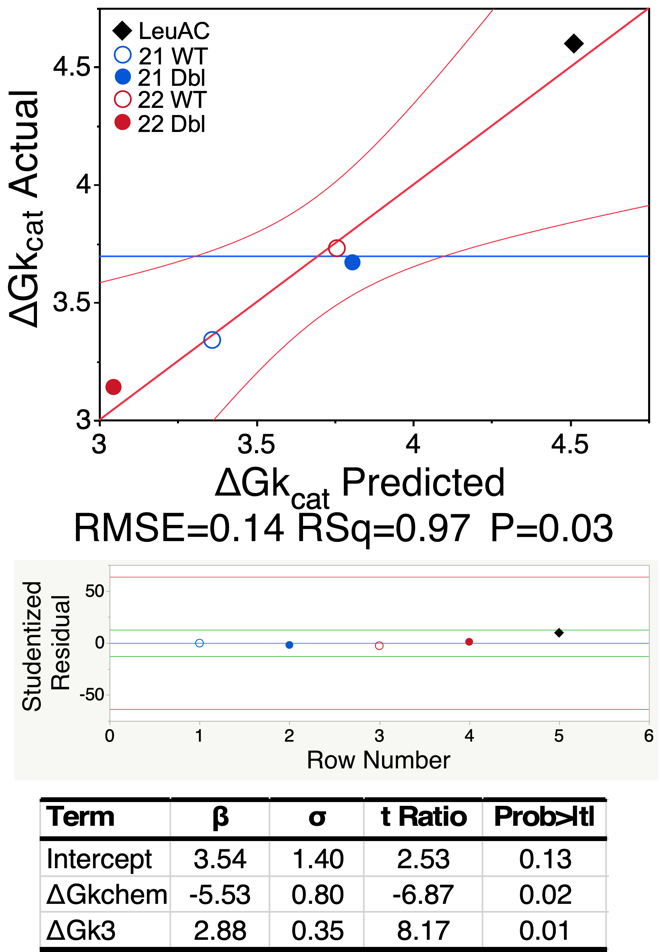

**Supplementary Figure S8.** Regression model for steady state ΔGk_cat_ in terms of the free energies for the two single-turnover rate constants. β and σ are regression coefficients and their standard deviations. The errors are thus < 15%. Their P values are more significant than the 95% level.
